# RNAcompare: Integrating machine learning algorithms to unveil the similarities of phenotypes based on patients’ clinical, multi-omics using Rheumatoid Arthritis and Heart Failure as Case Studies

**DOI:** 10.1101/2025.04.02.646760

**Authors:** Mingcan Tang

## Abstract

**Background:** Gene expression analysis is crucial for understanding the biological mechanisms underlying patient subgroup differences. However, most existing studies focus primarily on transcriptomic data while neglecting the integration of clinical heterogeneity. Although batch correction methods are commonly used, challenges remain when integrating data across different tissues, omics layers, and diseases.

This limitation hampers the ability to connect molecular insights to phenotypic outcomes, thereby restricting clinical translation. Furthermore, the technical complexity of analysing large, heterogeneous datasets poses a barrier for clinicians. To address these challenges, we present RNAcompare, a development of RNAcare, employing machine learning techniques to integrate clinical and multi-omics data seamlessly.

**Results:** RNAcompare overcomes these challenges by providing an interactive, reproducible platform designed to analyse multi-omics data in a clinical context. This tool enables researchers to integrate diverse datasets, conduct exploratory analyses, and identify shared patterns across patient subgroups. The platform facilitates hypothesis generation and produces intuitive visualizations to support further investigation.

As a proof of concept, we applied RNAcompare to connect omics data to pain, fatigue, drug resistance in rheumatoid arthritis (RA) and disease severity to RA and Heart Failure (HF). Our analysis reduced selection bias and managed heterogeneity by identifying key contributors to treatment variability. We discovered shared molecular pathways associated with different treatments. Using SHAP (Shapley Additive Explanations) values, we successfully classified patients into three subgroups based on age, and subsequent analyses confirmed these age-related patterns. Additionally, we uncovered hidden patterns influencing pain and disease severity across different tissues, omics layers, and diseases. Notably, by integrating Causal Forests and Double Machine learning with clinical phenotypes, RNAcompare provides a novel approach to bypass traditional batch correction methods.

**Conclusion:** We introduce RNAcompare, a computational platform designed to compare clinical and multi-omics data across diverse patient cohorts in real-time. This tool supports both user-generated and publicly available datasets, offering a robust solution for identifying phenotypic similarities and enhancing our understanding of complex diseases such as RA and HF.

The platform is available at https://github.com/tangmingcan/RNAcompare.

## BACKGROUND

Autoimmune and autoinflammatory diseases incorporate a heterogeneous group of chronic diseases with substantial morbidity and mortality, thereby posing significant health challenges. While there have been substantial advances in the treatment of RA and related rheumatic immune-mediated conditions, based on improved understanding of the underlying disease pathogenesis, there remains significant unmet clinical needs. Treatment responses remain unpredictable and inadequate response is common; some patients may respond to drugs with one mode of action, while others may respond to another drug or not at all ^1^. Furthermore, although recently with the development of machine learning, signatures were found by many previous studies connecting to drug resistance in RA, it still lacks a solid theory to inter-connect patients’ clinical heterogeneity and transcriptomic evidence with a generalised drug-independent treatment outcome ^2–4^.

The integration of multi-omics data has emerged as a promising approach to elucidate the molecular underpinnings of RA and improve treatment strategies. However, this integration faces significant challenges, notably batch effects and tissue-specific variations. Batch effects, arising from technical variations, can confound true biological differences. A recent review emphasizes the profound negative impact of batch effects in large-scale omics studies and underscores the urgent need for effective correction strategies ^5^; Identical molecular signatures may exhibit contrasting regulatory patterns across different tissues, complicating data interpretation. For instance, L1CAM interactions and Notch signaling have been observed to be downregulated in blood samples yet upregulated in dorsal root ganglia samples of RA patients^6^, highlighting the necessity for context-specific analyses.

RNAcare^6^ was developed based on tools of DEG analysis such as Phantasus^7,8^, CFViSA^8^, AmiC^9^, using Django, Regularisation Regression^10,11^ and distributed framework for accelerating the analysis process. It provides insightful results in terms of pain, fatigue and drug response for RA treatment. However, it still cannot help researchers analyse deeper: comparing data across multi-omics levels is still challenging and time consuming; batch correction may remove biological meanings; didn’t explore the synergic effects across metagenes; can’t find similar patterns across tissues because when combing them, they appear to be separate in UMAP^12^/PCA^13^ plots.

To address these complexities and bridge the gap between clinical heterogeneity and multi-omics data, we introduce RNAcompare. This development of RNAcare offers an automated pipeline that seamlessly integrates clinical and multi-omics data for clinicians, facilitating the identification of both differences and similarities across cohorts through advanced machine learning algorithms. By enabling comprehensive, multi-tissue/diseases analyses, RNAcompare aims to enhance our understanding of disease pathogenesis and inform the development of more effective, personalized treatment strategies.

## IMPLEMENTATION

The overall aim of RNAcompare is to provide a web-based tool simplifying the integration and analysis of transcriptomics and clinical data showing the analysis workflow in tabs as outlined in Figure 1 and described later in more detail.

**Figure 1.**
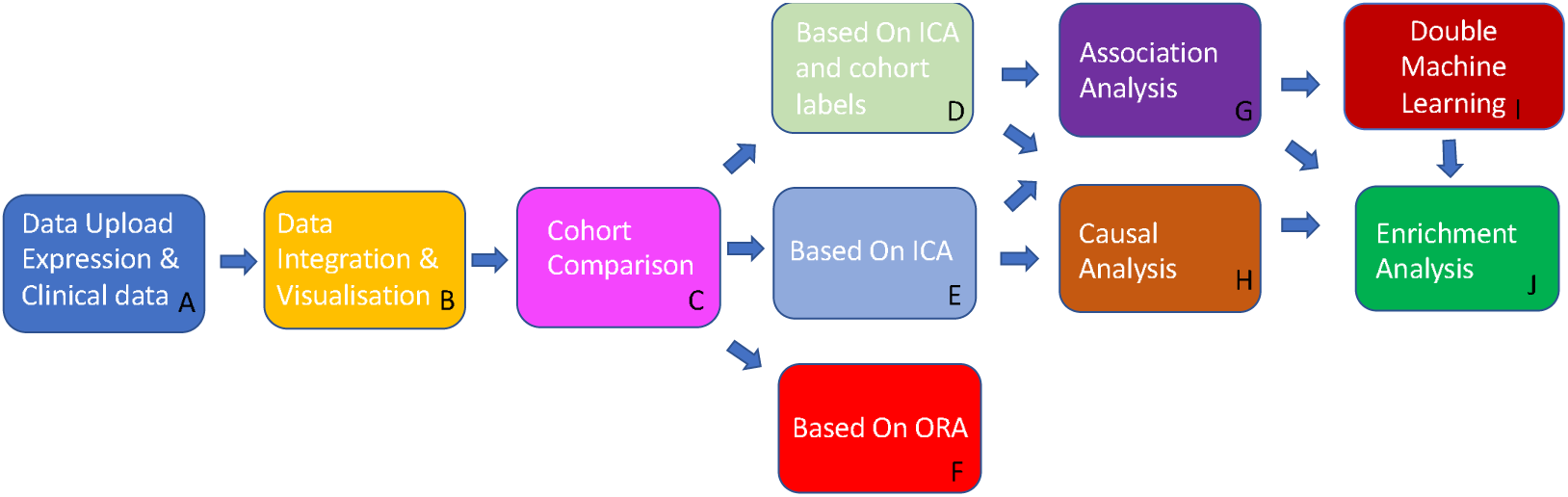
RNAcompare work flow. (A) In the first tab, user can upload transcriptomic, proteomic and clinical data (optional); (B) In Data Integration & Visualisation, users can combine different datasets and have the option to select different integration method and visualise the processed result; (C) Cohort Comparison is an entrance for the following 3 modules; (D) Comparison based on ICA and cohort labels allows users to apply Independent Component Analysis (ICA) on the cohorts separately, then do a combination; (E) Comparison based on ICA allows users to apply ICA on the whole dataset; (F) Comparison based on Over Representation Analysis (ORA) allows user to compare two cohorts directly based on their pathway enrichment score calculated by ORA. (G) Association Analysis is to find relationship between transcriptomic data, clinical data and the user designated label using machine learning algorithms; (H) Causal Analysis is to introduce causal inference model analysing causal relationship and reduce selection bias and handle heterogeneity. (I) Double Machine Learning without batch correction; (J) Enrichment Analysis for the metagene of interest.

RNAcompare was developed in Python, utilizing several open-source packages. The webserver was built using the Django framework, adhering to the FAIR (Findable, Accessible, Interoperable, and Reusable) principle. The platform employs the Plotly graphics system for generating interactive visualisations on the fly. The platform can be installed locally from https://github.com/tangmingcan/RNAcompare.

To enable several users working on the system, Django is used to allow group and authentication management. System administrators can easily assign group roles to specific users, enabling them roles to only view and operate the permitted data.

### Datasets

We introduced the following RA and HF datasets (Table 1) – Optimal management of RA patients requiring Biologic Therapy (ORBIT^14^), Pathobiology of Early Arthritis Cohort (PEAC^15^), RA-MAP^16^ as part of the IMID-Bio-UK consortium and GSE135055^17^.

**Table 1.**
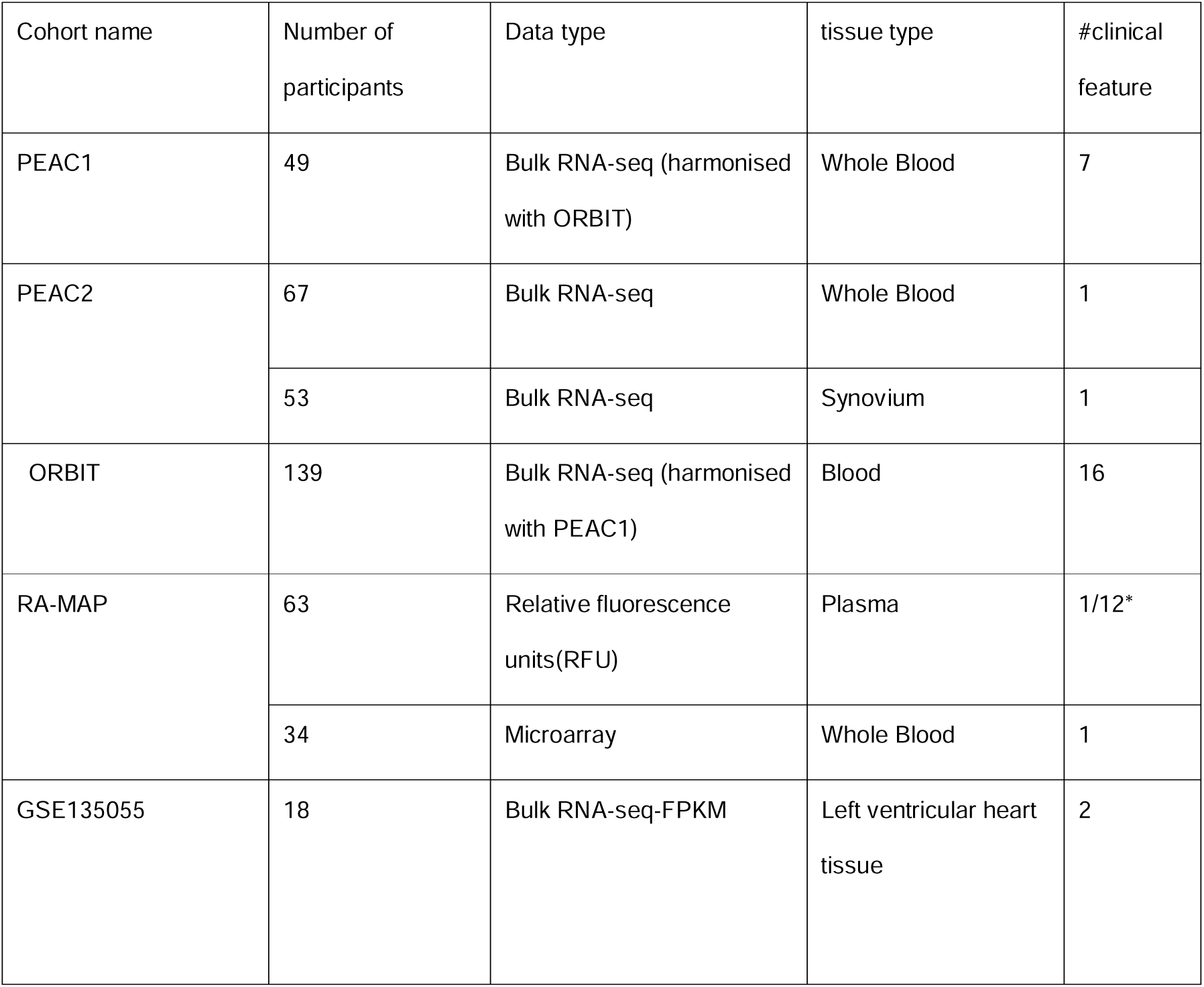
Overview of PEAC, ORBIT and GSE135055 used in RNAcompare. *For RA-MAP, when it is used for comparison between different tissues, we just use pain VAS; when it is used for comparison between different techniques (RNA-seq and Microarray), we used 12 clinical features.

| Cohort name | Number of participants | Data type | tissue type | #clinical feature |
| --- | --- | --- | --- | --- |
| PEAC1 | 49 | Bulk RNA-seq (harmonised with ORBIT) | Whole Blood | 7 |
| PEAC2 | 67 | Bulk RNA-seq | Whole Blood | 1 |
|  | 53 | Bulk RNA-seq | Synovium | 1 |
| ORBIT | 139 | Bulk RNA-seq (harmonised with PEAC1) | Blood | 16 |
| RA-MAP | 63 | Relative fluorescence units(RFU) | Plasma | 1/12* |
|  | 34 | Microarray | Whole Blood | 1 |
| GSE135055 | 18 | Bulk RNA-seq-FPKM | Left ventricular heart tissue | 2 |

The Pathobiology of Early Arthritis Cohort (PEAC) was established with the aim to create an extensively phenotyped cohort of patients with early inflammatory arthritis, including RA, linked to detailed pathobiological data. The clinical and transcriptomic data can be found on EBI ArrayExpress with accession E-MTAB-6141. When analysing with ORBIT, we used processed PEAC1, downloaded directly from RNAcare after its own qualification control; When analysing based on two tissues, we used the original PEAC from the source mentioned.

ORBIT was a study comparing Rituximab to anti-TNF treatments. Blood was taken from patients before drug treatment. RNA-Seq data are being submitted to Array Express as pseudo-anonymized data.

The RA-MAP Consortium is a UK industry-academic collaboration to investigate clinical and biological predictors of disease outcome and treatment response in RA, using deep clinical and multi-omic phenotyping. The raw data were retrieved from accessions GSE97810 and GSE97948 on ArrayExpress. The clinical data can be found here: https://doi.org/10.6084/m9.figshare.c.5491611.v1 GSE135055 was a study about Heart Failure (HF), collected from left ventricular heart tissue from 21 HF patients and 9 healthy donors as study cohort and generated multi-level transcriptomic data. We just used 18 patients with Dilated Cardiomyopathy (DCM), a condition where the heart’s left ventricle (the main pumping chamber) becomes enlarged and weakened, reducing its ability to pump blood effectively.

In the beginning of case study 5, we used simulated data to prove the validity by Causal Forests^18^/DML for recognizing significant metagenes across heterogeneous samples, which is a foundation for the later studies. We designed 2 groups: XA and XB, representing 2 totally different status of patients with shared metagene names from X0-X20, but no overlap values within the same pathway for two groups, then we assigned the similar abs(coefficient) to the two groups to compare the results.

### Clinical data used from the cohorts

RA disease activity was assessed using the validated DAS (Disease Activity Score using 28 joint counts) score, which was calculated from the original recorded 28 joint counts plus a blood marker of inflammation, typically the erythrocyte sedimentation rate (ESR) or the C-reactive protein (CRP) level^19^; Pain VAS^20^ is short for pain Visual Analog Scale (VAS) for self-reported pain, which is a unidimensional measure of general pain intensity, used to measure patients’ current pain level. The pain VAS is not specific to RA and has been widely used in a range of patient populations, including those with other rheumatic diseases, patients with chronic pain, cancer, or even cases with allergic rhinitis^21^, which provides an possibility for integration of different diseases for our later discussion.

HYNA (New York Heart Association^22^) is a system used to classify the severity of HF based on a patient’s symptoms and physical activity limitations. It has four functional classes: Class I: No symptoms and no limitations in ordinary physical activity; Class II: Mild symptoms and slight limitation during ordinary activities; Class III: Significant limitation in activity due to symptoms; comfortable only at rest; Class IV: Severe limitations; symptoms present even at rest.

To combine the disease severity of both diseases, we min-maximised each severity index (DAS and HYNA) to 0-1 respectively, and then multiplied by 100. The second step is not necessary. DAS score in RA-MAP is standardised. So, in case study 9, we first standardised DAS score in PEAC+ORBIT, then combined its clinical data with RA-MAP’s.

All this clinical information is stored on csv files and, after curation, loaded into RNAcompare.

### A Data Upload

The user has the option to use data uploaded previously, upload their data on the fly or a combination of both. As proof of concept, we included the five datasets described above, including clinical and expression data. RNA-Seq requires a read count matrix, which will be normalised later. Other omics data need to be normalised initially before uploading. Clinical data are uploaded as a table. The aim of normalisation is for association analysis, but for Causal analysis, normalisation is not necessary.

### B Data Integration & Visualisation

For association analysis, data integration & visualization is recommended for finding differences across cohorts. This is applied for situations where batches are from the same tissue and same disease. Integration will help increase the number of samples while removing batch effects.

Similar to RNAcare, RNAcompare detects whether the format of the expression data are integers or non-integers before data integration. For integers, the platform handles the expression data as RNA-Seq data, which are transformed from raw counts to CPM (count per million). For non-integers, for example microarray/proteomic data, the platform handles the expression data as normalised data, so users need to pre-normalize these data types if they want.

The user has the option to log1p transform their data. RNA-Seq data is often highly skewed, with large differences in scale between genes. Log transformation helps stabilise the variance. If this option is selected, the gene expression data will undergo log1p transformation and clinical numeric data will remain the same. Conversely, if the log1p transformation is not selected, the gene expression data will remain unchanged as well.

Next, the user has two options for the integration, Harmony and Combat, and three options for feature reduction, principal component analysis (PCA^13^), t-distributed stochastic neighbour embedding (t-SNE^23^) and Uniform Manifold Approximation and Projection (UMAP^12^). In general, it is unclear a priori which integration will generate the best data, and this will vary for different datasets, the user has different options for this in RNAcompare.

After integration, user can view the result based on feature reduction method they selected before association analysis.

### C Cohort Comparison

In this section, users have options to choose how to compare different cohorts. We provide 3 different level options for result consistency check. Comparison based on Independent Component Analysis (ICA^24^) and cohort labels; Comparison based on only Independent Component Analysis; Comparison based on Enrichment Score with Over Representation Analysis(ORA). Their pros and cons will be discussed later in their respective modules. After users choose the option, a new page corresponding to the method will be opened.

Cohorts can have the different definition, depending on the scenarios. In our case studies, it is a categorical label representing the heterogeneous attribute, such as drugs, tissue types or disease types. Other options can be a label standing for 2 different stages of one disease. As we all know DEGs, sensitive to batch correction, is used for finding the different expression genes between the cohorts. Our platform is trying to looking for similarities, where records from different cohorts with strong heterogeneity usually are not easily merged together after UMAP and clustering methods such as K-Means^25^.

### D Based on ICA and cohort labels

Independent component analysis (ICA) attempts to decompose a multivariate signal into independent non-Gaussian signals. We introduced ICA into multi-omics to decompose the gene expression matrix into independent components. Each independent component was treated as a metagene, characterised by a co-expression pattern and was associated with certain meaningful biological pathway. In practice, we suggest that all clinical fields uploaded before must be named as strings starting with “c_”, such as “c_das”, which will be easy for the program to recognise the expression data and apply ICA algorithm.

In the option, ICA is applied to the cohort labels separately. Say, for ORBIT, we have 2 drugs (anti-TNF and Rituximab) corresponding to 2 different response results: responder and non-responder. Then at first, we need to find the similarities between anti-TNF responders and Rituximab responders. Then we need to find the differences between Rituximab responders and anti-TNF non-responders. In each step, ICA will be applied to the two different states separately and we use the fixed features from Rituximab responders as the bridge, connecting anti-TNF responder and non-responder. The advantage is each disease cohort has its independent ICA components, representing its own state where imbalanced records across different cohorts will not be a problem and during the down streaming analysis, we know where the feature comes from. This is very important because during the analysis we may find some malignant feature, and we can know whether cohort A or B has this malignant feature especially when none of the cohorts is a control group. However, the disadvantage is some of the components across different cohorts may be highly correlated. Therefore, a feature selection method needs to be done after the combination of all components from different cohorts. In our case, we calculate the Pearson correlation matrix^26^, and use the threshold (0.65 as default) set by users to exclude overlapping features.

Or we can first group the above-mentioned records into 2 cohorts: responders and non-responders before uploading. It all depends on the chosen granularity as long as the number of records in each cohort is enough (at least 10, we recommend).

### E Based on ICA

In the option, ICA is applied to all the cohorts at one time. The advantage is that it decreases the possibility of generating overlapping components. The disadvantage is if cohorts are imbalanced, ICA may not capture the specific component, and as mentioned user will not know whether a malignant feature is more closed to cohort A or B if none of them is a control group. For the former issue, Synthetic Minority Oversampling Technique (SMOTE^27^) will be a potential solution via over-sampling the small-sized cohort and under-sampling the large-sized cohort. Another down side is that as the total number of the records processed by ICA increases, its final result may be not stable. To solve this, user needs to increase the number of iterations and number of components to get a relatively stable result. However, it will challenge the time complexity of the online real-time analysis.

### F Based on ORA

Over-representation analysis (ORA) is used to determine which a priori defined gene sets are more present (over-represented) in a subset of “interesting” genes than what would be expected by chance^28^. To infer functional enrichment scores, we will run the ORA method. As input data, it accepts an expression matrix. The top 5% of expressed genes by sample are selected as the set of interest. The final score is obtained by log-transforming the obtained p-values, meaning that higher values are more significant.

We then map the score into pathways according to the annotated gene sets in Molecular Signatures Database (MSigDB^29^). Users can choose different gene sets to compare among cohorts, for example Hallmark or Reactome. In our example, we choose Reactome and cell type signature.

### G Association Analysis

Here, the user has the option to associate selected clinical data parameters with the (harmonised) omics data in order to study phenotypes. We provide Random Forests^30^, XGBoost^31^ and LightGBM^32^ with its corresponding parameters for basic tuning. The data will be split into training and test (20%) datasets. AUC-ROC^33^ or MSE^34^ will be shown according to different cases. This module will provide SHAP^35^ feature importance plot and SHAP dependence plot.

SHAP (SHAPley Additive exPlanations) values are a way to explain the output of any machine learning model. It uses a game theoretic approach that measures each player’s contribution to the final outcome. In machine learning, each feature is assigned an importance value representing its contribution to the model’s output.

SHAP values show how each feature affects each final prediction, the significance of each feature compared to others, and the model’s reliance on the interaction between features.

### H Causal analysis

In the option, users can do causal analysis on treatment efficacy by reducing the selection bias between treatment and control group and handle heterogeneity among patients.

The platform provides Causal Forests based on PCA/ICA processed components for cohorts. Causal Forests is a machine learning method used for estimating heterogeneous treatment effects (HTE) in observational data. It is an extension of Random Forests designed specifically for causal inference.

The reason why we provide both PCA and ICA processed components is because we think when focusing on HTE, we think PCA will be more robust. However, when focusing on explainable components, ICA will be better.

When using this module, user needs to designate T and Y as the parameters. T is the label representing treatment effect or the predefined cohorts, which we limit only for categorical variables; Y is the dependent variable for phenotype.

### I Double Machine Learning

After that we generialised the algorithm to double machine learning (DML), which means two machine learning models to overcome Causal Forests’ limitation.

We then extend Causal Forests and DML to the similarity exploration across different tissues, omics levels and diseases. Under this situation, data harmalisation step is not necessary, which will avoid from potentially removing the biological meaning.

### J Enrichment analysis

As the previous analysis will return components of interest, this tab helps to identify biological pathways and signature sets significantly enriched in the dataset.

Another feature of RNAcompare is to download all the results of the different analysis steps, results as csv and figure as SVG for further processing and use in publications.

## Results

Here we present the first tool that allows users without bioinformatics skill to integrate clinical and transcriptomics data, establish an automated pipeline, combining patients’ clinical, multi-omics evidence and phenotypes all together, looking for differences and especially similarities/crosstalk between cohorts by integrated machine learning algorithms.

### Case Study 1: find similar baseline pathways for two response groups with different drug treatments

Resistance to drug treatment is a major problem in treatment responses of RA^36^. Approximately 30%-40% of patients fail to respond to anti-TNF treatment.

Until now, most studies focus on one drug treatment without considering patients clinical parameters. For example, Thurlings et al. showed that there was a better clinical response to Rituximab in the type I IFN low signature group^37^. Hennie G Raterma et al. also showed that the type I IFN signature towards prediction of non-response to Rituximab in rheumatoid arthritis patients^38^. However, they didn’t generalise this finding; Besides, Shengjia Chen et al. introduced a Drug Response Prediction framework using machine learning to model drug effectiveness in RA patients^39^. The study proposed a novel two-stage machine learning framework to predict patient responses to anti-TNF treatments, demonstrating promise in supporting clinical decisions based on electronic health record information.

Nonetheless, they didn’t explain the mechanisms underlying the model. It seems there still exists a gap between explainable machine learning model and generalised drug resistance studies. Although in our previous paper^6^, we tried to correlate the pathways influenza infection with HSP90AA1 to type I IFN signaling as the major contributor influencing drug resistance, the conclusion was still limited to anti-TNF. Here, we propose RNAcompare with 3 different granularities to generalise this conclusion.

Figure 2 shows the 1^st^ level granularity. We performed ICA separately for three groups: anti-TNF responders (ICA1), Rituximab responders (ICA2), and anti-TNF non-responders (ICA3). We then mapped Rituximab responders onto the ICA1 space, anti-TNF responders onto the ICA2 space, Rituximab responders again onto the ICA3 space, and anti-TNF non-responders onto the ICA2 space. This allowed us to combine and compare anti-TNF responders with Rituximab responders, as well as anti-TNF non-responders with Rituximab responders, using the Rituximab responders as a bridge. We can see Figure 2-A shows how similar between anti-TNF (0, on the left) and Rituximab responders (1, on the right) via SHAP values. Figure 2-B shows how different between Rituximab responders (0, on the left) and anti-TNF non-responders (1, on the right) via SHAP values.

**Figure 2.**
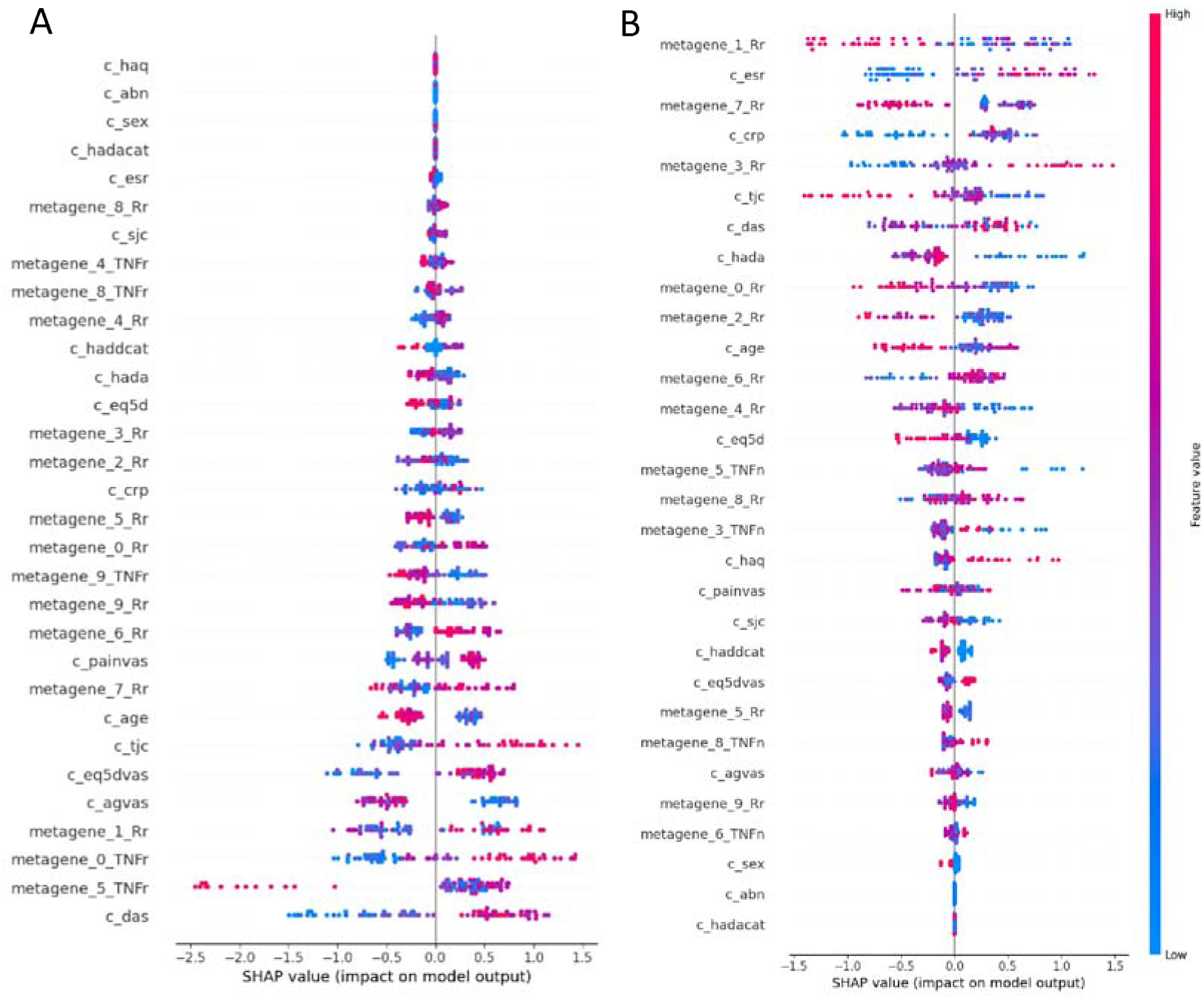
Comparison of anti-TNF responder, Rituximab responder and anti-TNF non-responder; (A) Comparison of anti-TNF responder (0) and Rituximab responder (1); (B) Comparison of Rituximab responder (0) and anti-TNF non-responder (1).

We are looking for the bridge with the suffix ‘Rr’ between the 2 plots. Therefore, in Figure 2-A, metagene_8_Rr, metagene_4_Rr, metagene_3_Rr, metagene_2_Rr and metagene_0_Rr stand for how similar for the two response groups because the red and blue dots almost merge together. While in Figure 2-B, those metagenes make response group and non-response group separate far away. The feature importance for every feature can be downloaded to compare. Therefore, we can calculate the difference of metagene_8_Rr diff(metagene_8_Rr) = 0.19215-0.03598=0.15617 and others with their Enrichment analysis (Table 2). Also Figure 3 lists the significant pathways for those metagenes. Of note, for metagene_4_Rr, its significant pathways including influenza infection, however its relationship contradicts what we found in RNAcare. This may be because HSP90AA1 is involved in both type I and type II IFN pathways.

**Figure 3.**
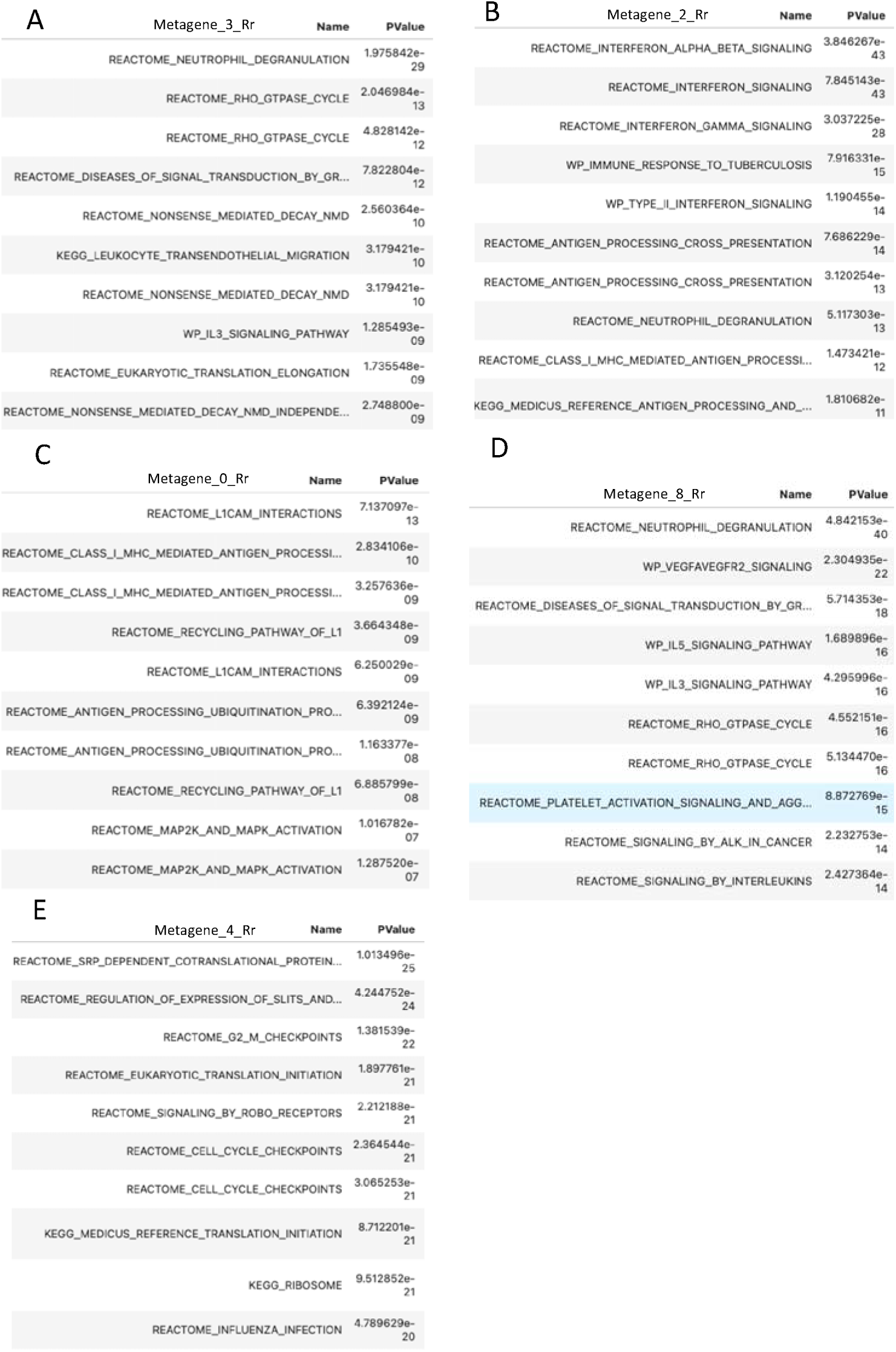
Pathway analysis for significant metagenes; (A) Pathway analysis for metagene_3_Rr; (B) Pathway analysis for metagene_2_Rr; (C) Pathway analysis for metagene_0_Rr; (D) Pathway analysis for metagene_8_Rr; (E) Pathway analysis for metagene_4_Rr.

**Table 2.** Metagene explanation.

| Metagenes | Diff | Explanation | Relationship to drug resistance |
| --- | --- | --- | --- |
| metagene_3_Rr | 0.29601 | In Figure 3-A, metagene_3 is significant in Rho GTPase cycle and eukaryotic translation elongation. This align with what we found previously connected to Type I IFN, namely the expression level is higher in anti-TNF non-responder compared to Rituximab responder. Rho GTPase cycle induces type I and II IFN production while prevents type I IFN-induced apoptosis in B cells<sup>40,41</sup>. | + |
| metagene_2_Rr | 0.20031 | In Figure3-B, metagene_2 is significant in IFN. Since we already know type I IFN is negative to drug response, we assume metagene_2 functions more as type II IFN. Therefore, we know that low fatigue at baseline is correlated with good drug response based on RNAcare<sup>6</sup>. From Figure 2, we can also assume anti-TNF responders' type II IFN > Rituximab responders' > anti-TNF non-responders'. | - |
| metagene_0_Rr | 0.16945 | In Figure 3-C, metagene_0 is significant in L1CAM interactions, which was once discussed related to pain in RNAcare<sup>6</sup>. Therefore, we know that low pain at baseline correlated with good drug response. | - |
| metagene_8_Rr | 0.15617 | In Figure 3-D, metagene_8 is similar to metagene_3_Rr but also significant in interleukins signaling, interacted with JACK1. Its important genes include IL6R. IL-6 is a key cytokine in the JAK/STAT3 pathway, involved in tumor progression through inhibition of cancer cell apoptosis, stimulation of angiogenesis, and drug resistance. | + |
| metagene_4_Rr | 0.10408 | In Figure 3-E, metagene_4 is significant in eukaryotic translation initiation and influenza infection. This aligns with what we found previously connected to Type I IFN. However, from Figure 2-B, we can see the almost half of the anti-TNF non-responders' expression level is lower compared to Rituximab responder. **This may be because of the interaction with RHO GTPase cycle via Type I IFN. This further analysis will show in Appendix-2.** | **Non-linear** |

Considering metagene_3_Rr and the additional drug Rituximab, we found that RHO GTPase cycle pathway also includes gene HSP90AA1, which means it might be highly correlated with influenza infection pathway. Then we analysed their synergic effects to pain in Appendix-2.

On the other hand, we can also see the ESR is significantly different between responder (lower) and anti-TNF non-response group (higher).

Figure 4 shows the 2^nd^ level granularity, where we combined anti-TNF responders and Rituximab responders together as the responders (1) and anti-TNF non-responders and Rituximab non-responders together as the non-responders (0). We can see the combination increases the difficulty of model training based on the feature plot because not too many important features separate two cohorts far away as before in Figure 2-B. Through Figure 4-A, we can see ESR is still negatively related to the response group; metagene_0_r is positively related to the response group, metagene_1_r is obvious positively related to the response group. However, the most important metagene_2_r exhibits a non-linear relationship to the response group. Therefore, we draw a SHAP dependence plot for metagene_2_r (related to metagene_9_n, Figure 4-B), and did enrichment analysis for the 4 metagenes, see Table 3.

**Figure 4.**
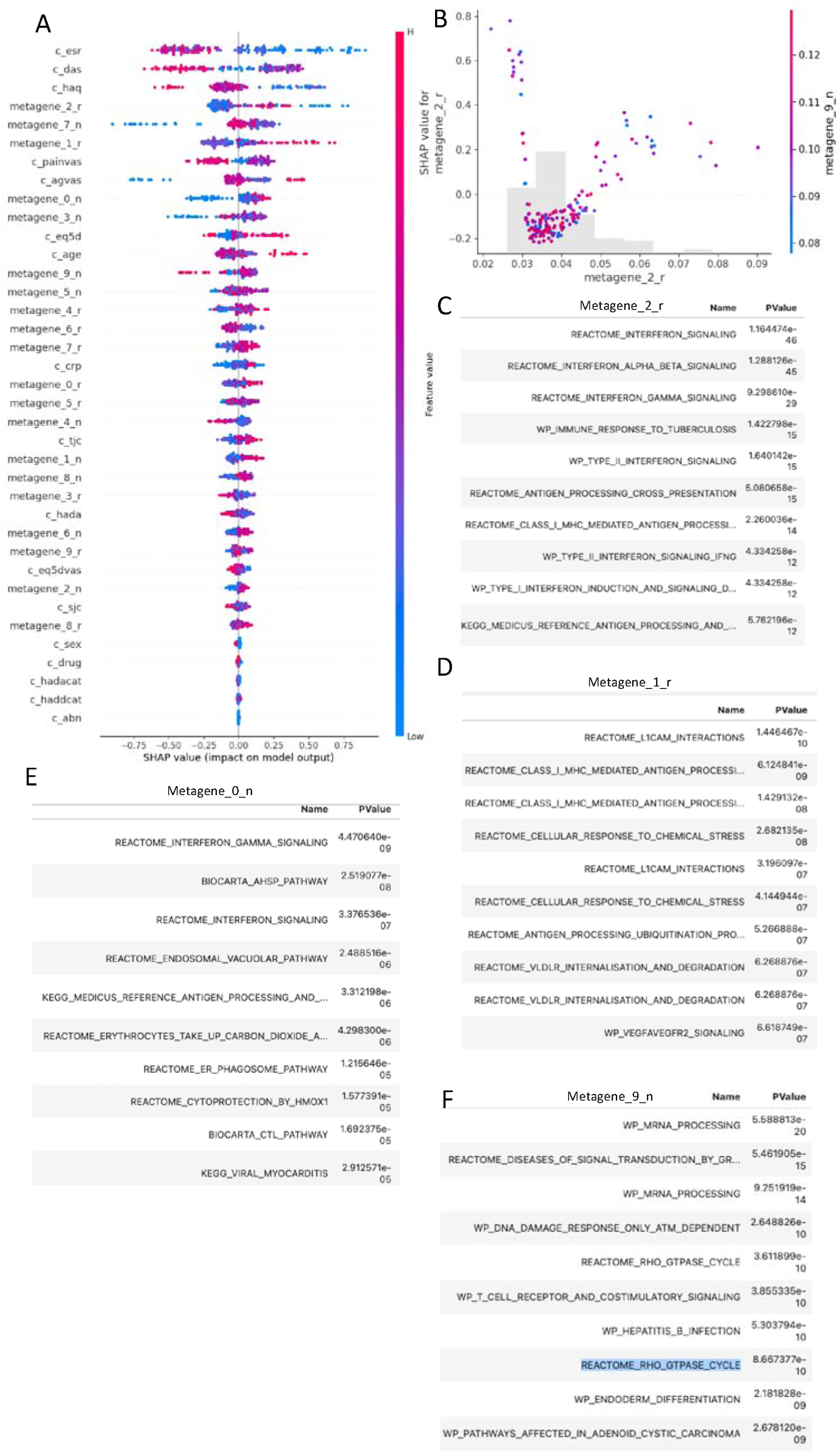
2nd level result. (A) SHAP feature importance plot; (B) SHAP dependence plot between metagene_2_r and metagene_9_n; (C) Pathway analysis for metagene_2_r; (D) Pathway analysis for metagene_1_r; (E) Pathway analysis for metagene_0_n; (F) Pathway analysis for metagene_9_n.

**Table 3.** Metagenes explanation.

| Metagenes | explanation | relationship to drug resistance |
| --- | --- | --- |
| metagene_2_r | In Figure 4-C, it is significant in both type I and type II IFN signaling, which is the main reason making the relationship non-linear, based on the interpretation of metagene_9_n and SHAP dependence plot in Figure 4-B. | **Non-linear** |
| metagene_1_r | In Figure 4-D, it is significant in pain related genes. | - |
| metagene_0_n | In Figure 4-E, it is significant in type II IFN signaling (fatigue related genes). | - |
| metagene_9_n | In Figure 4-F, it is significant in RHO_GTPASE_CYCLE, related to type I IFN signaling. | + |

Figure 5-A shows the 3^rd^ level, where we applied ICA over all (0-Non-responders, 1-Responders). RNAcare also has the similar function. Generally speaking, metagene_1 is positive and metagene_5 is partly negative to the drug response. So, we did enrichment analysis for the two genes. Figure 5-B shows metagene_1 is enriched in pain-related pathway, which is positive to the response as expected.

**Figure 5.**
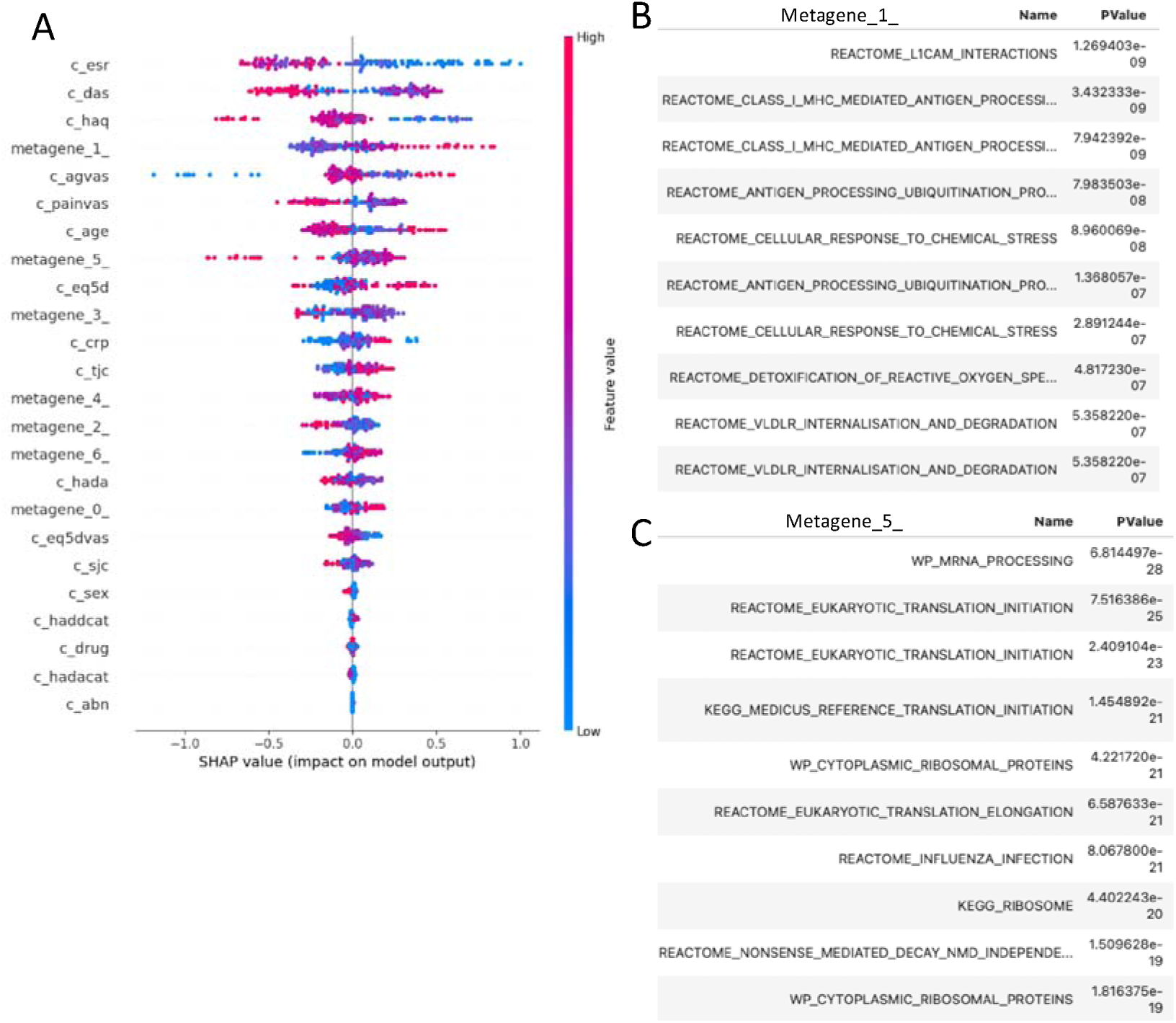
3^rd^ level result. (A) SHAP feature importance plot; (B) Pathway analysis for metagene_1; (C) Pathway analysis for metagene_5.

Figure 5-C shows metagene_5 is enriched in eukaryotic translation initiation and influenza infection which we discussed before about Figure-2AB metagene_4_Rr.

Conclusion: Using ICA to decompose data at a lower granularity based on phenotypes will help explain the underlying patterns better; Type I IFN is negative to drug response; Type II IFN is positive to drug response (although it will increase the disease activity & allograft rejection risk, talk later in case study 5 and appendix 4); Lower fatigue and pain level at baseline will be helpful to drug response; Rituximab can probably treat patients with higher type I IFN induced by RHO GTPase and part of anti-TNF non-responders with a high expression of influenza infection pathway.

### Case Study 2: based on ORA

To compare with the result in case study 1, we ran ORA based on Reactome gene sets in MSigDB. First, we ran ORA method to calculate the enrichment score, based on the input expression data. By default, the top 5% of expressed genes by sample were selected as the set of interest. We then analysed all the significant pathways based on grouped response and non-response cohorts similar to the procedure of getting DEGs. At last, we plotted the heat-map across four cohorts to see whether the two response groups showed a similar pattern in the specific pathway and vice versa. Figure 6-A shows the result for the differential expression pathways of non-response for the 4 cohorts. We can easily see that non-response group share the similar underlying patterns such as:

ACTIVATION_OF_C3_AND_C5, LRR_FLII_INTERACTING_PROTEIN_1_LRRFIP1_ACTIVATES_TYPE_I_IFN_PRO DUCTION, ALPHA_DEFENSINS, RHOC_GTPASE_CYCLE,

MET_PI3K_AKT_SIGNALING (will analyse in later case).

**Figure 6.**
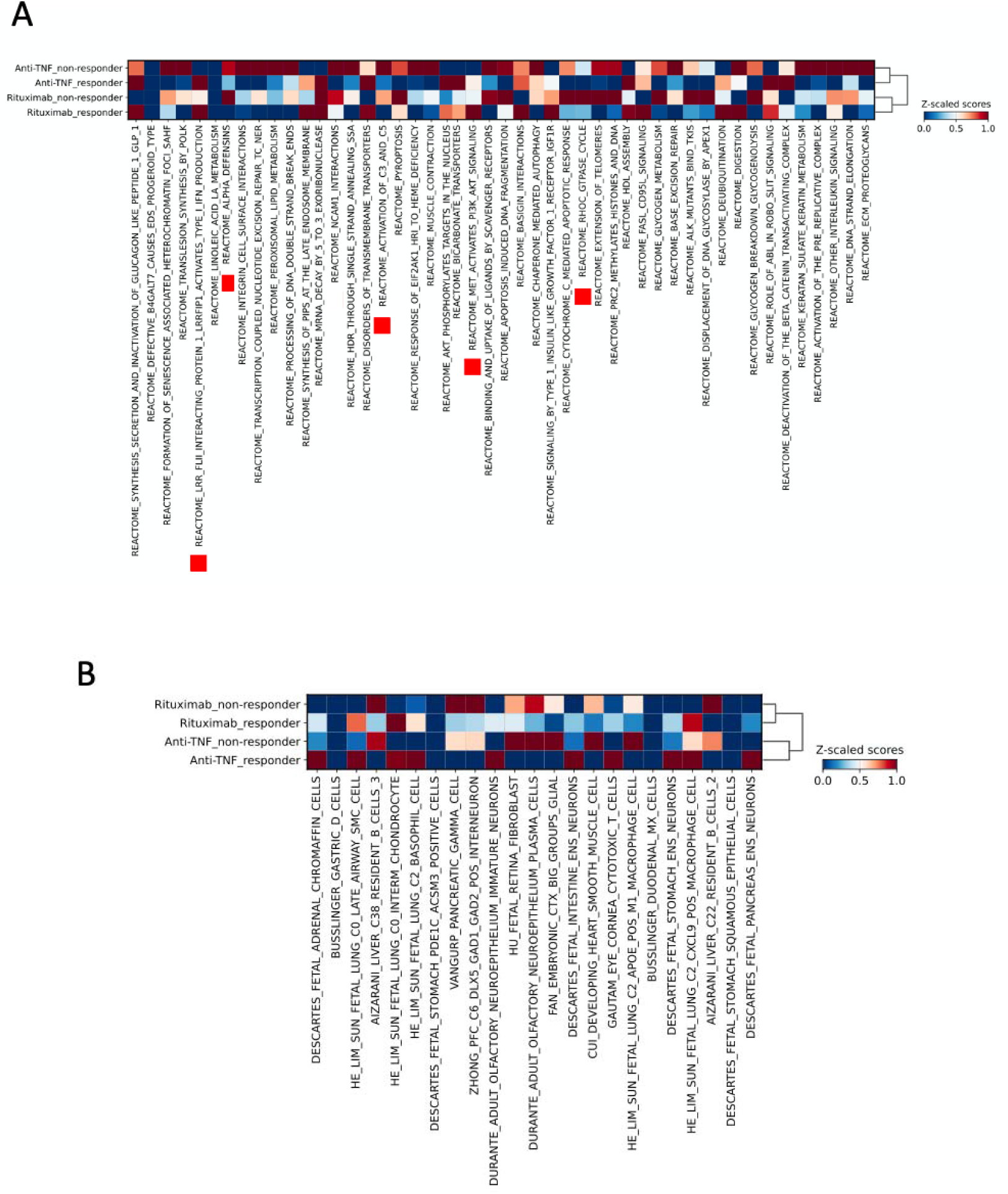
ORA based on 4 different drug response cohorts. (A) using Reactiom database; (B) using cell type signature databaset

There is a strong link between C3/C5 Activation and drug resistance through complement C3 activation in cancer-associated fibroblasts, which drives chemo-resistance and immune evasion via NF-κB signaling^42^. Considering about case study 5.2, we know this pathway may also be correlated to IL-1 related pathway, which has already partly proved by paper^43^.

LRRFIP1 is enriched in Anti-TNF responders and Rituximab patients. It has double roles in mediating type I IFN: the decreased binding of LRRFIP1/GCF2 activates NF-kB and causes over-expression of TNF-α in various autoimmune diseases, such as RA^44,45^ and it also promotes the production of IFN-β^46^.

On the other hand, we can change to another database for a new perspective. Using Figure 6-B we know what cell types are resistant to both drug treatments, such as: liver c22 resident b cells, liver c38 resident b cells, which means Rituximab cannot help as well.

However, we also found issues when trying to integrate with ML for interpretation (see Appendix 3), because the 5% top genes possibly limit the expressiveness and predictive power. One potential solution may be ssGSEA, but time consuming.

### Case Study 3: Causal analysis for the treatment efficacy of two drugs

ORBIT project was aimed to test the hypothesis that using Rituximab would be clinically non-inferior and cheaper compared with TNF inhibitor treatment in biological-treatment naive patients with rheumatoid arthritis. It concluded that Initial treatment with Rituximab is non-inferior to initial TNF inhibitor treatment in patients seropositive for rheumatoid arthritis and naive to treatment with biologicals, and is cost saving over 12 months. However, this conclusion was based just on clinical data without considering patients’ transcriptomic evidence at baseline. Although it claimed the choice of drugs for patients was randomized, we tried to introduce causal machine learning algorithms to reduce the potential selection bias and handle heterogeneity issues and re-evaluated the HTE for the patients.

Jacques-Eric et al. found that among patients with rheumatoid arthritis previously treated with anti-TNF drugs but with inadequate primary response, a non-TNF biologic agent was more effective in achieving a good or moderate disease activity response at 24 weeks than was the second anti-TNF medication^47^; Heike et al. found that patients who did not respond to anti-TNF therapy exhibited a significant expansion of apoptosis-resistant TNFR2+IL23R+ T cells. These cells expressed higher levels of IFN gamma^48^. We already unintentionally covered part of the issue previously in case study 1, that Rituximab responders have lower level of type I IFN expressions compared those of anti-TNF non-responders (Figure 2-B metagene_3_Rr) and Rituximab responders express higher level of type II IFN than anti-TNF non-responders do (Figure 2-B metagene_2_Rr). In this part, we will continue to explore more and focus on issues of selection bias and heterogeneity.

For data engineering part, we applied Principal Component Analysis (PCA) and ICA separately on the whole expression data at baseline, then combine the processed components with clinical parameters for comparison. The reason why we introduced PCA, is because we think PCA might be better compared to ICA in estimating HTE as a baseline, but ICA will be better for feature explanation. Additionally, we added features like IMF score, which generally refers to an index or score used in immune-related research to assess the immune microenvironment of tumors or other tissues, often involving the composition of immune cells (CD8_T, CD4_T, B cells, NK cells), myeloid cells (Monocytes, Macrophages, Neutrophils, DC), and fibroblasts (Fibroblasts, Myofibroblasts). This type of scoring system can be used to understand the immune landscape and how the presence of different cell types influences the disease environment, including treatment response and prognosis, see the distributions of 3 scores in Figure 7-A.

**Figure 7.**
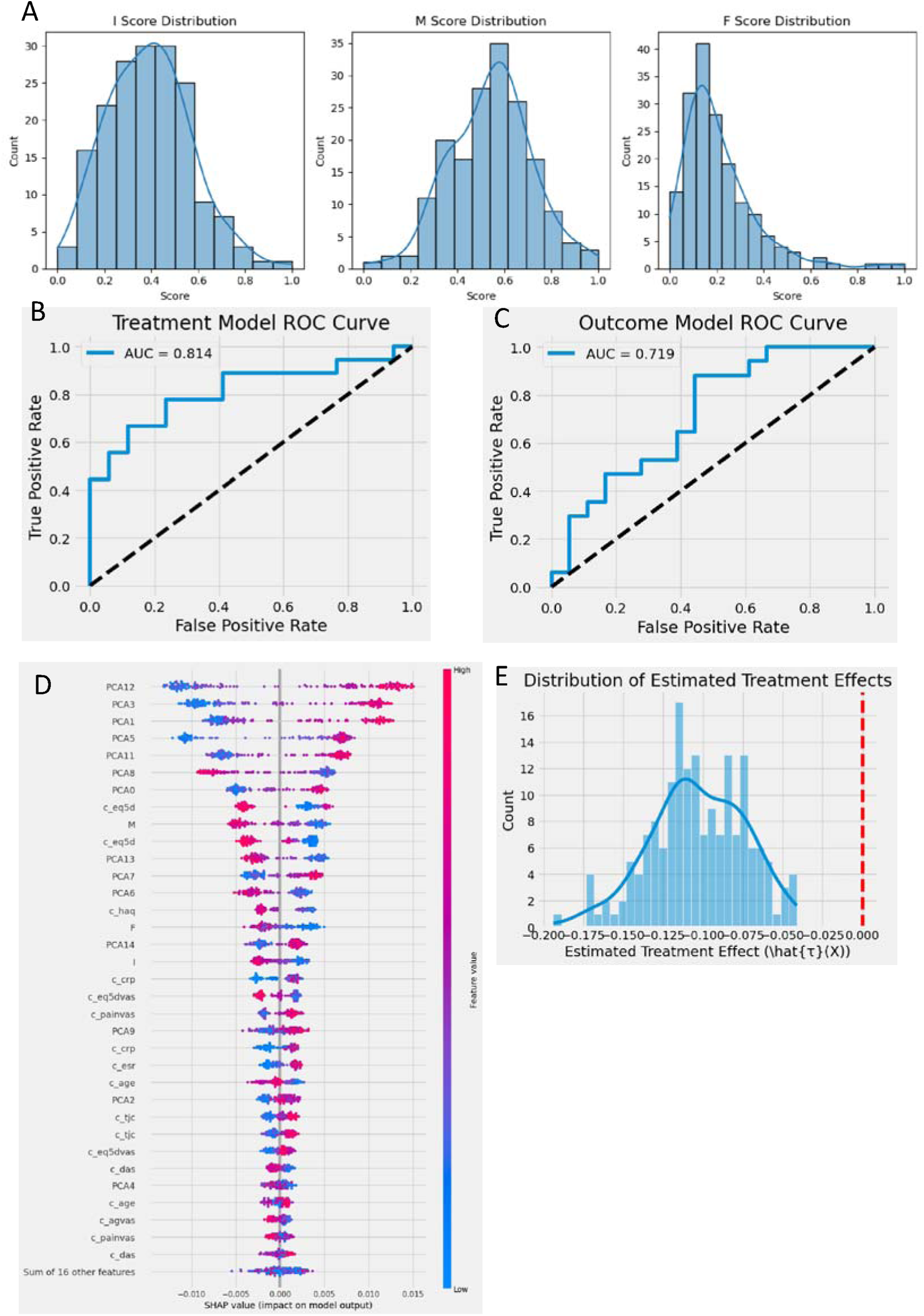
Causal analysis for different treatments using PCA processed data. (A) Distributions of I, M and F scores; (B) Test data - Treatment Model ROC cure for Causal Forests; (C) Test data - Outcome Model ROC curve for Causal Forests; (D) SHAP values feature importance plot to heterogeneity for Causal Forests (E) Distribution of estimated HTE of training data for Causal Forests.

For the modeling part, we used Causal Forests to estimate the HTE (0-anti-TNF, 1-Rituximab), which is commonly used in causal inference combating with selection bias and heterogeneity.

Figure 7-B is the ROC curve of test dataset (0.2 split) for the model we used for the 1^st^ stage of Causal Forests. Since the AUC is 0.814, we assume that the ORBIT project didn’t realise a fully randomised selection of different drugs based on transcriptomic evidence. Figure 7-C is the ROC curve of the same test dataset for the outcome model of Causal Forests. Figure 7-D is SHAP values of feature importance based on Causal Forests. Figure 7-E is the HTE for testing by Causal Forests. We can also see the HTE is less than 0, which means anti-TNF is slightly better on average as well.

When interpreting the results of Figure 7-D, E, we think they shared many commons between models, and for clinical parameter c_eq5d, ranging from −0.59 to 1, where 1 is the best possible health state. It is used in both clinical and research settings, particularly to evaluate the impact of diseases and treatments on a patient’s well-being. So, anti-TNF will be better if patient undergoes a good health condition while Rituximab will reduce the average treatment difference if patient have a bad health condition. Another finding is that anti-TNF will be better for patients with high level expression of M score while Rituximab will reduce the average treatment difference for patients with low level expression of M score. It sounds reasonable because Rituximab targets CD20 on B cells^49^.

Next, we used ICA for further analysis. Figure 8-A, B shows compared with PCA, ICA sacrifices the AUC to get interpretability in return. Figure 8-C shows the SHAP plot for important features influencing the most to the heterogeneity. We can see eq5d, C-Reactive Protein (CRP) and type II IFN (Figure 8-D) are the most important features to the patients’ heterogeneity. It also tells us that Rituximab will be more effective to patients with high level expression of CRP, type II IFN, influenza infection and IL6 involved type I IFN as well as bad eq5d (Figure 8-C-F).

**Figure 8.**
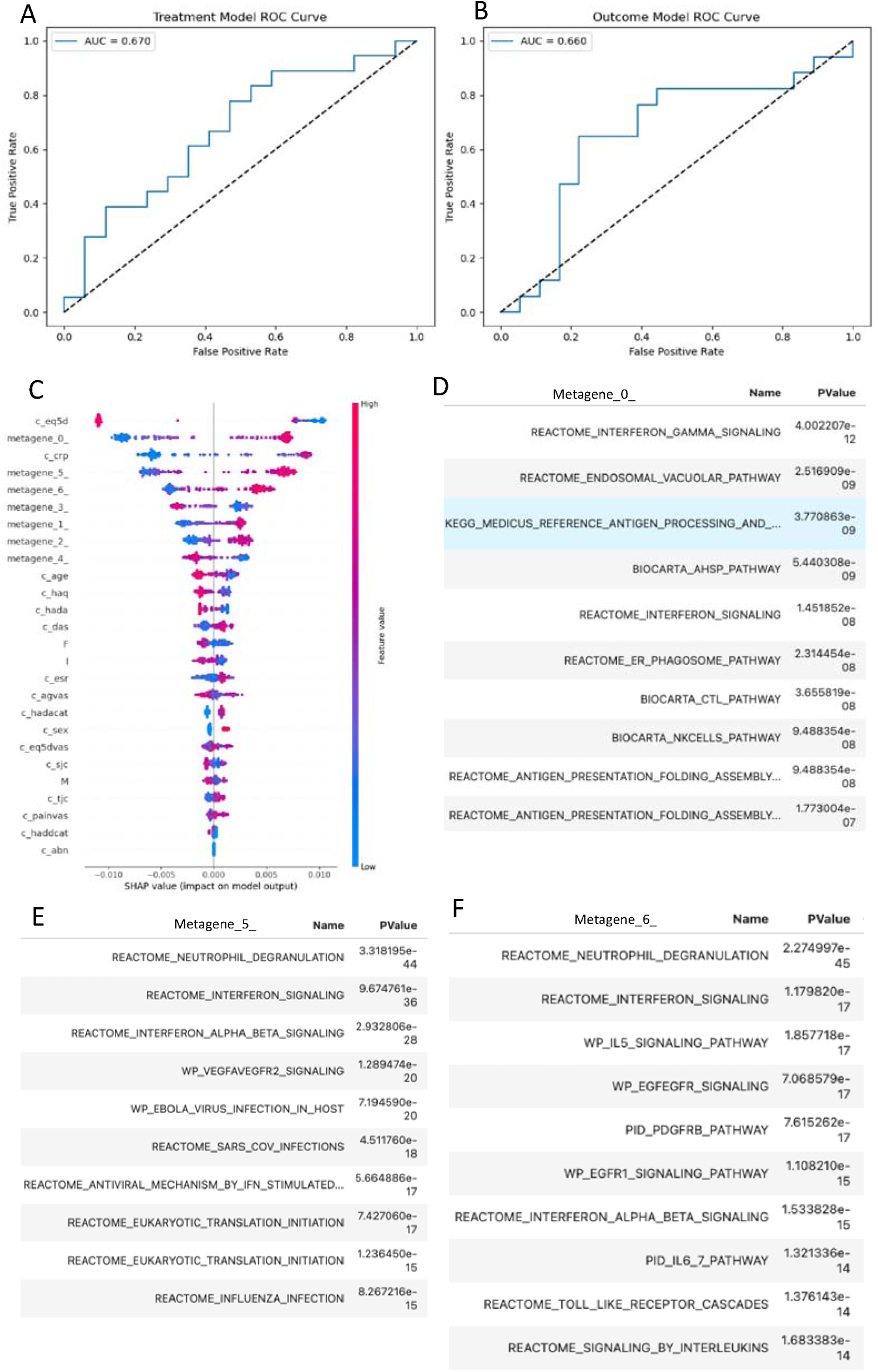
Causal analysis for different treatments with Causal Forests using ICA processed data. (A) Test data - Treatment Model ROC cure for Causal Forests; (B) Test data - Outcome Model ROC curve for Causal Forests; (C) SHAP values feature importance plot to heterogeneity for Causal Forests; (D) Pathway analysis for metagene_0_; (E) Pathway analysis for metagene_5_; (F) Pathway analysis for metagene_6_.

Besides, we can see Causal Forests can detect type II IFN as the most important metagene contributing to the response/non-response heterogeneity using 3^rd^ level granularity data, namely with the increase of type II IFN, those anti-TNF responders can change to non-responders (SHAP value is 0), then further, they will change to Rituximab responders. However, case study 1 with same data level didn’t priorities type II IFN. Although, the final results for the different drugs are totally flipped with the influence by type II IFN, the underlying mechanism/pathway influencing human body is still the same. That is the advantage of applying Causal Forests in this scenario, and this conclusion makes preparation for our further case studies.

Conclusion: We found most top 3 important features contributing to patients’ heterogeneity: eq5d, CRP and type II IFN; In terms of Average HTE, anti-TNF is more effective for test dataset; We discussed under what circumstances, Rituximab will reduce the treatment difference compared to anti-TNF; We proved that Rituximab will be helpful for those anti-TNF non-responders whose type II IFN increased after anti-TNF treatments by Causal Forests.

### Case Study 4: Why aged RA patients have a good response to the treatment

After Case Study 1, we tried to subgroup patients according to their clinical parameters and find common patterns among subgroups based on our previous conclusions. By using the 3^rd^ level granularity features (without IMF scores), we plotted SHAP dependence plot based on patients’ age. Figure 9-A describes the interaction between age and pain. What we found was, patients whose ages between about 55 to 68 (Figure 9-B) exhibit a significant character of being resistant to the drugs while patients aged above 70 receiving good response. Therefore, using RNAcare, we grouped the patients into 3 groups: low aged group (<55, 63 people), middle aged group (55-68, 55 people) and high aged group (>68, 21 people). In Figure 9-C, we can see the middle-aged group is lower in CRP, higher in ESR and lower in metagene_1 which is more related to type I IFN and type II IFN (Figure 9-D). We examined the signatures related to type II IFN in the metagene and found the important ones (Figure 9-E). According to our previous conclusion in case study 1, low expression of type I IFN is good, and low expression of type II IFN is better for anti-TNF rather than Rituximab treatment.

**Figure 9.**
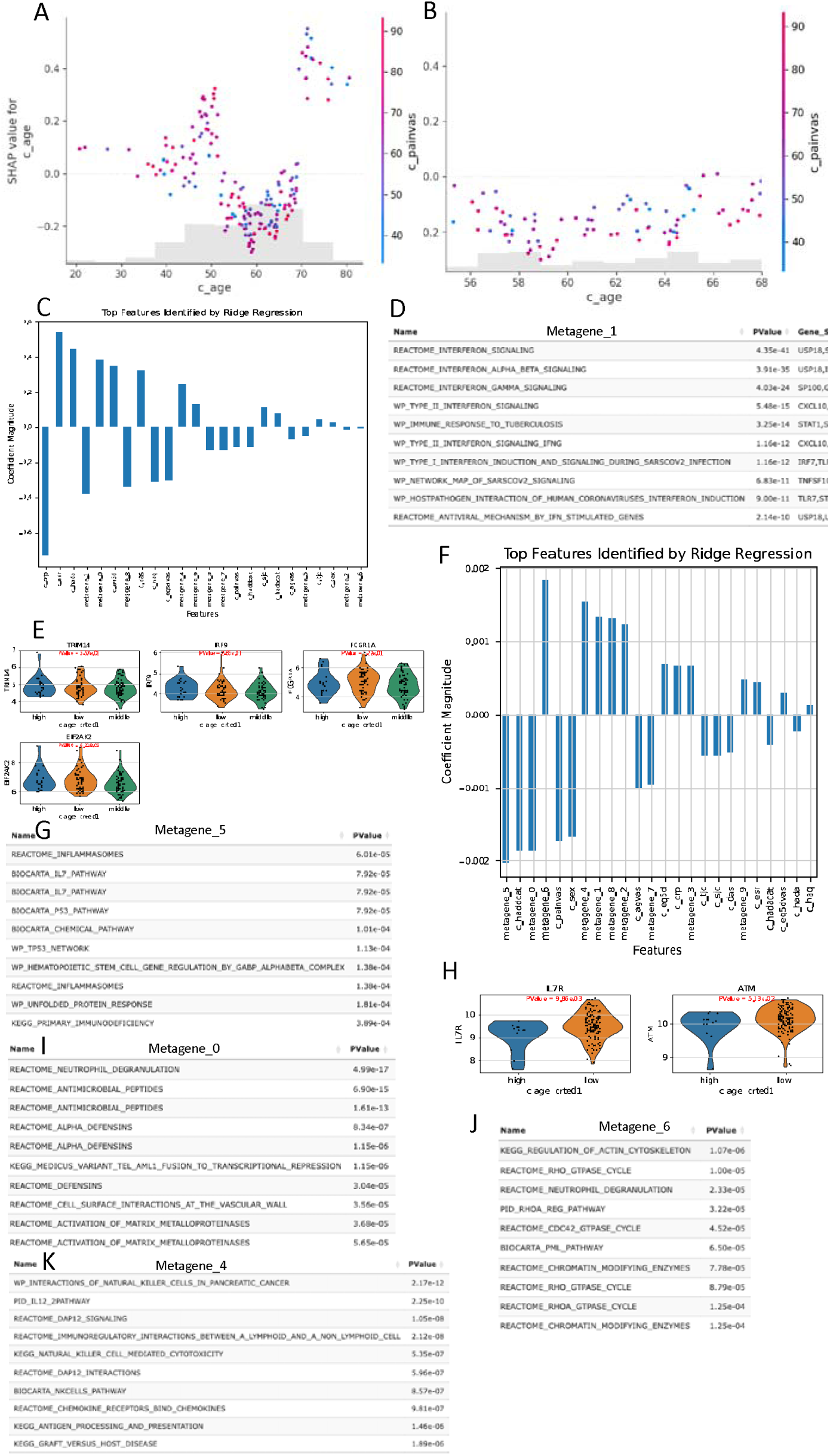
Analysis for aged patients. (A) SHAP dependence plot between age and pain VAS; (B) SHAP dependence plot with a specific age range; (C) Feature importance for middle aged group (55-68) in RNAcare; (D) Pathway analysis of metagene_1 for middle aged group (55-68); (E) Significant biomarkers for type II IFN in metagene_1 for 3 age groups; (F) Feature importance for high aged group (>70); (G) Pathway analysis of metagene_5 for aged group; (H) Significant biomarkers for IL-7 pathway in metagene_5 for two age groups; (I) Pathway analysis of metagene_0 for aged group; (J) Pathway analysis of metagene_6 for aged group; (K) Pathway analysis of metagene_4 for aged group.

Next, we created a new label in RNAcare to check the high aged group (aged above 70, 18 people vs others 121 people). Figure 9-F shows the high-aged group is lower in metagene_5, pain and metagene_0 but higher in metagene_6 and metagene_4. Table 5 shows the corresponding explanation. Therefore, the only reason causing this group ineffective to drug treatment is RHO GTPase cycle pathway, which has already proved by paper that both anti-TNF and Rituximab (indirectly) can potentially reduce its activity in RA through TNF alpha ^50^ and anti-TNF is more direct (c_age in Figure 8-C).

**Table 4.** Causal Forests performance based on test datasets.

| Model | min_pred | max_pred | Average HTE |
| --- | --- | --- | --- |
| Causal Forests | -0.193988 | -0.060212 | -0.09011 |

**Table 5.** Metagene explanation for high-aged group.

| Metagenes | Explanation | Relationship to drug resistance |
| --- | --- | --- |
| metagene_5 | It is enriched in IL7 pathway in Figure 9-G. IL7 is a cytokine essential for the development and maintenance of T and B cells, stimulated by IFN<sup>51</sup>. Low level of IL7 pathway means low level of IFN. Using the previous way, we found its biomarkers in blood: IL7R ant ATM, see Figure 9-H. Generally speaking, anti-TNF treatment will be more helpful to low expression of IL7 pathway for this group. | No define |
| metagene_0 | In Figure 9-I, it is enriched in alpha defensins inducing IL-6 which can enhance the response of neutrophils to type I IFN<sup>52,53</sup>. | + |
| metagene_6 | It is enriched in RHO GTPase signaling in Figure 9-J. | + |
| metagene_4 | In Figure 9-K, it is significant in PID_IL12_2PATHWAY, intricately connected to IFN-γ production through IL-12<sup>54</sup>. | - |

Conclusion: We successfully mapped age parameter to the drug resistance, and found common explainable patterns among subgroups based on our previous conclusions; We also gave different treatment advice to different subgroups characterised by age.

### Case Study 5: Connecting subgroups via DML (same disease)

The reason why we can introduce Causal Forests here is because of the strong heterogeneity across batches. Therefore, Causal Forests can handle this issue and successfully recognise two cohorts during SHAP plot.

### 5.1 Connecting simulated metagenes to phenotype

#### 5.1.1 Causal Forests

Based on Figure 8-C, we can have another interpretation at 3^rd^ level granularity: Initially, we assigned 0 to anti-TNF and 1 to Rituximab for treatment type and are looking for underlying metagenes influencing both cohorts, sharing similar patterns. They might represent common mechanisms in both cohorts, such as inflammation, oxidative stress, or immune activation.

When plotting the SHAP feature importance, these metagenes will have its average HTE close to 0 across samples, and two colours of dots representing cohorts/diseases should overlap or distribute in a sides-symmetric way, while the metagenes of our interest, should have a wider spread representing the phenotype prediction, and because of the heterogeneity there is always a transition from negative to positive SHAP values: In Figure 8-C, we can see metagene_0 looks well sides-symmetric and with the increase of its expression from left side, the efficacy of anti-TNF reduced to the centre and with the decrease of its expression from right side, the efficacy of Rituximab reduced to the centre, which means the treatment effects equal between two drugs driven by metagene_0. However, if we exclude this range of value change (purple dots in the plot), for the rest of the two clusters (blue-antiTNF and red-Rituximab), delta expression of metagene_0_ influences two drugs equally. To view this spread from another perspective, we can check the SHAP dependence plot for the metagene in a 0-centred symmetric way, which means the pathway modifies pain similarly in both cohorts after excluding the overlap: since AUC for T is close to 1, SHAP values should reflect how the feature contributes to pain difference within cohort A before the overlap, and reflect how the same feature contributes to pain difference within cohort B after the overlap, and it is within the overlap that the pathway drives the pain difference close to each other.

Furthermore, we can generalise this conclusion: we don’t necessarily need to look for metagenes with a global 0-centred symmetric spread. Instead, we just need to find those with a local one, namely that in a specific range of expression, the metagene exhibits same patterns for both cohorts.

To approve our assumption, we used simulated data for test. We designed 2 balanced groups: XA (T-0) and XB (T-1) with a sample size of 500 in total (0.2 for test split), representing 2 totally different status of patients with shared metagenes named from X0-X20, but no overlap values within the same x_i_ for both to guarantee AUC(T)=1, then we gave coefficients to X0 within the two groups seperately for the contribution of Y1, Y2. Then we scaled Y1, Y2 separately and combined together.

As shown in Figure 10, In our simulations, we observed that Causal Forests perform better on test data when the within-group variance (σ_i_) is higher and the difference in variances between groups (σ_A_ and σ_B_) is larger. Under these conditions, Causal Forests effectively identify the significant X0 associated with Y. This suggests that the model’s performance depends more on the variance structure than on the actual coefficients of X0. As a result, the range of SHAP values spread doesn’t fully reflect the magnitude of the metagene’s effect but still capture its relative importance within each group.

**Figure 10.**
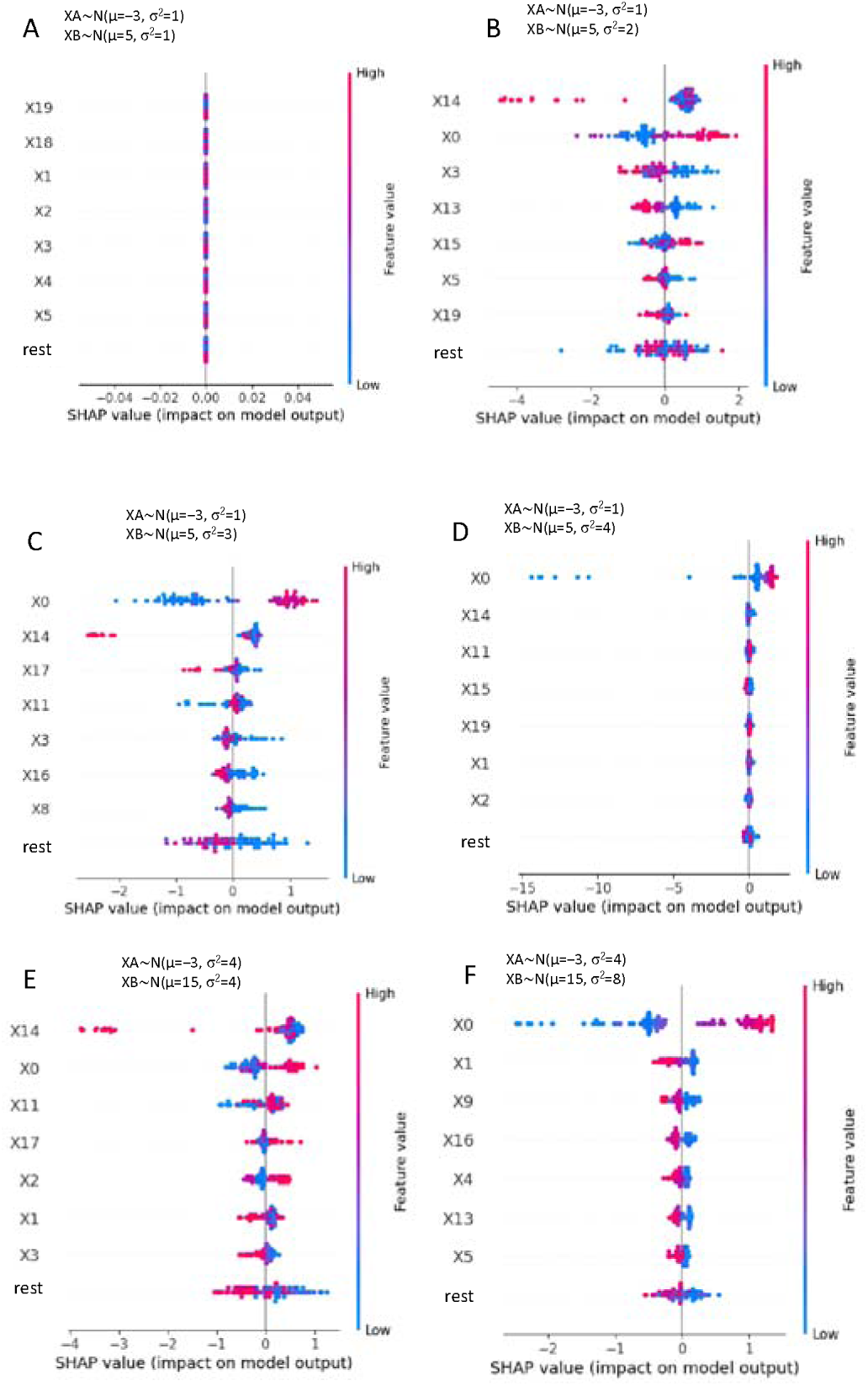
Connecting simulated metagenes to phenotype with abs(coefficients)>1 for X0 in each group. (A) Feature importance by spread for XA ∼N(μ=−3, σ^2^ =1), XB∼N(μ=5, σ^2^ =1); (B) Feature importance by spread for XA ∼N(μ=−3, σ^2^ =1), XB∼N(μ=5, σ^2^ =2); (C) Feature importance by spread for XA ∼N(μ=−3, σ^2^ =1), XB∼N(μ=5, σ^2^ =3); (D) Feature importance by spread for XA ∼N(μ=−3, σ^2^=1), XB∼N(μ=5, σ^2^=4); (E) Feature importance by spread for XA ∼N(μ=−3, σ^2^ =4), XB∼N(μ=15, σ^2^ =4); (F) Feature importance by spread for XA ∼N(μ=−3, σ^2^ =4), XB∼N(μ=5, σ^2^ =8);

#### 5.1.2 (Generalised) Double Machine Learning

After the success of Causal Forests, there is still a limitation: Causal Forests is built on randomised trials, which means the underlying theory for AUC(T) should close to 0.5. In our application, Causal Forests are not used for causal inference any more but for association analysis, where we generalised the effect from the coefficient of X0 without thinking its sign. So, we can also build a double machine learning system to measure the feature importance of X0 separately in XA and XB, and combine the SHAP importance values from two cohorts together, namely SHAP1 represents the importance of X0 to scaled Y1, SHAP2 represents the importance of X0 to scaled Y2.

SHAP1 x SHAP2 will measure the importance to both cohorts. Furthermore, it can be applied to connecting more than two cohorts to find underlying patterns.

### 5.2 Connecting PEAC and ORBIT to pain

We tried to explore the integrated data from RNAcare based on case study 1 at 3^rd^ granularity level. When processing with PEAC (0) and ORBIT (1), different from RNAcare, we treated pain VAS as a continuous variable (log1ped).

#### 5.2.1 Causal Forest

Figure 12-A shows the testing performance similar to before. 0.95 AUC for treatment model on test dataset means there exists a very serious difference between the two cohorts although for the same disease after batch correction. 0.226 MSE for test dataset. Figure 12-B shows SHAP feature importance ordered by the spread (range) of SHAP values after running XGBoost. Figure 12-C-F shows the important metagenes and its pathway analysis results. The grey boxes in the Figure represents the range.

**Figure 12.** Connect PEAC and ORBIT with pain via Causal Forests I. (A) Test dataset-Treatment model AUC curve; (B) Test dataset -Performance of outcome model; (C) SHAP importance plot based on spread; (D) SHAP dependence plot of metagene_3_o; (E) Pathway analysis of metagene_3_o; (F) Pathway analysis of metagene_4_o.

We can see from Figure 12-B that the model thinks metagene_3_o is the most important one that separates two cohorts but functions the same (if we ignore grey-box area in Figure 11-D) and interacts with metagene_4_o (synergetic effect). metagene_3_o is enriched in IFN, and metagene_4_o is enriched in BIOCARTA_NKCELLS_PATHWAY and BIOCARTA_CTL_PATHWAY which are involved in immune surveillance and immune regulation.

**Figure 11.**
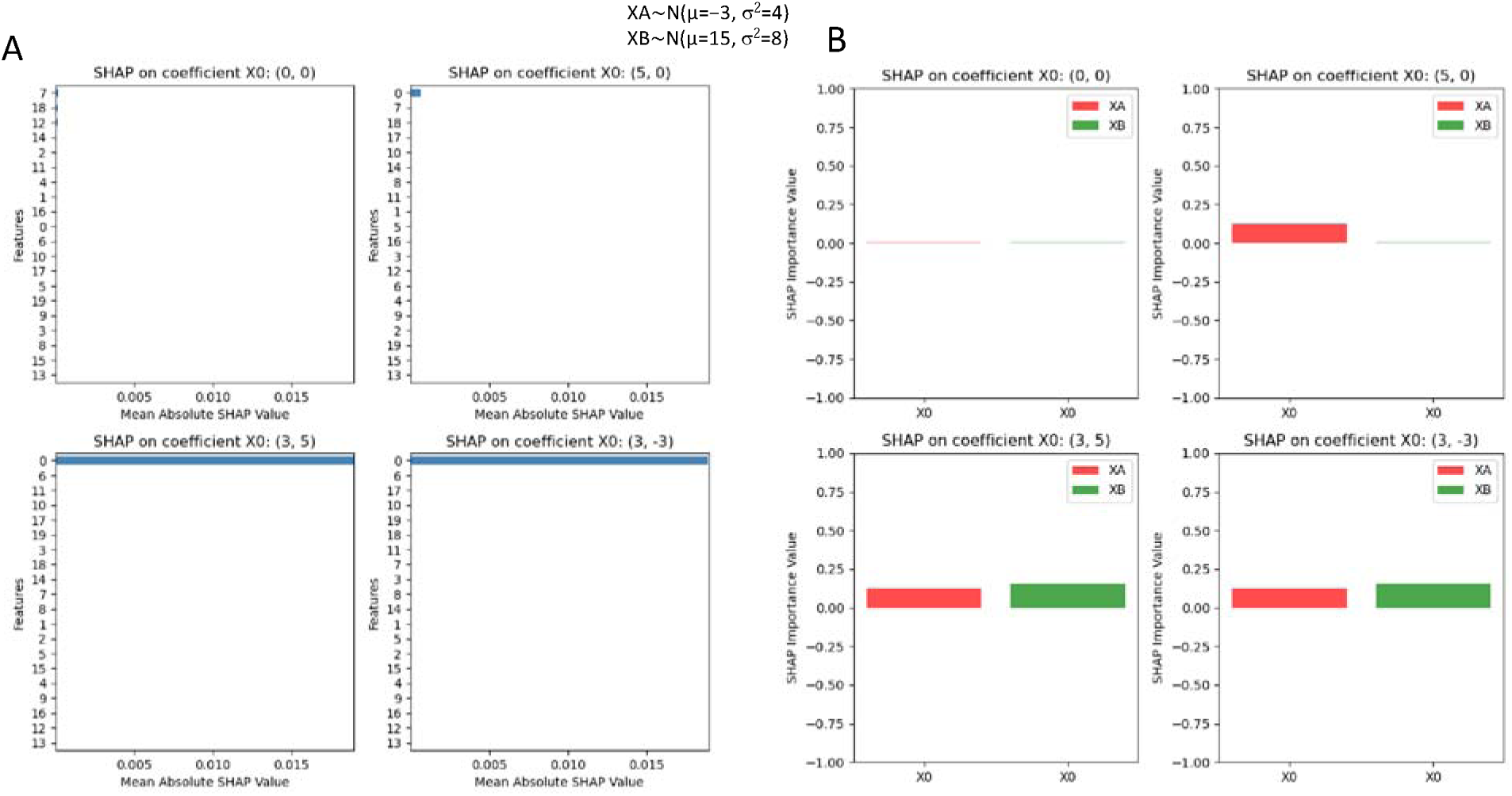
Connecting simulated metagenes to phenotype with different coefficients of X0 for each group. (A) combined SHAP importance value for coefficients of X0 in XA and XB; (B) Coefficients separate contribution to X0 on SHAP values.

Metagene_6_o doesn’t exhibit a balanced shape in Figure 12-B, so it is locally functional. We can see its average effect to PEAC is stronger than that to ORBIT if we do an average horizontal line at both sides after excluding the grey region. It is enriched in IL-1 related pathway (Figure 13-AB): Interleukin-1 receptor antagonist (IL-1Ra), specifically in the form of anakinra, has been studied as a treatment for RA. Clinical trials have demonstrated that anakinra can reduce the signs and symptoms of RA, with a significant percentage of patients achieving a 20% improvement in disease activity^55^, disease activity is highly positively related to pain which we talked before in RNAcare. It is also enriched in NFKB related pathway: Nuclear factor kappa-B (NF-κB) is a transcription factor that plays a key role in the development of rheumatoid arthritis (RA) pain. NF-κB is an inflammatory pathway that can lead to bone erosion, cartilage destruction, and joint damage^56^;

**Figure 13.** Connect PEAC and ORBIT with pain via Causal Forests II. (A) SHAP dependence plot of metagene_6_o; (B) Pathway analysis of metagene_6_o; (C) SHAP dependence plot of metagene_0_o; (D)Pathway analysis of metagene_0_o; (E) Pathway analysis of metagene_0_p;

We find L1CAM related pathway is enriched in metagene_0_o (Figure 13-C-E); In all, we consider that after batch correction at transcriptomic level, features with strong heterogeneity may not easily be detected by RNAcare.

#### 5.2.2 DML

We applied DML to the same case. Figure 14-A shows the importance based on our definition. Figure 14-B is pathways of metagene_0_p, which is overlapped with metagene_1_o (Figure 14-C), related to Non-sense mediated decay NMD and selenoamino acid metabolism. Metagene_0_o is enriched in IFN signaling, aligning with Figure 12-E. Non-sense mediated decay NMD is correlated with RHO GTPase cycle in Figure 3-A.

**Figure 14.** Connect PEAC and ORBIT with pain via DML. (A) Feature importance of SHAP values; (B) Pathway analysis of metagene_0_p; (C) Pathway analysis of metagene_1_o; (D) Pathway analysis of metagene_0_o; (E) Feature contribution for each cohort.

The issue of this method, is for the two machine learning methods, they can’t always use the same feature (maybe correlated one) for splitting.

### Case Study 6: Signatures connecting different tissues to pain via DML

To best demonstrate our platform, we tried to explore the possibility of common underlying mechanisms to pain between different tissues, which may stand for a potential crosstalk or migration between different tissues. We use PEAC for the exploration with two different tissues: whole blood (0-67 records) and synovium (1-53 records).

This time, we didn’t include other clinical features other than pain VAS, because there is an overlap of patients between different tissues. Other technique to solve this problem can be inferCNV to infer copy number variants^57^.

### 6.1 Causal Forests

Using the same method, we did log1p transformation for the pain VAS, and got high AUC for treatment model with 0.453 MSE for outcome model (Figure 15-A). In Figure 15-B, it shows the important metagenes influencing both tissues. Figure 15-C is the most one: metagene_3_wh, representing neutrophil degranulation and antimicrobial peptides with shared signatures CTSG, LTF, LCN2; Figure 15-D is the SHAP dependence plot for metagene_3_wh, we can see there is no metagene interacting with it. Similarly, we did all pathway analysis and SHAP dependence plots for all important metagenes, see Figure 15 and Figure 16. After that, we explained in the Table 6.

**Figure 15.** Signatures connecting different tissues to pain I. (A) Test dataset-Treatment model AUC curve; Test dataset -Performance of outcome model; (B) SHAP importance plot based on spread; (C) Pathway analysis of metagene_3_wh; (D) SHAP dependence plot of metagene_3_wh; (E) SHAP dependence plot of metagene_0_wh; (F) Pathway analysis of metagene_0_wh.

**Figure 16.** Signatures connecting different tissues to pain II. (A) Test dataset-Treatment model AUC curve; Test dataset -Performance of outcome model; (B) SHAP importance plot based on spread; (C) Pathway analysis of metagene_3_wh; (D) SHAP dependence plot of metagene_3_wh; (E) SHAP dependence plot of metagene_0_wh; (F) Pathway analysis of metagene_0_wh.

**Table 6.** Explaination of signatures of different tissues to pain.

| Metagenes | Explanation | Interesting signatures |
| --- | --- | --- |
| metagene_3_wh | We think **CTSG, LTF, LCN2** imply blood-synovial crosstalk, because they are released by neutrophils, which migrate from blood to synovium in RA | **CTSG, LTF, LCN2** |
| metagene_0_wh | Enriched in **IFN signalling** pathway with signature CXCL10<sup>58</sup>. | **CXCL10, CCL2** |
| metagene_1_sy | **PI3K/Akt signaling pathway plays a crucial role in the pathogenesis of rheumatoid arthritis (RA) and facilitates cross-talk between the blood and synovial tissues.** (1) The p110δ isoform of PI3K is predominantly expressed in leukocytes and plays a crucial role in immune cell functions. Activation of p110δ in T cells, B cells, and neutrophils enhances their proliferation, survival, and migration. This activation facilitates the movement of these cells from the bloodstream into inflamed tissues, such as the synovium in RA<sup>59</sup>. (2) PI3K signaling is integral to the chemotactic responses of immune cells. Upon activation by chemokines, PI3K generates lipid second messengers that recruit downstream effectors, leading to cytoskeletal rearrangements and directed cell movement. This mechanism enables blood-derived immune cells to migrate towards inflammatory signals present in the synovium<sup>60</sup>. (3) PI3K and S1PR1-Mediated Migration:<br>The S1P receptor 1 (S1PR1), coupled with PI3K signaling, regulates lymphocyte egress from lymphoid organs and their trafficking to peripheral tissues. PI3K activation downstream of S1PR1 engagement promotes lymphocyte migration, potentially contributing to their accumulation in the synovium during RA<sup>61</sup> | **ERBB3, FGFR3** |
| metagene_5_wh | Neutrophil extracellular traps (NETs) activate the **intrinsic pathway** of coagulation by exposing negatively charged surfaces, primarily through their DNA component, which triggers the activation of factor XII (Hageman factor), initiating a cascade of reactions leading to thrombin generation and clot formation<sup>62</sup>; In the context of synovial inflammation, fibrin clots indicate the presence of an inflammatory process within a joint, as the formation of fibrin clots within synovial fluid is directly related to the level of inflammation, with increased fibrin deposition occurring in conditions like rheumatoid arthritis, where the synovial membrane becomes inflamed and leaks fibrinogen, which then clots into fibrin within the joint space; essentially, the more fibrin clots present, the more severe the synovial inflammation is considered to be<sup>63</sup>. | **PROS1, GP1BA** |
| metagene_1_wh | **CCL2, also known as MCP-1, plays a significant role in the crosstalk between blood and synovia fluid in RA. CCL2 acts as a bridge between systemic inflammation (blood) and local joint inflammation (synovial fluid)** (1) Recruitment of Monocytes to Inflamed Joints<sup>64</sup>; (2) Correlation with Disease Activity<sup>65</sup>; (3) Therapeutic Targeting of the CCL2/CCR2 Axis<sup>66</sup>;<br>**IL-18 Induces CCL2:** IL-18 stimulates macrophages, leading to the activation of the PI3K/Akt and MEK/ERK1/2 signaling pathways. This activation results in the production of CCL2, which plays a crucial role in recruiting leukocytes to inflammatory sites<sup>67</sup>; In RA, elevated levels of IL-18 in the synovial fluid can induce the production of CCL2. This chemokine attracts monocytes from the bloodstream into the synovium, where they differentiate into macrophages and perpetuate inflammation, contributing to joint damage<sup>68</sup>. | **CCL2, JUN** |
| metagene_6_wh | In the context of inflammatory bowel disease (IBD), **TNFSF12** has been observed to induce an inflammatory transcriptional program in colonic fibroblasts, leading to the upregulation of adhesion molecules such as ICAM-1 and VCAM-1, and increased secretion of the chemokine CCL2. These changes facilitate the recruitment and activation of monocytes, suggesting a mechanism by which TWEAK-exposed fibroblasts can promote immune cell infiltration into tissues<sup>69</sup>. **NME1**: included in ARF6 downstream pathway, Arf6 plays a significant role in regulating the migration of macrophages. Research indicates that Arf6 influences various cellular processes essential for macrophage movement and ARF6 further assists actin polymerization and remodeling by the directed transport of lipid rafts, **Rho GTPases**, and Rho signaling components<sup>70</sup>. | **TNFSF12, NME1** |

### 6.2 DML

Figure 17 shows DML applied on the same data. We didn’t find pathway result for metagene_1_wh, but metagene_2_wh, metagene_0_sy, metagene_4_wh and metagene_3_wh are corresponding to Figure 15-F, B, D, Figure 14-C.

**Figure 17.** Connect tissues for PEAC with pain via DML. (A) Feature importance of SHAP values; (B) Pathway analysis of metagene_2_wh; (C) Pathway analysis of metagene_0_sy; (D) Pathway analysis of metagene_4_wh; (E) Pathway analysis of metagene_3_wh; (F) Feature contribution for each cohort.

**Conclusion: We found after performing a zero-centered procedure**^71^**, batch correction is no needed, because Causal Forests and DML compute the contributions from features separately in their own groups then unify them under the same scale of the clinical phenotype. We found CCL2, PROS1 plays a critical cross talk between two tissues.**

### Case Study 7: Signatures connecting transcriptomics and proteomics to pain via DML

Based on what we discussed so far, we therefore assume whether RNAcompare can recognise relationship/crosstalk for omics datasets using different techniques. We introduced RA-MAP, including both transcriptomic (wh) and proteomic (pro) data, and expect a cross talk in terms of fatigue (see Appendix 4) and pain (log1p-ed). The combination of two levels greatly reduce the number of features for analysis.

### 7.1 Causal Forests

Figure 18 shows the result. Figure 18-A is the treatment model AUC and outcome model MSE 0.296 based on test data. Figure 18-B is the SHAP importance plot. Pathway analysis and the corresponding dependence plots are Figure 18-C-F. Table 7 is the result explanation.

**Figure 18.** Signatures connecting transcriptomics and proteomics to pain I. (A) Test dataset-Treatment model AUC curve; Test dataset -Performance of outcome model; (B) SHAP importance plot based on spread; (C) Pathway analysis of metagene_3_wh; (D) SHAP dependence plot of metagene_3_wh; (E) SHAP dependence plot of metagene_0_wh; (F) Pathway analysis of metagene_0_wh.

**Figure 19.** Signatures connecting transcriptomics and proteomics to pain II. (A)SHAP dependence plot of metagene_6_wh; (B) Pathway analysis of metagene_6_wh; (C) SHAP dependence plot of metagene_3_pr; (D) Pathway analysis of metagene_3_pr; (E) SHAP dependence plot of metagene_4_wh; (F) Pathway analysis of metagene_4_wh; (G) Pathway analysis of metagene_1_wh.

**Table 7.**
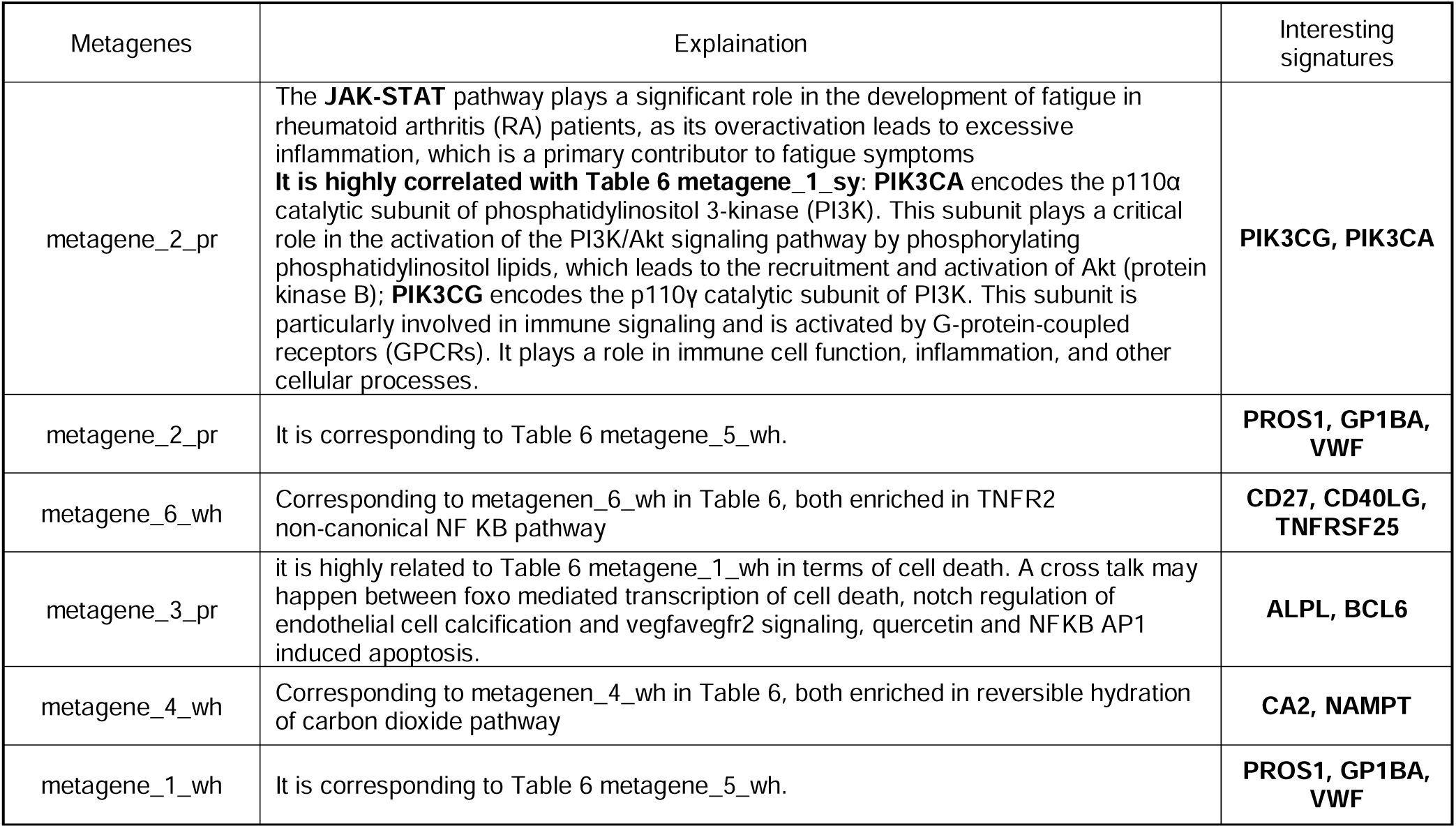
Explaination of signatures of multi-omics to pain.

### 7.2 DML

We then applied DML to the same study. Figure 20 shows the results. Interestingly, we found signature PIK3CG functions importantly in both DML and Causal Forests model. Then we assume PIK3CG plays a crosstalk between transcriptomic and proteomic levels.

**Figure 20.** Connect omics for RA-MAP with pain via DML. (A) Feature importance of SHAP values; (B) Pathway analysis of metagene_4_tr; (C) Pathway analysis of metagene_0_pr; (D) Pathway analysis of metagene_2_tr; (E) Pathway analysis of metagene_6_tr; (F) Feature contribution for each cohort.

We seem to be also able to assume a unified model in RA via blood-synovium crosstalk (CCL2) and the blood transcriptomic-proteomic crosstalk (PIK3CG):

1. CCL2, a chemokine elevated in RA blood and synovium, recruits monocytes, macrophages, and T cells from the bloodstream into inflamed synovium via its receptor CCR2; PIK3CG drives PI3Kγ-AKT-mTOR signalling (proteomically), promoting immune cell activation, cytokine production and chemokine release.
2. Integrated Model Connecting CCL2 and PIK3CG:

1 **PIK3CG Transcriptomic Activation in Blood**: Systemic inflammation upregulates PIK3CG mRNA in circulating immune cells such as monocytes and T cells, causing increased PIK3CG protein, which amplifies PI3Kγ signalling. This leads to AKT/mTOR activation and NF-kB-mediated transcription of pro-inflammatory genes.
2 **PIK3CG-Driven CCL2 production in blood**: PIK3CG signalling enhances CCL2 transcription in blood and increased CCL2 protein is secreted into circulation^72,73^.
3 **CCL2 Mediates Blood-to-Synovium Crosstalk**: circulating CCL2 binds to CCR2 on synovial endothelial cells, promoting vascular permeability; CCL2 recruits CCR2+ monocytes and T cells from blood into synovium.
4 Synovial Inflammation Feeds Back to Blood.

**Conclusion: by combining case study 6,7, we successfully connected underlying mechanisms between tissues and omics data, finding significant pathways/signatures across different batches with their signatures.**

### Case Study 8: Connecting different diseases to disease score via DML

To best demonstrate our platform, we also tried to explore the possibility of common underlying mechanisms connecting Heart Failure (HF-0) and RA(1) to disease severity via Causal Forests.

Heart failure (HF) has been recognized as global pandemic with a high rate of associated hospitalization, morbidity and mortality. Although numerous advances have been made, its representative molecular signatures remain largely unknown, especially the role of genes in HF progression. The aim of the present prospective follow-up study was to reveal potential biomarkers associated with the progression of heart failure.

Since we had pre-knowledge of pain related pathways in RA from RNAcare, based on which, we used ORBIT + PEAC (normalised and log1p CPMed, 139+49 records) and GSE135055 (FKPM, 18) to explore the similarity between two diseases.

We established a Causal Forests with AUC 1 and 0.245 MSE on test data (Figure 21-A). Figure 21-B shows the SHAP importance features where we can see the metagene_1_hf is the most significant signature contributing to the difference of disease activity between HF and RA. From Figure 21-C, we see the model thinks with the age increases, the severity of having HF increases. In our HF dataset, we only have patients aged between 19-60. Therefore, Causal Forests used the combined dataset to predict the elder patients which we need to be cautious when interpretation. We made a deeper research for metagene_1_hf. Figure 21-D is its pathway analysis. Figure 21-E is its SHAP dependence plot. We can see this metagene is enriched in response of EIF2AK4/GCN2 to amino acid deficiency, which we think is correlated to both disease in terms of severity^74^. It is also enriched in nonsense mediated decay NMD^75^;

**Figure 21.** Connect PEAC, ORBIT and GSE135055 with DAS via Causal Forests I. (A) Test dataset-Treatment model AUC curve; Test dataset -Performance of outcome model; (B) SHAP importance plot based on spread; (C) SHAP dependence plot for age; (D) Pathway analysis of metagene_1_hf; (E) SHAP dependence plot of metagene_1_hf; (F) SHAP dependence plot of metagene_5_op; (G) Pathway analysis of metagene_5_op.

**Figure 22.** Connect PEAC, ORBIT and GSE135055 with DAS via Causal Forests II. (A) SHAP dependence plot of metagene_1_op; (B) Pathway analysis of metagene_1_op; (C) Pathway analysis of metagene_5_hf; (D) SHAP dependence plot of metagene_5_hf; (E)Pathway analysis of metagene_6_hf; (F) SHAP dependence plot of metagene_6_hf;

For metagene_5_op, it is enriched in neutrophil degranulation^76^ and nuclear receptor metapathway^77,78^.

For metagene_1_op, it is also enriched in MMP9 which we analysed before the importance in RA. Studies also discuss the importance in HF^79–81^. Notably, we note COL6A2 and MMP9 for assembly of collagen fibrils and other multimeric structure. This is highly related to COL1A1^17^. Studies suggest that COL6A2 stabilizes and organizes collagen fibrils (like COL1A1), enhancing tissue strength^82,83^.

For metagene_5_hf, it is enriched in selenoamind acid metabolism^84,85^. Interestingly, it is also enriched in influenza infection, which means both diseases are potentially related to type I IFN and even RHO GTPase pathway^86,87^.

For metagene_6_hf, COX7B is the shared signatures between both^88,89^.

**Conclusion: via Causal Forests/DML, we don’t need to consider batch correction anymore, because the importance from features to the clinical phenotype is calculated separately, unified with the same scale across diseases. The issue changes to whether there exists a clinical measure bridging two different cohorts.**

**Conclusion: by comparing two different diseases via Causal Forests, we find the similar hidden pathways and their interactions toward disease severity.**

### Case Study 9: Connecting different techniques to pain via DML

In this case, we combined log1p CPMed RNA-seq data (0-ORBIT+PEAC) and Microarray data (1-RA-MAP), based on what we discussed so far about the prior knowledge to pain.

Notably, DAS score in RA-MAP is standardised. So, when combine two datasets, DAS score in ORBIT+PEAC needs to be standardised as well.

Figure 23 and Figure 24 shows the result. Figure 23-A is the treatment model AUC and outcome model MSE 0.189 based on test data. Figure 23-B is the SHAP importance plot. Pathway analysis and the corresponding dependence plots are Figure 23-C-G and Figure 24. Table 8 is the result explanation.

**Figure 23.** Connect different techniques to pain via Causal Forests I. (A) Test dataset-Treatment model AUC curve; Test dataset -Performance of outcome model; (B) SHAP importance plot based on spread; (C) SHAP dependence plot for DAS score; (D) Pathway analysis of metagene_3_seq; (E) SHAP dependence plot of metagene_4_seq; (F) SHAP dependence plot of metagene_4_seq;

**Figure 24.** Connect different techniques to pain via Causal Forests II. (A) SHAP dependence plot of metagene_5_seq; (B) Pathway analysis of metagene_5_seq; (C) SHAP dependence plot of age; (D)Pathway analysis of metagene_0_seq.

**Table 8.**
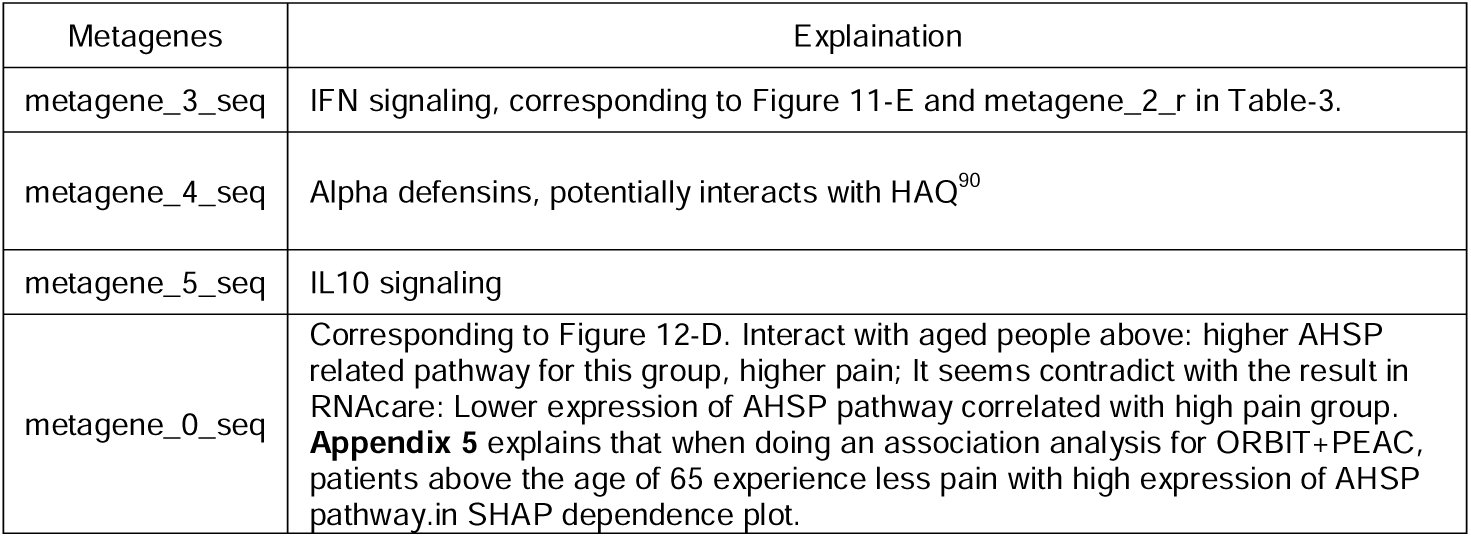
Explaination of metagenes across different techniques to pain.

**Conclusion: By comparing expression data with bulk RNA-seq and microarray, we find common pathways influencing pain at transcriptomic level, namely IFN signalling, AHSP related pathway, which was the same result got from RNAcare.**

## CONCLUSION

RNAcompare has been designed to enable comparison across different data types, including tissues, omics levels, diseases by offering multiple machine learning algorithms for association inference and suggests a new way to bypass batch correction with the help of clinical data. We find signatures with correlated pathways influencing generalised drug response, hidden patterns for crosstalk between tissues and omics levels, similar mechanisms between RA and HF.

Apart from RNAcare, current tools don’t have the ability to incorporate clinical data with the omics data. One of the reasons might be that to date, clinical parameters are not highly focused when researching omics data. However, we postulate that just by having those parameters, more integrative analysis can be performed, allowing to leverage the existing data in more depth. At the same time, our case studies show potential avenues of analysis, albeit some of the results did not get fully proved by published papers and experiments.

Anyhow, we hope that due to the AI wave, more data will become available and RNAcompare will facilitate user without a bioinformatics background to explore data and generate new hypotheses.

## Availability and requirements

Project name: RNAcompare

Project home page: https://github.com/tangmingcan/RNAcompare

Operating system(s): Platform independent, tested on Linux (Ubuntu)

Compatible browsers: Firefox/Chrome

Programming language: Python, JavaScript

Other requirements: Python >= 3.11, Django >= 4.2, Nginx(optional)

Any restrictions to use by non-academics: licence needed

## List of abbreviations

FAIR: Findable, Accessible, Interoperable, and Reusable
DEG: Differential Expression Gene
IMID: Immune-mediated inflammatory diseases
PEAC: Pathobiology of Early Arthritis Cohort
ORBIT: Optimal management of rheumatoid arthritis patients who Require Biological Treatment
CPM: Count Per Million
VAS: Visual Analogue Scale
DAS28: Disease Activity Score using 28 joint counts
PCA: Principal Component Analysis
t-SNE: t-Distributed Stochastic Neighbour Embedding
UMAP: Uniform Manifold Approximation and Projection
REST: Representational State Transfer
CSR: Class Switch Recombination
ESR: Erythrocyte Sedimentation Rate
HYNA: New York Heart Association
HTE: heterogeneous treatment effects

## Declarations

- Ethics approval and consent to participate

For the ORBIT data, all participants provided written, informed consent. The study protocol ORBIT was approved by the West of Scotland Research Ethics Committee on 3/11/2009 (REC reference number: 09/S0703/109/ EudraCT number: 2009-011268-13)

For other studies, data were published, see PEAC ^15^/ RA-MAP ^15^.

### Consent for publication

We have consent for publication.

### Availability of data and materials

The code of can be found https://github.com/tangmingcan/RNAcompare.

The RNA-Seq data from ORBIT can be found under the accession numbers Pending waiting for the publication of RNAcare, which is currently under review by BMC Medicine.

### Authors’ contributions

MT – Construction of tool. Data analysis. Writing of first draft. TS – technical support of implementation

## Acknowledgements

We thank Senior Lecturer Tim Storer at computing science, University of Glassgow, Professor James Pan and Jan Chow from the Business Analytics Department at the National University of Singapore for their helps to this paper.

## Appendix 1

In this section, we listed all the clinical features for datasets used in the paper.

**Appendix 1-Table 1.**
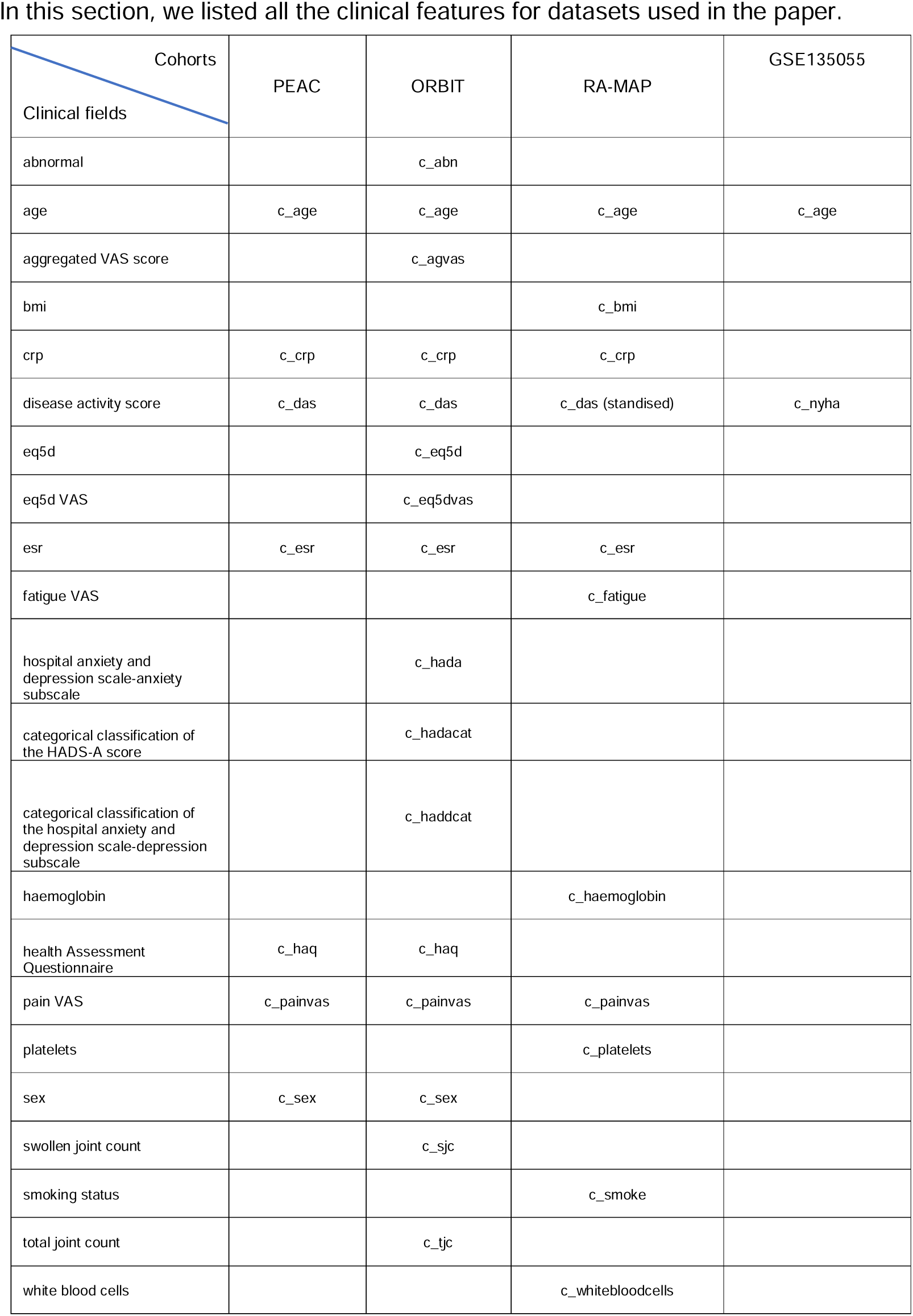
Clinical fields for datasets used in the paper.

## Appendix 2

By using the 1^st^ level granularity data, we reran the model, and got the similar components corresponding to the ones of interest in case study 1, see Figure 1. 0 is Rituximab responders, and 1 is anti-TNF non-responders. From Figure 1-D, we know that RHO GTPase interacts with influenza infection, but RHO GTPase takes the dominance, which is positive to anti-TNF non-response.

**Appendix 2- Figure 1.** 1st level data analysis. (A) SHAP value importance plot; (2) Pathway analysis of metagene_3_Rr; (3) Pathway analysis of metagene_6_TNFn; (4) SHAP dependence plot.

## Appendix 3

We found an issue when integrating ORA data with LightGBM. As it shows in Figure 1 (0-non-response, 1-response), the result is totally different compared with one in case study 4: with the increase of age, older patients will be more likely to drug non-response.

**Appendix3- Figure 1.** SHAP dependence plot for LightGBM on ORA data

## Appendix 4 Signatures connecting omics to fatigue

Figure 1 shows the results for transcriptomics (wh) and proteomics (sy). Figure 1-A is the treatment model ROC curve and outcome fitting with an MSE: 1.677 for test data. Figure 1-B is the SHAP importance plot for metagenes. We analysed the result in Table 1. Unlike RNAcare, since we exclude other clinical parameters, we can see that most important metagenes are first related to inflammation.

**Appendix4- Table 1.**
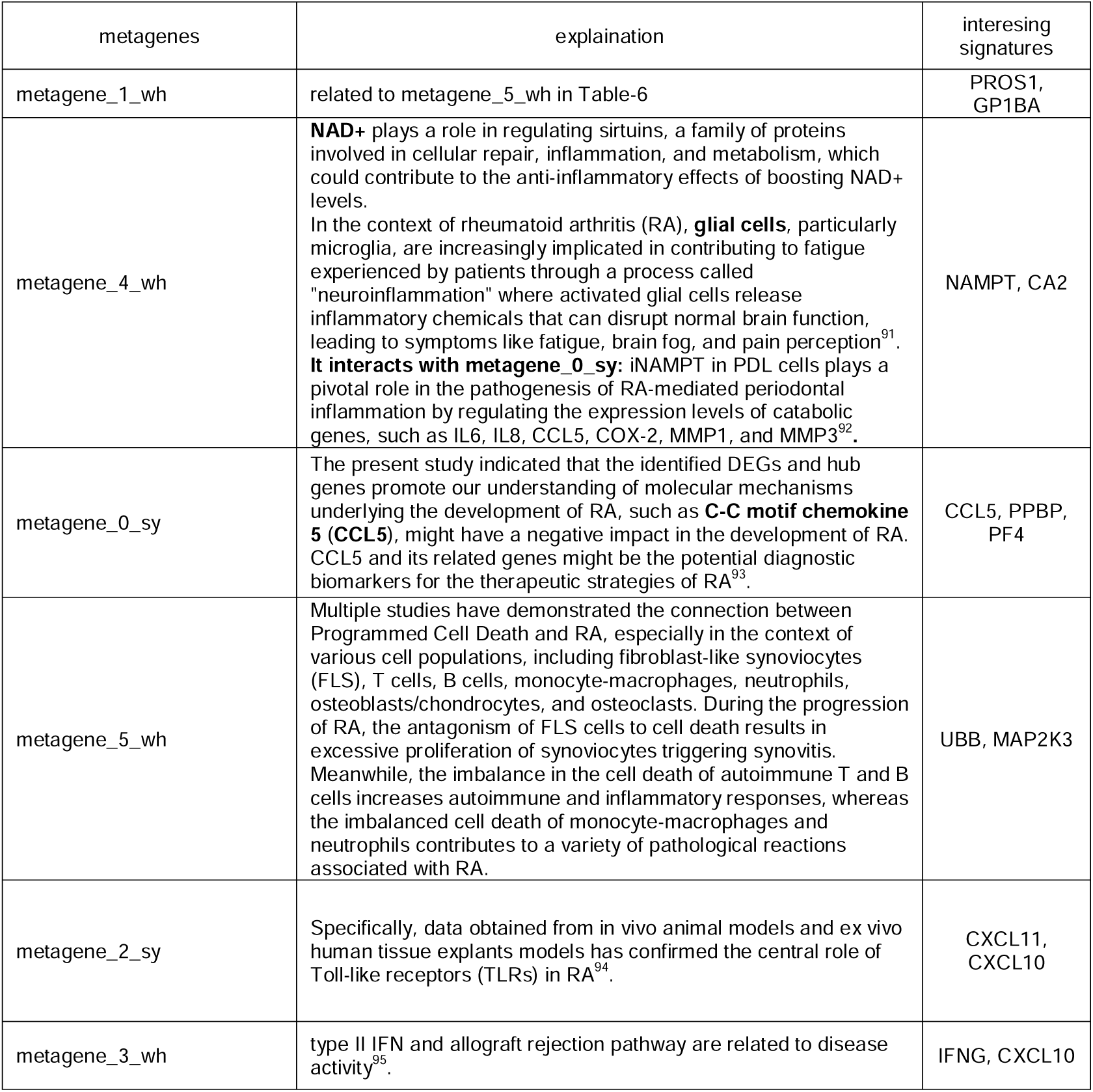
Signatures connecting omics to fatigue.

**Appendix 4-Figure 1.** Signatures connecting transcriptomics and proteomics to fatigue I. (A) Test dataset-Treatment model AUC curve; Test dataset -Performance of outcome model; (B) SHAP importance plot based on spread; (C) Pathway analysis of metagene_1_wh; (D) SHAP dependence plot of metagene_1_wh; (E) SHAP dependence plot of metagene_4_wh; (F) Pathway analysis of metagene_4_wh; (G) Pathway analysis of metagene_0_sy.

**Appendix 4- Figure 2.** Signatures connecting transcriptomics and proteomics to fatigue II. (A)SHAP dependence plot of metagene_5_wh; (B) Pathway analysis of metagene_5_wh; (C) SHAP dependence plot of metagene_2_sy; (D) Pathway analysis of metagene_2_sy; (E) SHAP dependence plot of metagene_3_wh; (F) SHAP dependence plot of metagene_3_wh; (G) Pathway analysis of metagene_2_wh; (H) Pathway analysis of metagene_2_wh;

## Appendix 5

As it shows in Figure 1, patients above the age of 70 exhibit an obvious trend that those with higher expression of metagene_0_o have more pain. Compared to the result from RNAcare, one explanation is when we grouped the patients into low and high pain group, patients with lower expression of metagene_0_o were more likely put into high pain group regardless of their ages, which is approve by Figure 1-C, showing the result of the distribution of the two groups based on metagene_0_o.

**Appendix5-Figure 1.** The interaction of age and metagene_0_o to pain. (A)SHAP dependence plot; (B) Pathway analysis of metagene_0_o. (C) metagene_0_o distribution in the two pain-related groups.

