## Supplementary material for "RNAcompare: Integrating machine learning algorithms to unveil the similarities of phenotypes based on patients’ clinical, multi-omics using Rheumatoid Arthritis and Heart Failure as Case Studies": supp figure

### Slide 1
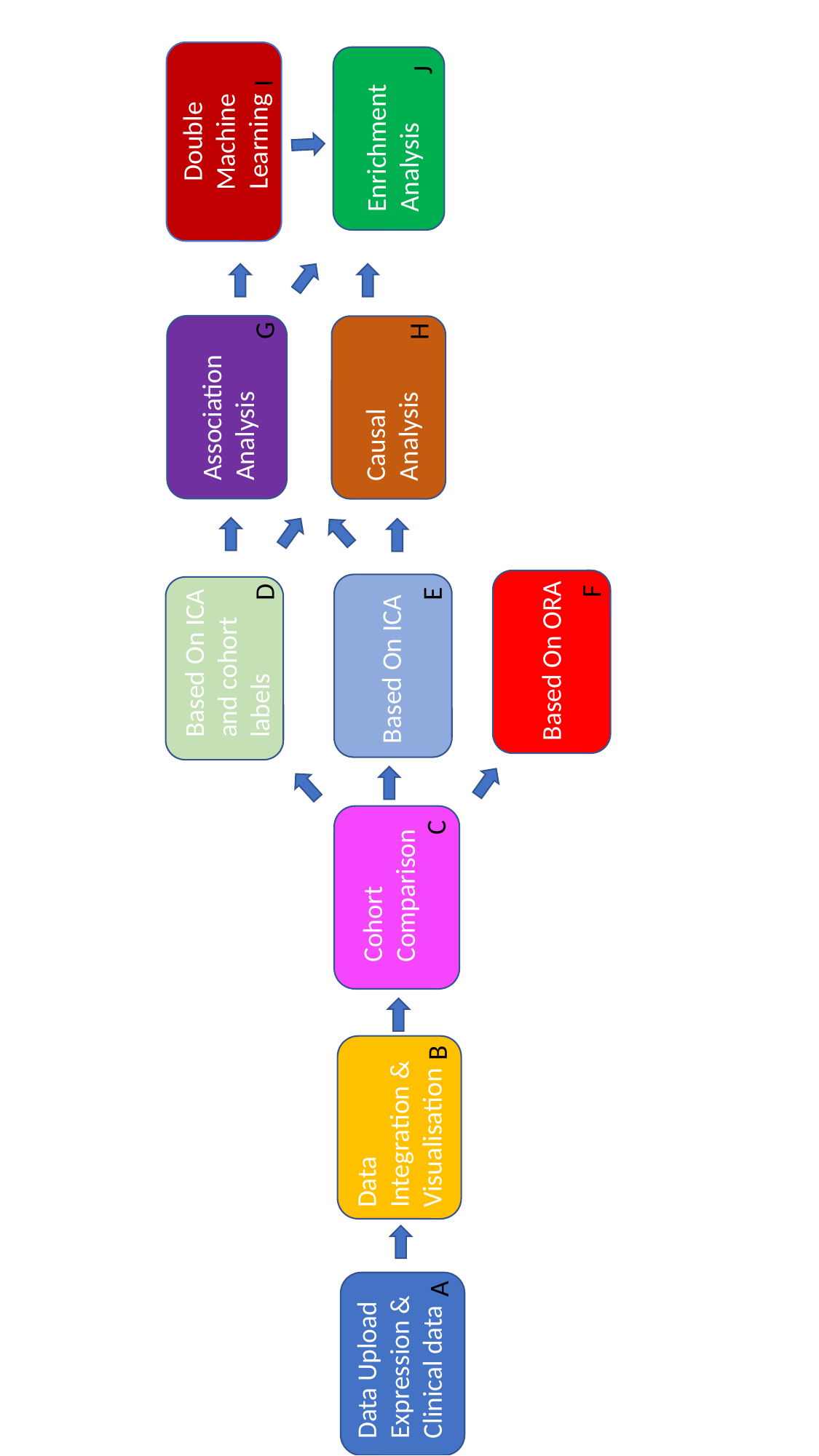

Based On ICA and cohort labels
Double Machine Learning
Association Analysis
I
G
D
Causal Analysis
Enrichment Analysis
Cohort Comparison
Based On ICA
Data Integration & Visualisation
Data Upload
Expression & Clinical data
H
J
E
C
B
A
Based On ORA
F

### Slide 2
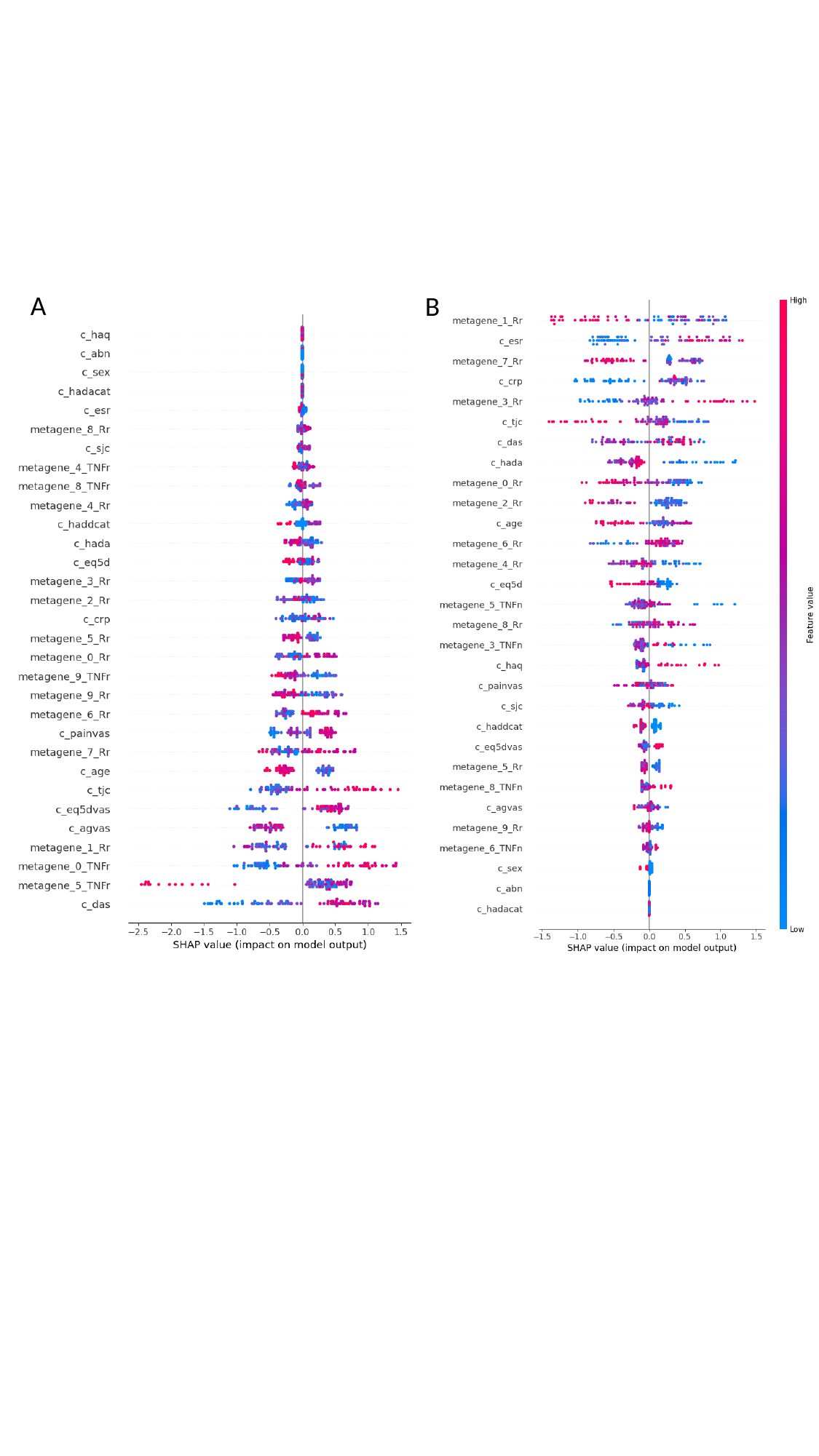

A
B

### Slide 3
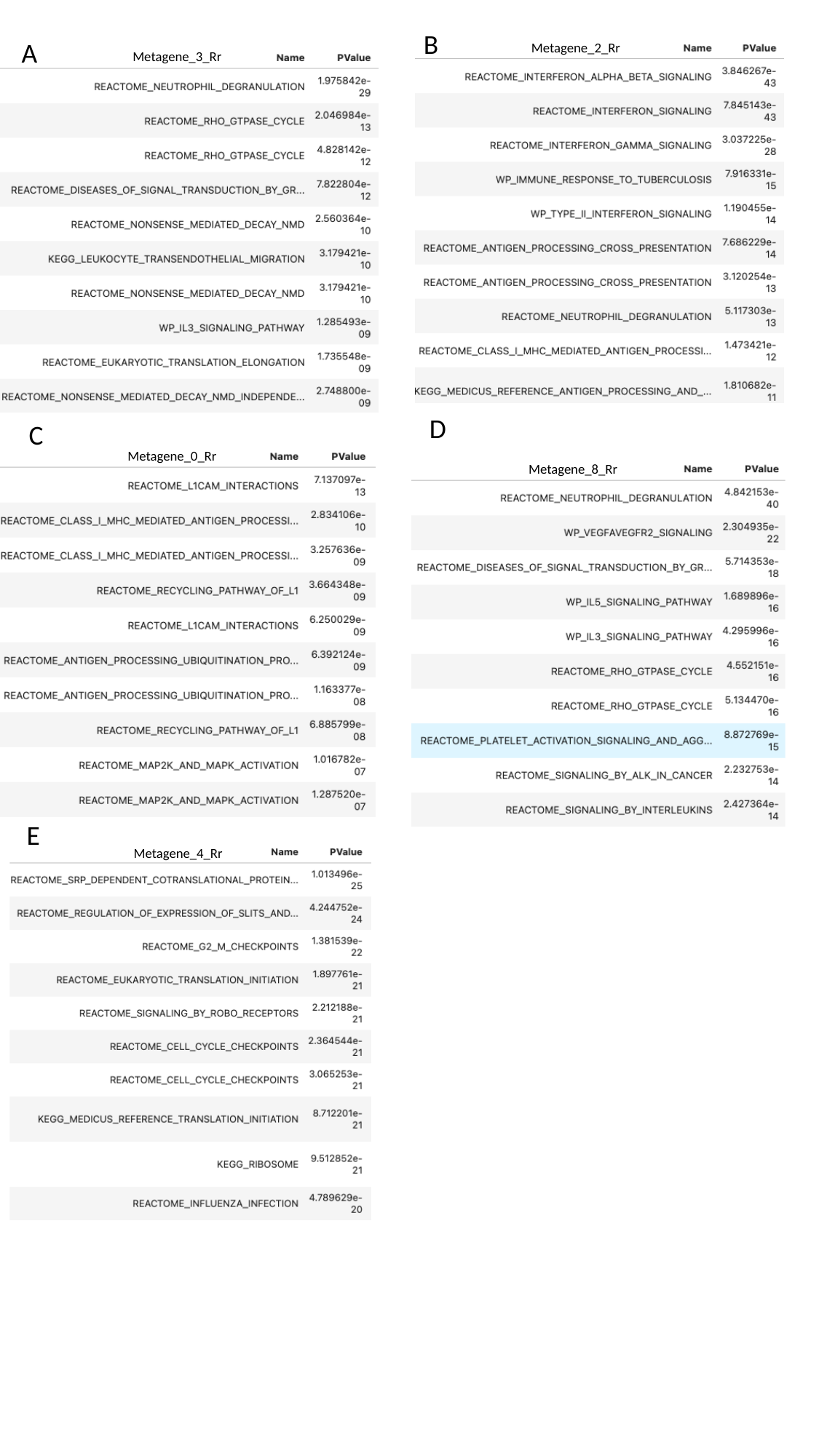

B
A
Metagene_2_Rr
Metagene_3_Rr
D
C
Metagene_0_Rr
Metagene_8_Rr
E
Metagene_4_Rr

### Slide 4
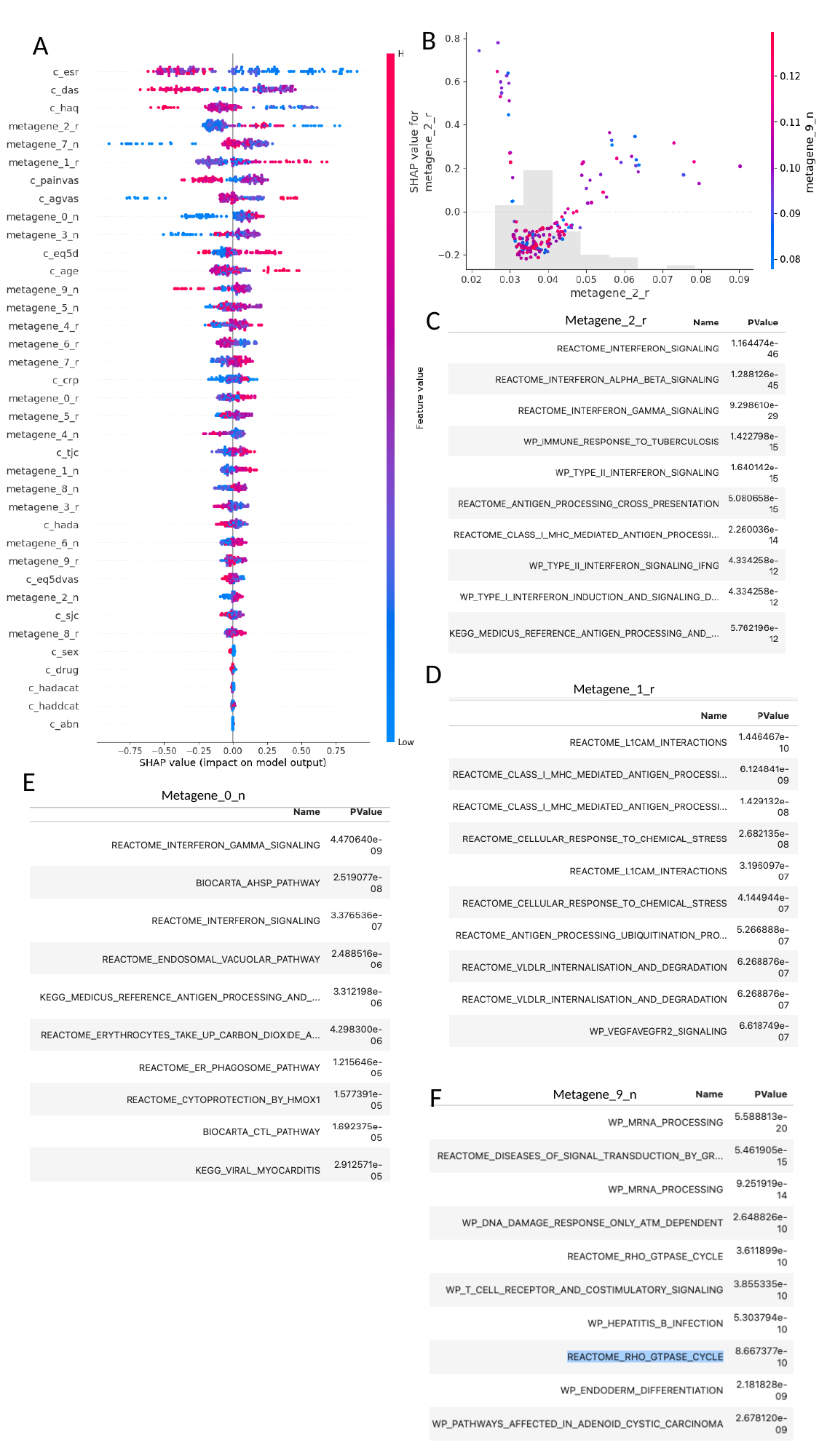

B
A
C
Metagene_2_r
D
Metagene_1_r
E
Metagene_0_n
F
Metagene_9_n

### Slide 5
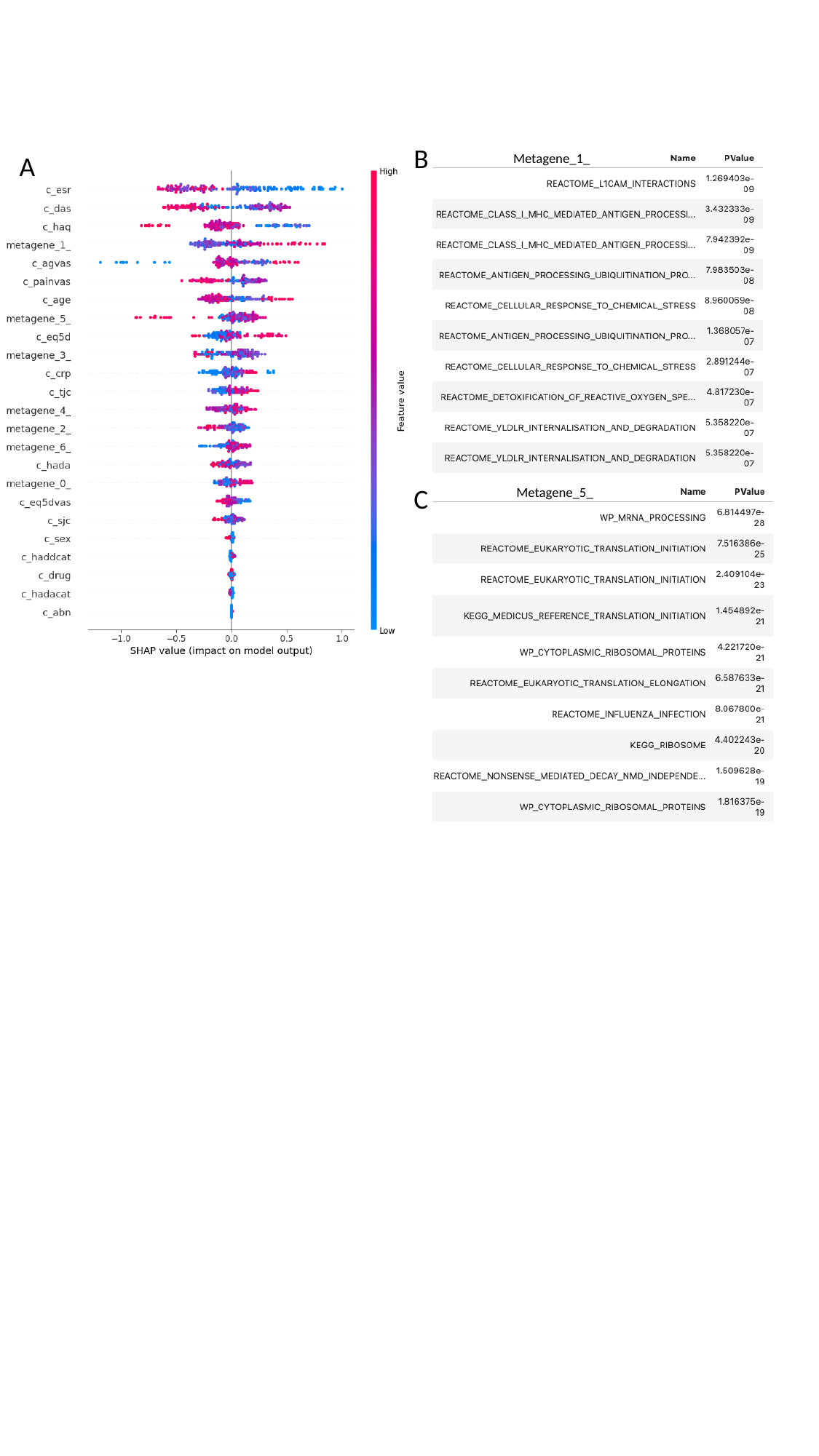

B
A
Metagene_1_
C
Metagene_5_

### Slide 6
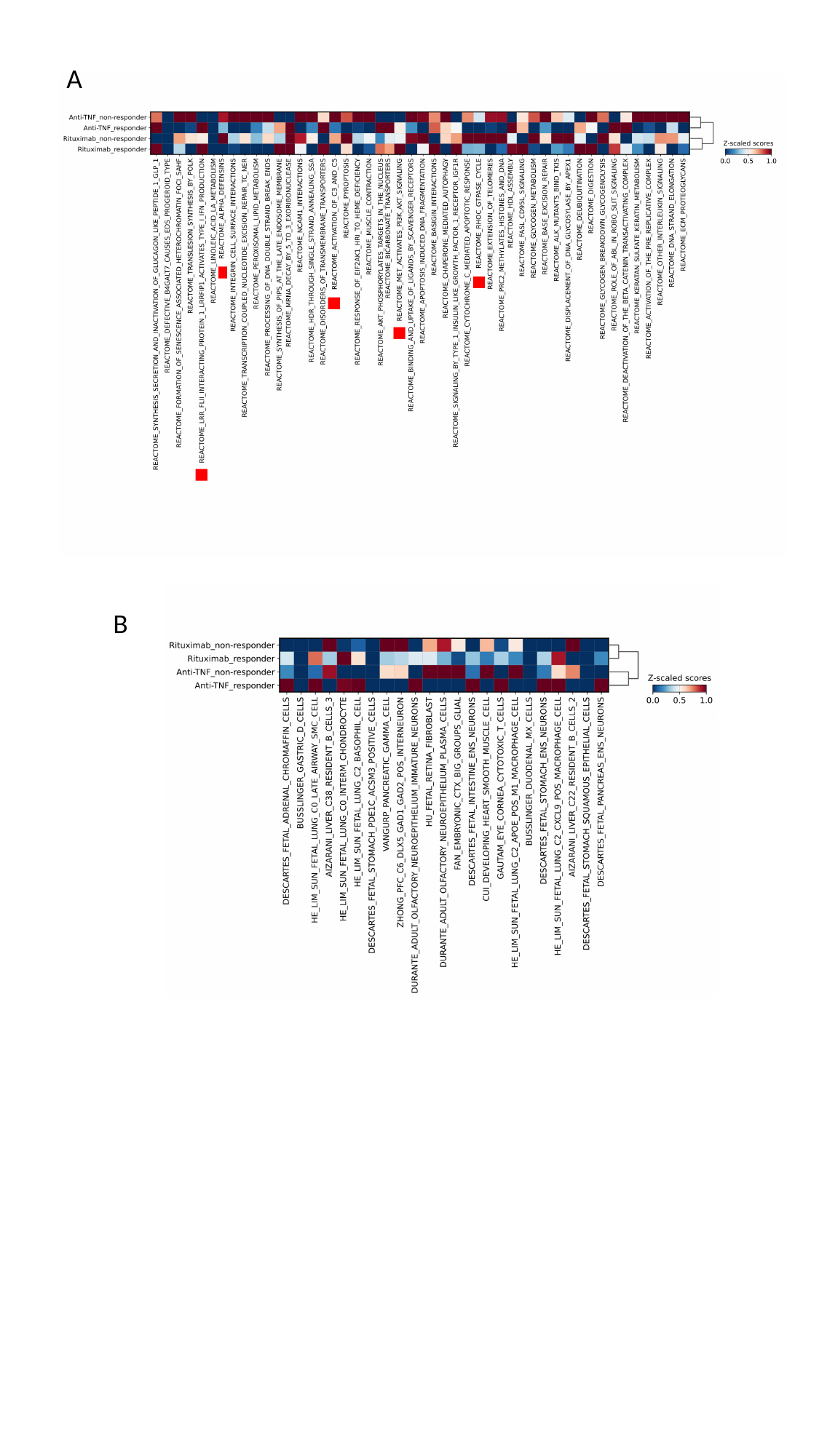

A
B

### Slide 7
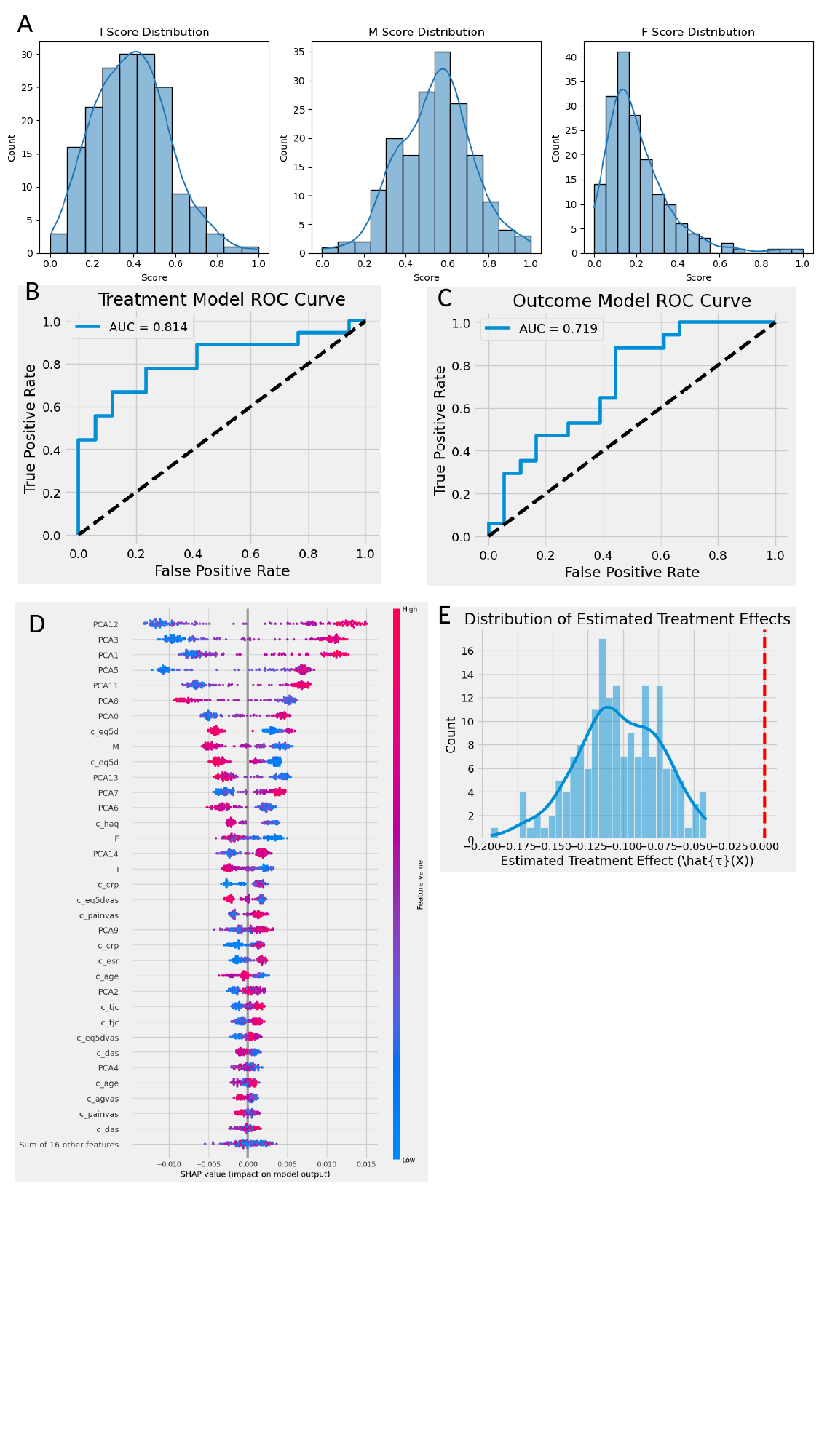

A
B
C
E
D

### Slide 8
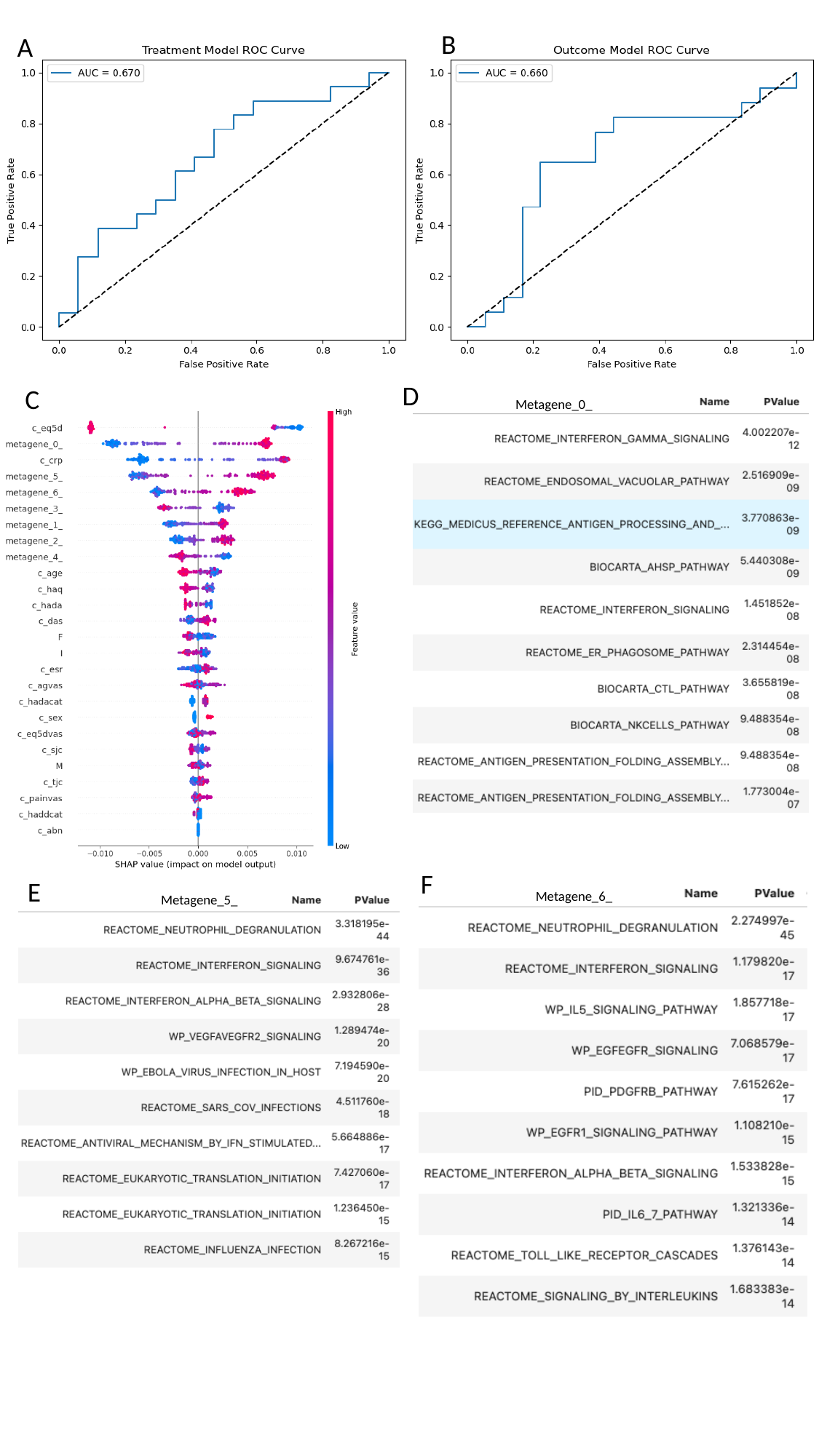

B
A
D
C
Metagene_0_
F
E
Metagene_6_
Metagene_5_

### Slide 9
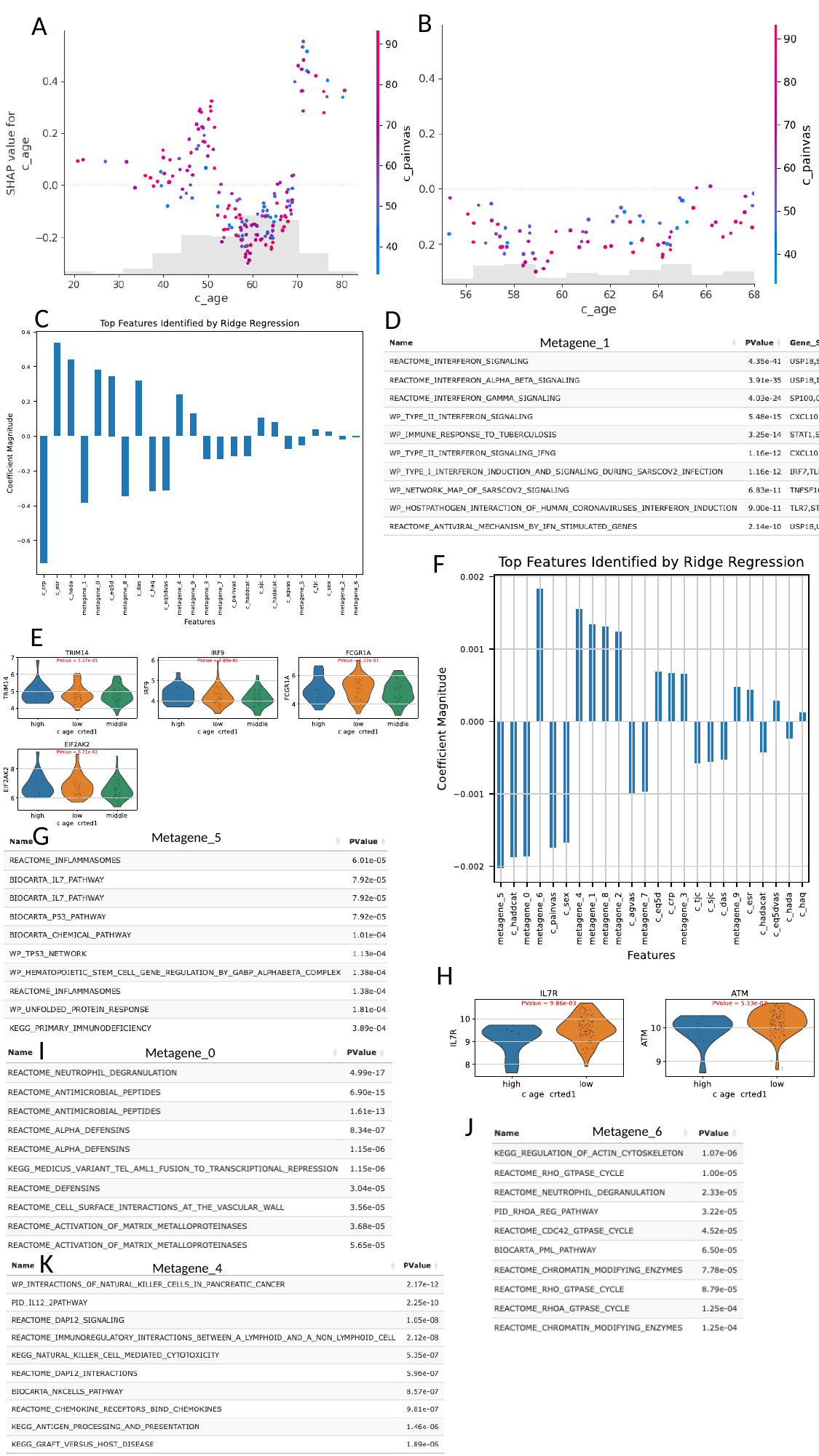

B
A
C
D
Metagene_1
F
E
G
Metagene_5
H
I
Metagene_0
J
Metagene_6
K
Metagene_4

### Slide 10
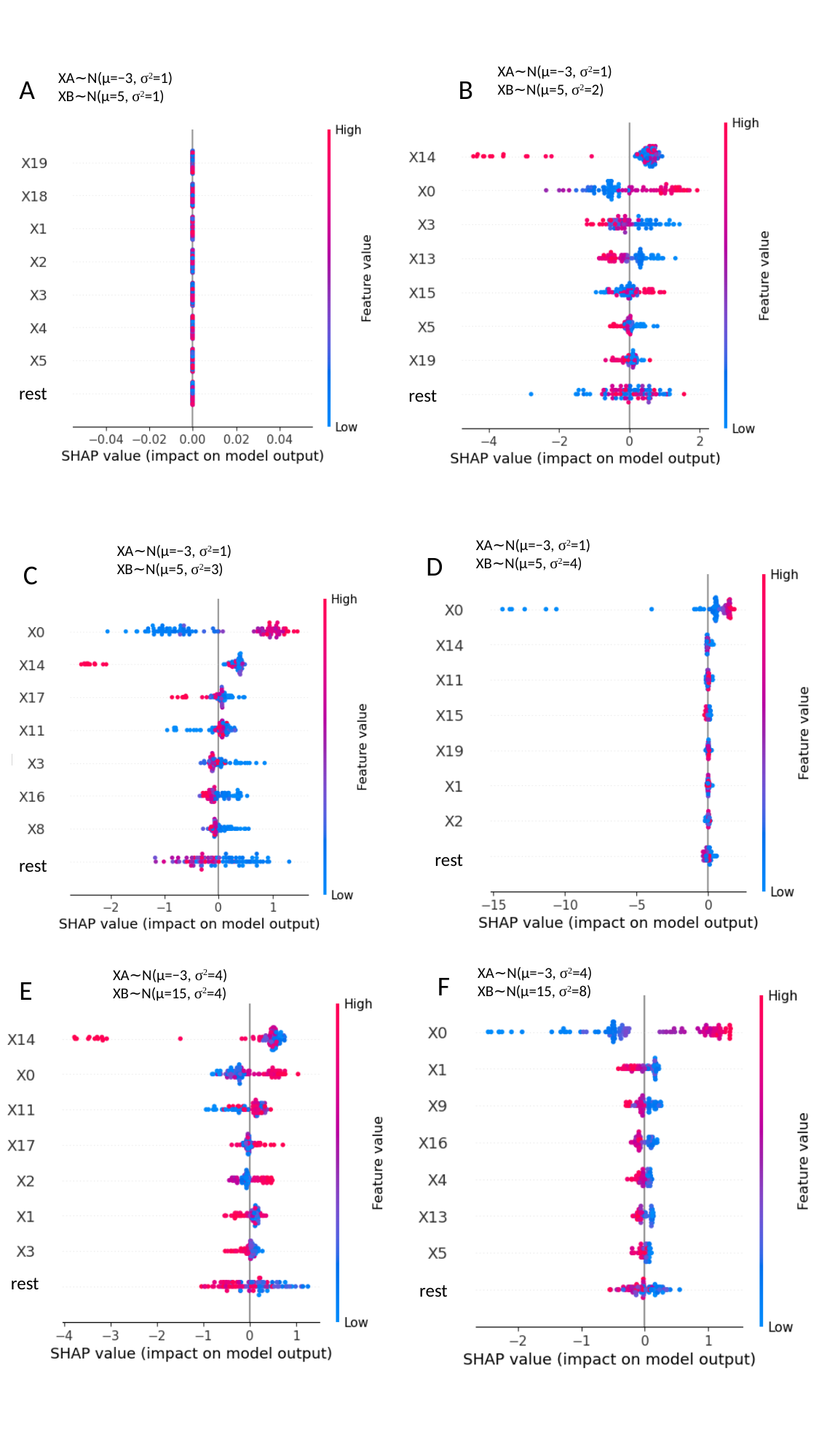

XA​∼N(μ=−3, σ^2^=1)
XB∼N(μ=5, σ^2^=2)
XA​∼N(μ=−3, σ^2^=1)
XB∼N(μ=5, σ^2^=1)
A
B
rest
rest
XA​∼N(μ=−3, σ^2^=1)
XB∼N(μ=5, σ^2^=4)
XA​∼N(μ=−3, σ^2^=1)
XB∼N(μ=5, σ^2^=3)
D
C
Rest
rest
rest
XA​∼N(μ=−3, σ^2^=4)
XB∼N(μ=15, σ^2^=8)
XA​∼N(μ=−3, σ^2^=4)
XB∼N(μ=15, σ^2^=4)
F
E
rest
rest

### Slide 11
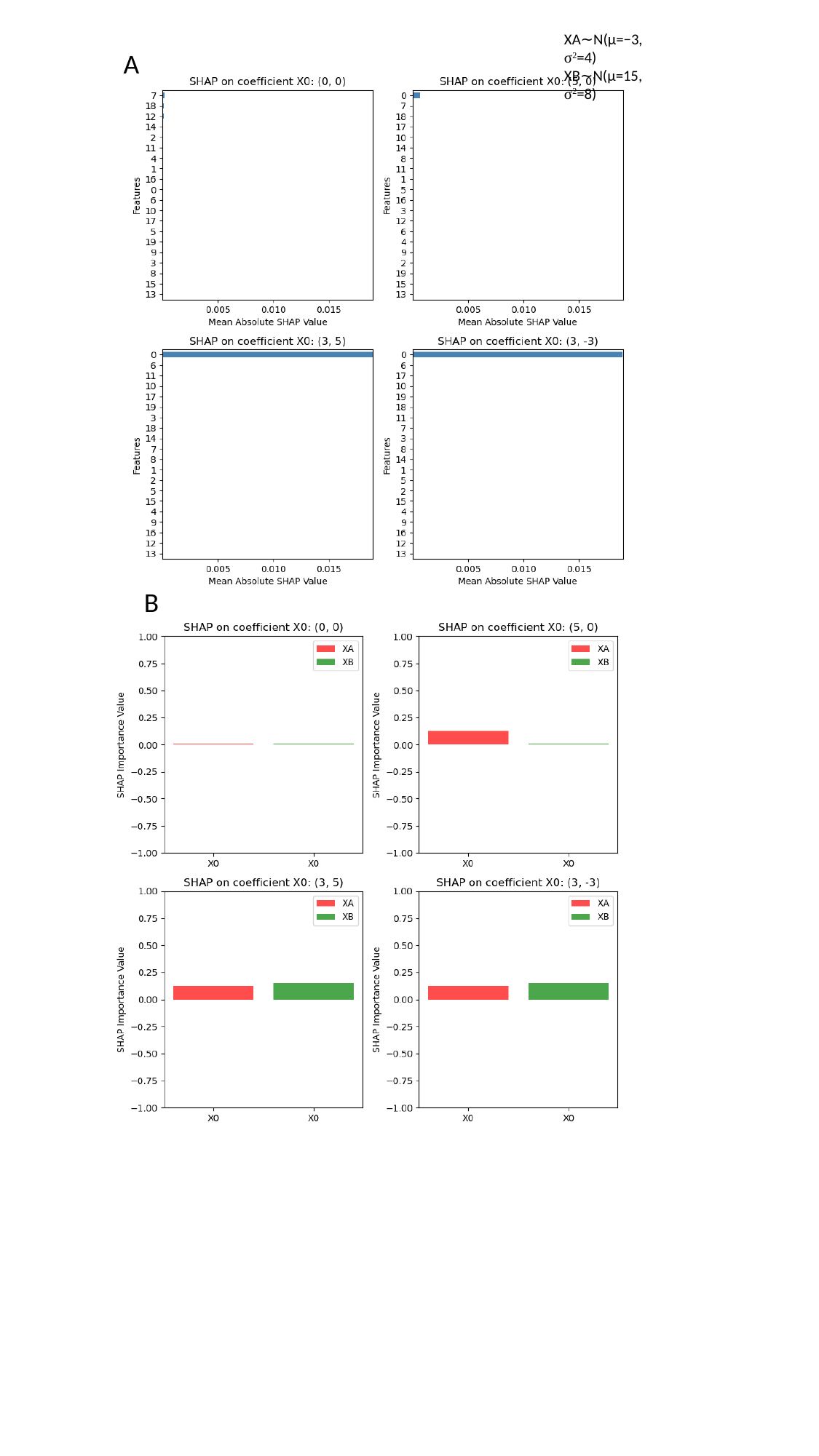

XA​∼N(μ=−3, σ^2^=4)
XB∼N(μ=15, σ^2^=8)
A
B

### Slide 12
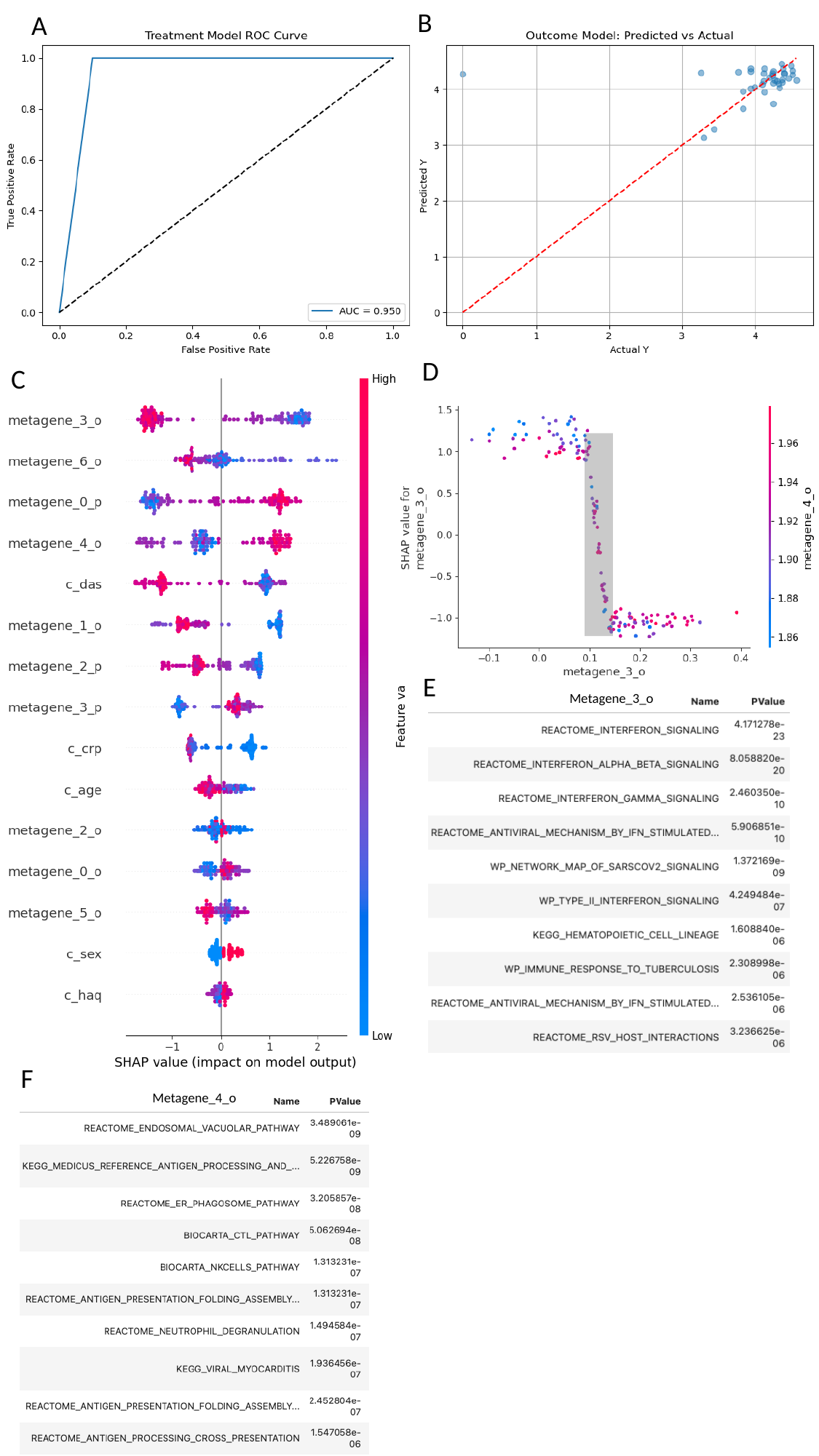

B
A
D
C
E
Metagene_3_o
F
Metagene_4_o

### Slide 13
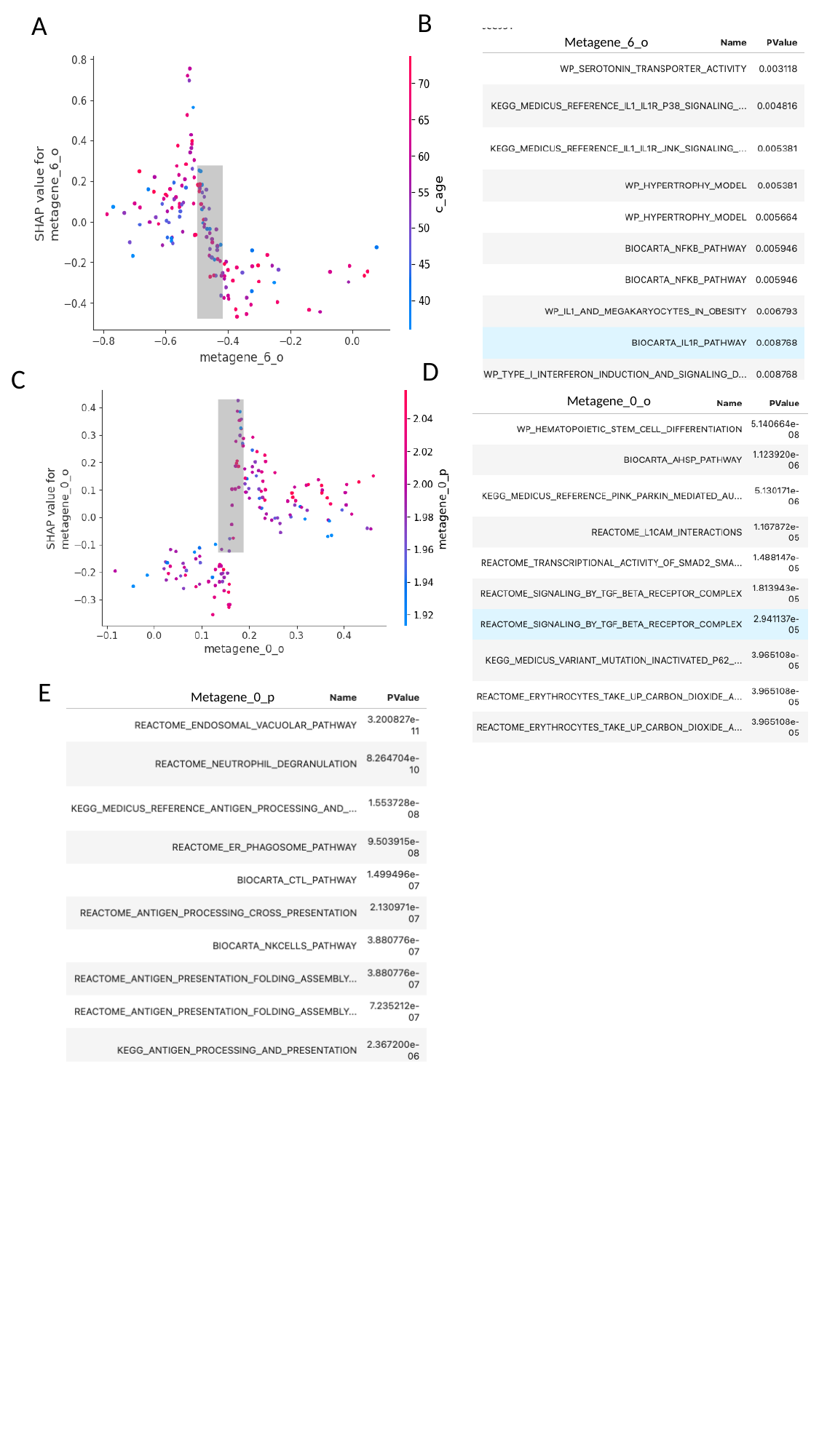

B
A
Metagene_6_o
D
C
Metagene_0_o
E
Metagene_0_p

### Slide 14
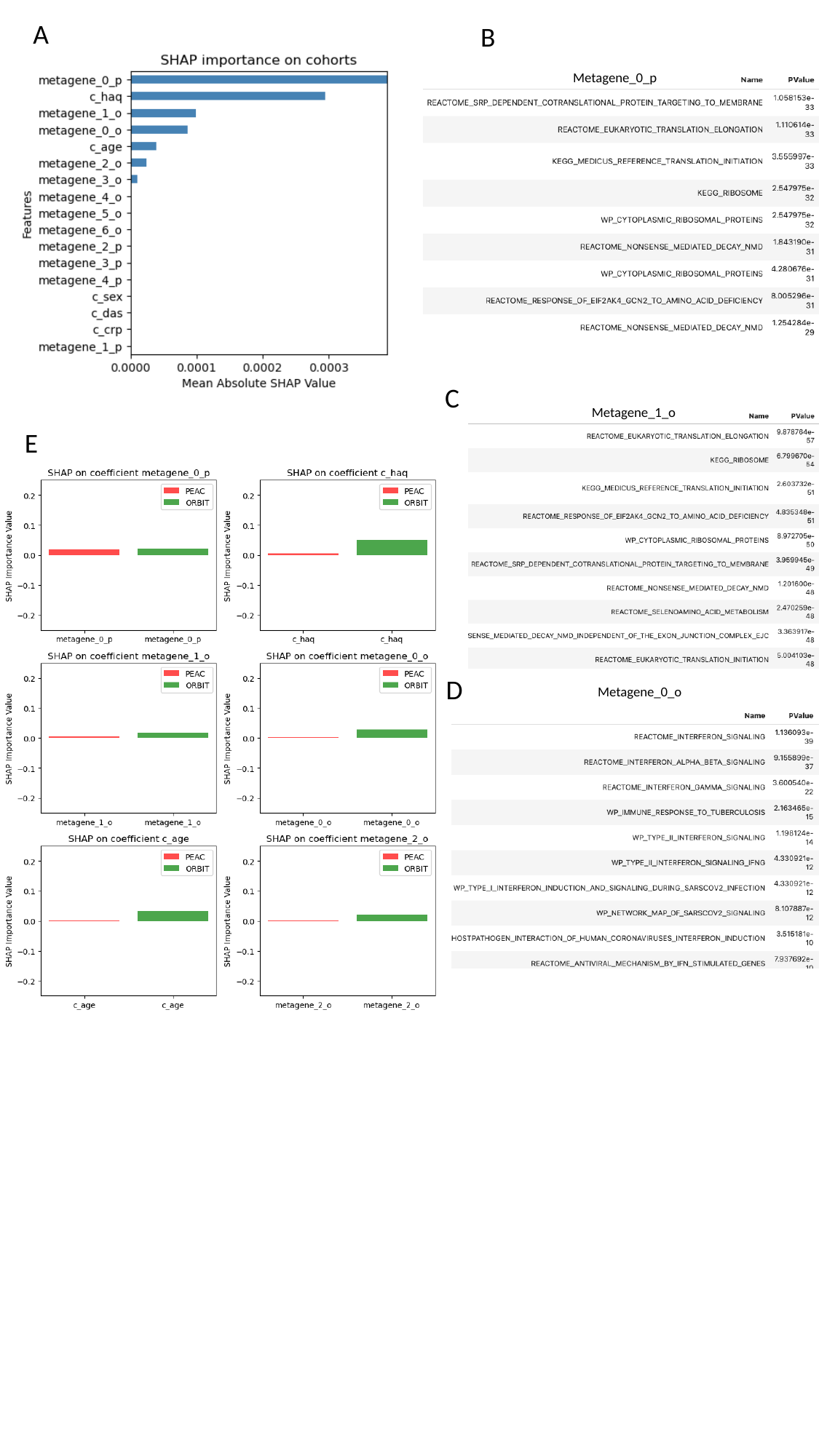

A
B
Metagene_0_p
C
Metagene_1_o
E
D
Metagene_0_o

### Slide 15
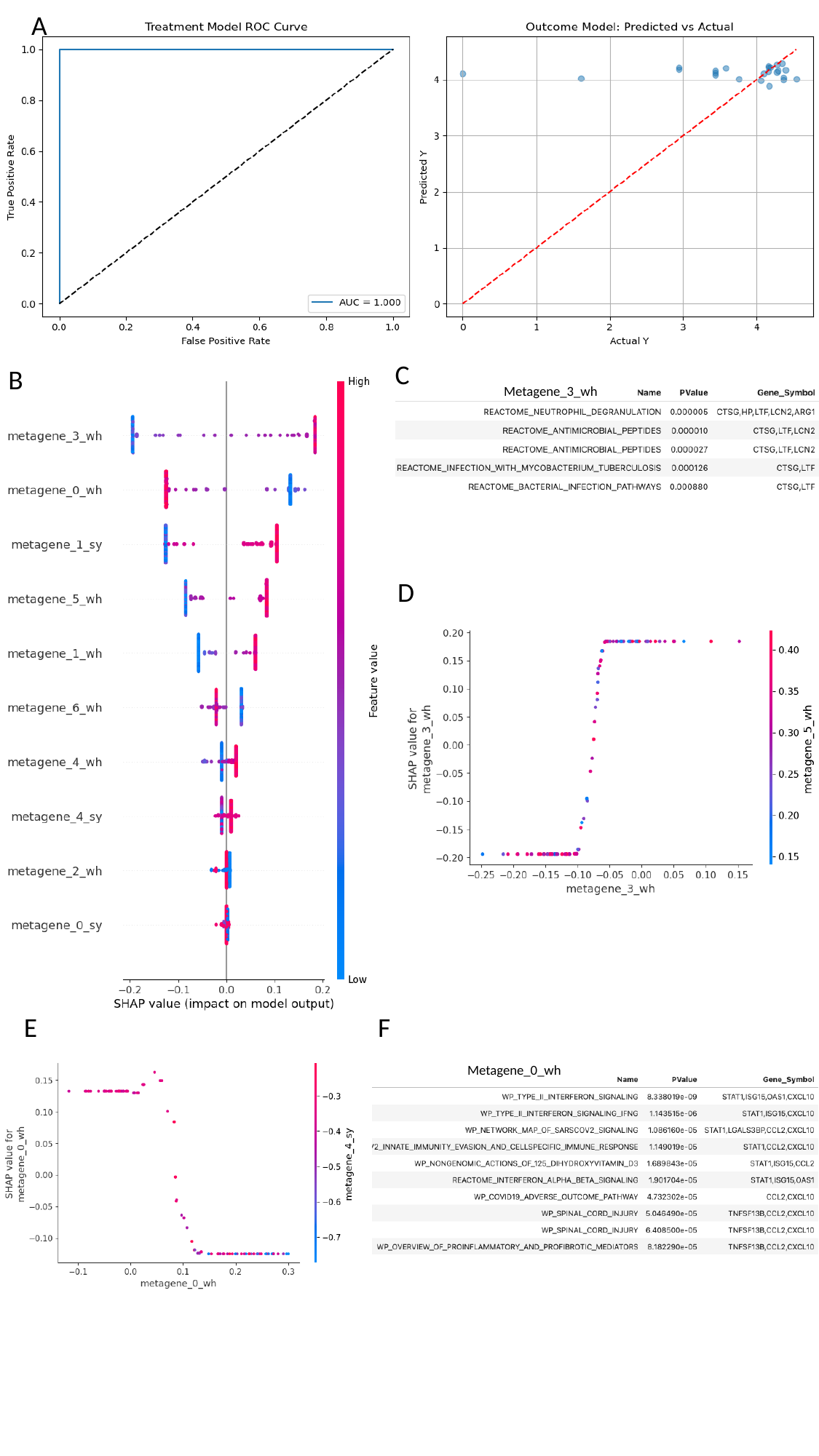

A
C
B
Metagene_3_wh
D
E
F
Metagene_0_wh

### Slide 16
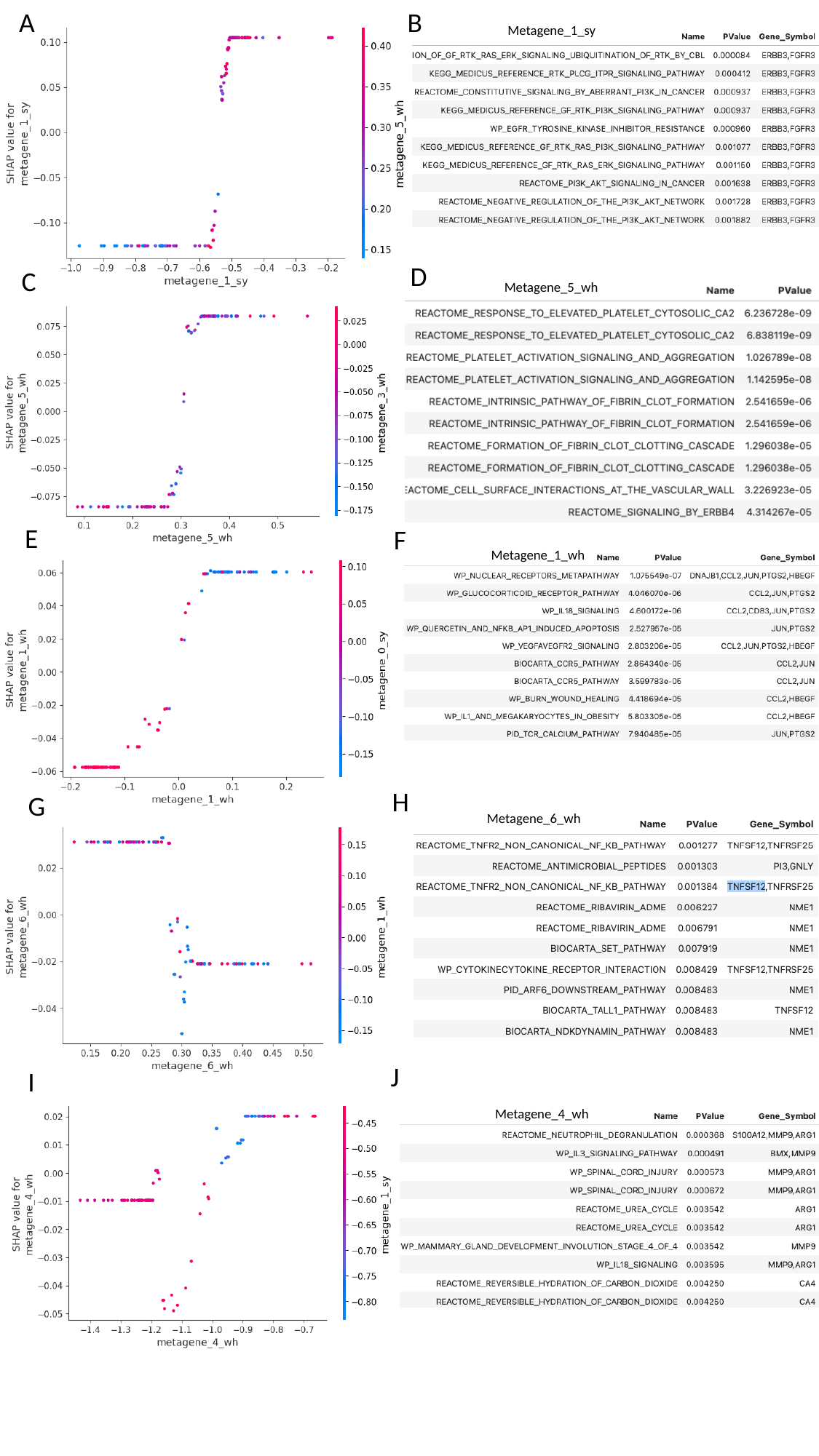

A
B
Metagene_1_sy
D
C
Metagene_5_wh
E
F
Metagene_1_wh
H
G
Metagene_6_wh
J
I
Metagene_4_wh

### Slide 17
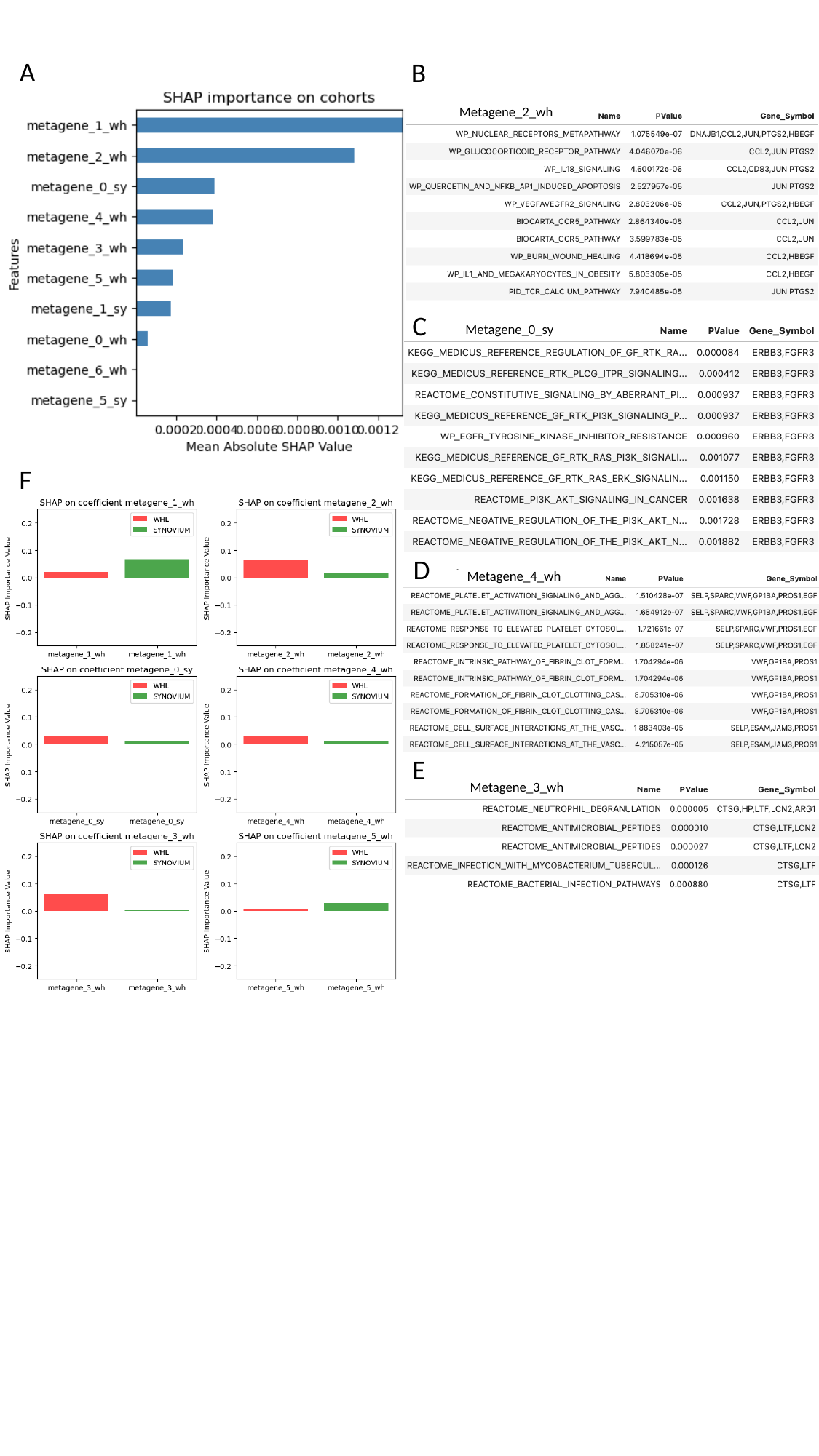

A
B
Metagene_2_wh
C
Metagene_0_sy
F
D
Metagene_4_wh
E
Metagene_3_wh

### Slide 18
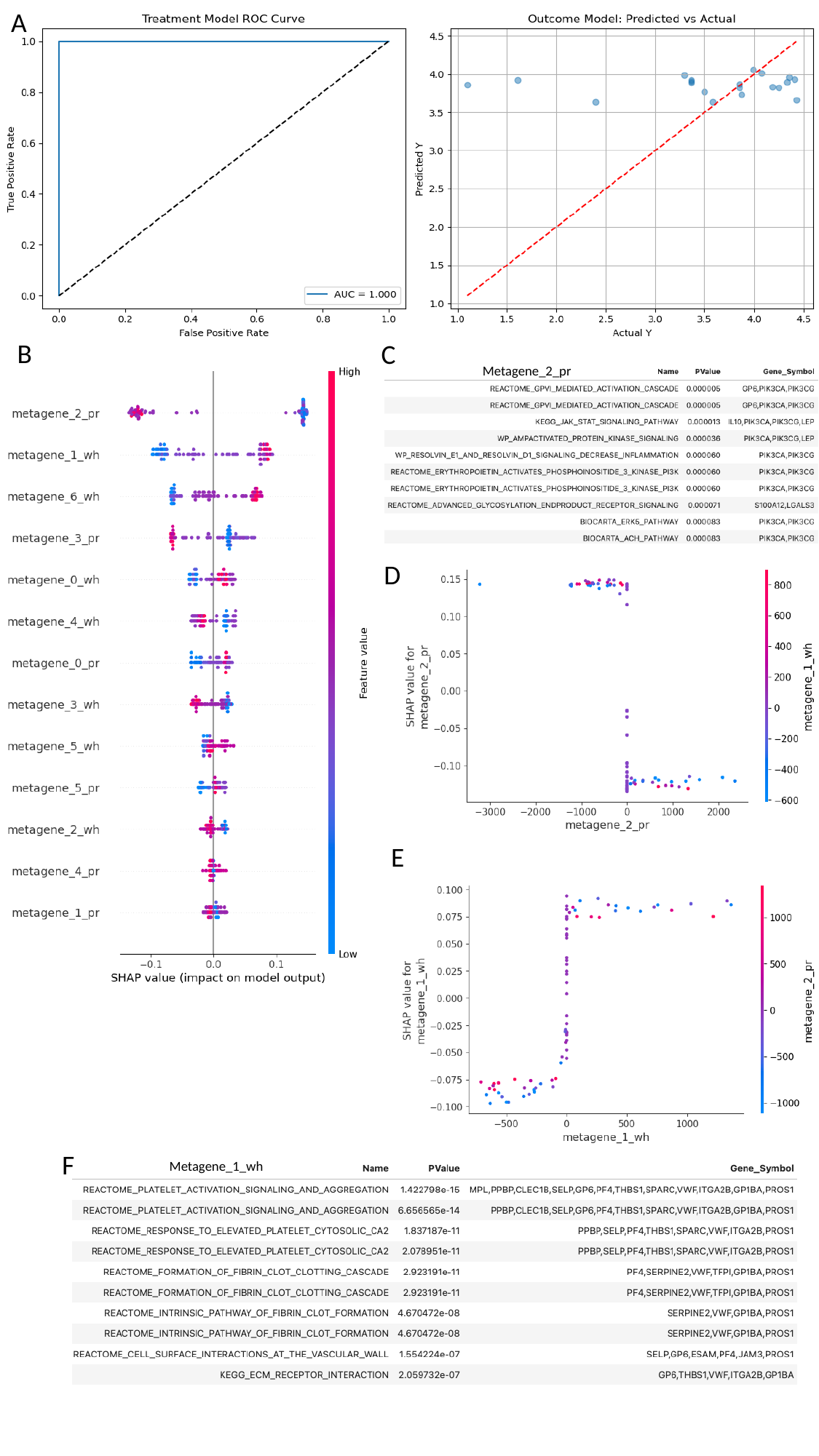

A
B
C
Metagene_2_pr
D
E
F
Metagene_1_wh

### Slide 19
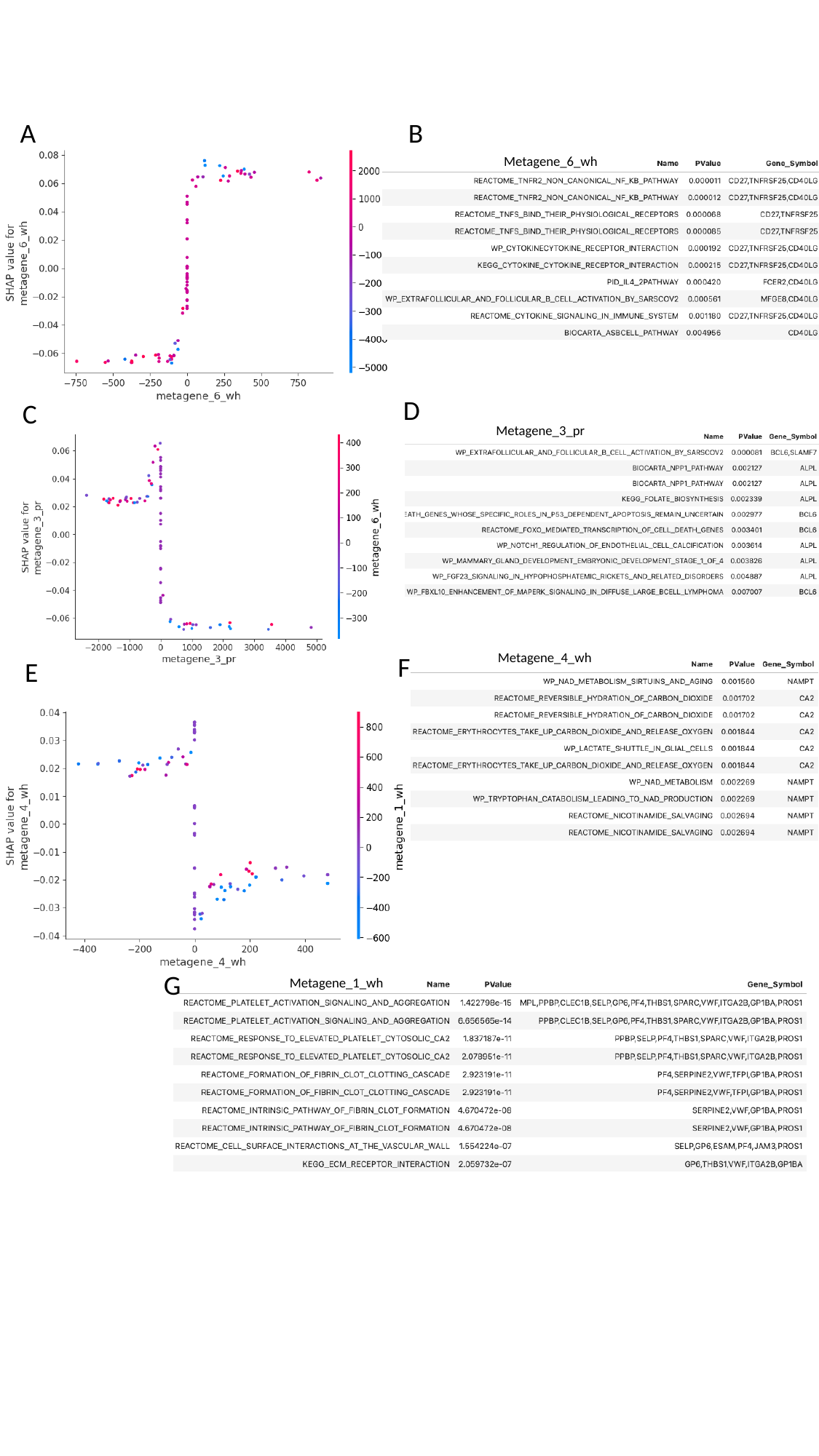

A
B
Metagene_6_wh
D
C
Metagene_3_pr
Metagene_4_wh
F
E
G
Metagene_1_wh

### Slide 20
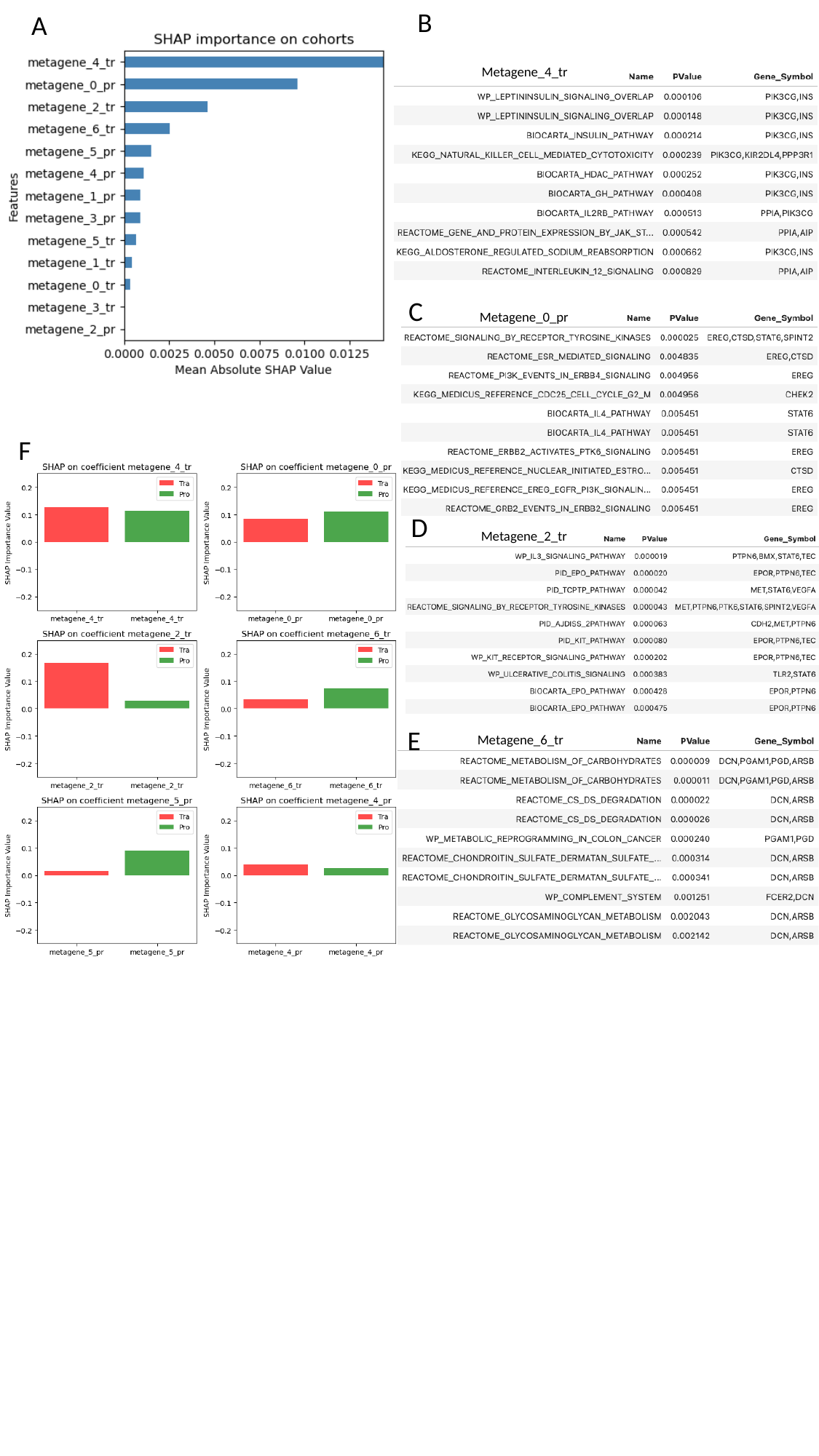

B
A
Metagene_4_tr
C
Metagene_0_pr
F
D
Metagene_2_tr
E
Metagene_6_tr

### Slide 21
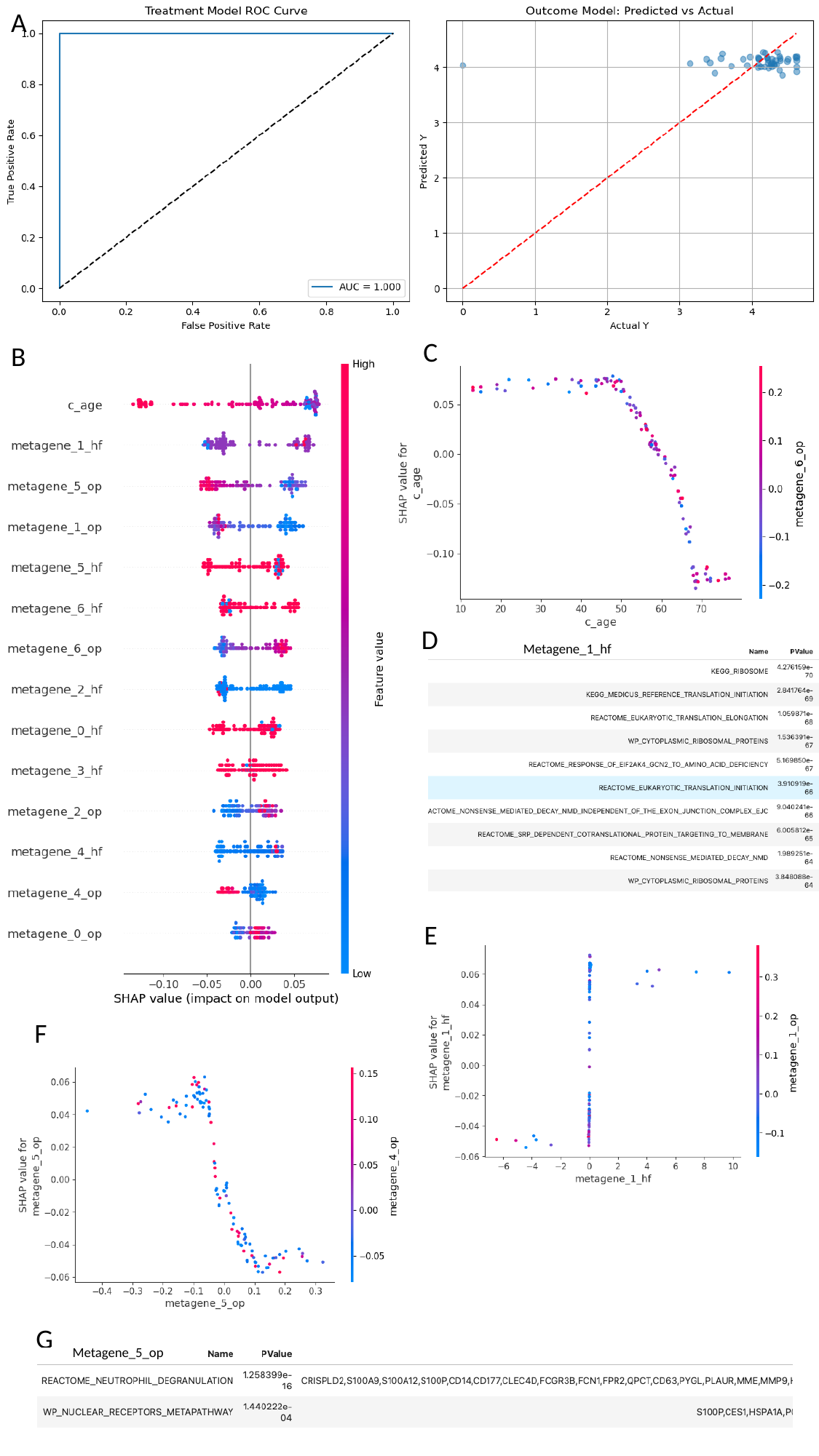

A
C
B
D
Metagene_1_hf
E
F
G
Metagene_5_op

### Slide 22
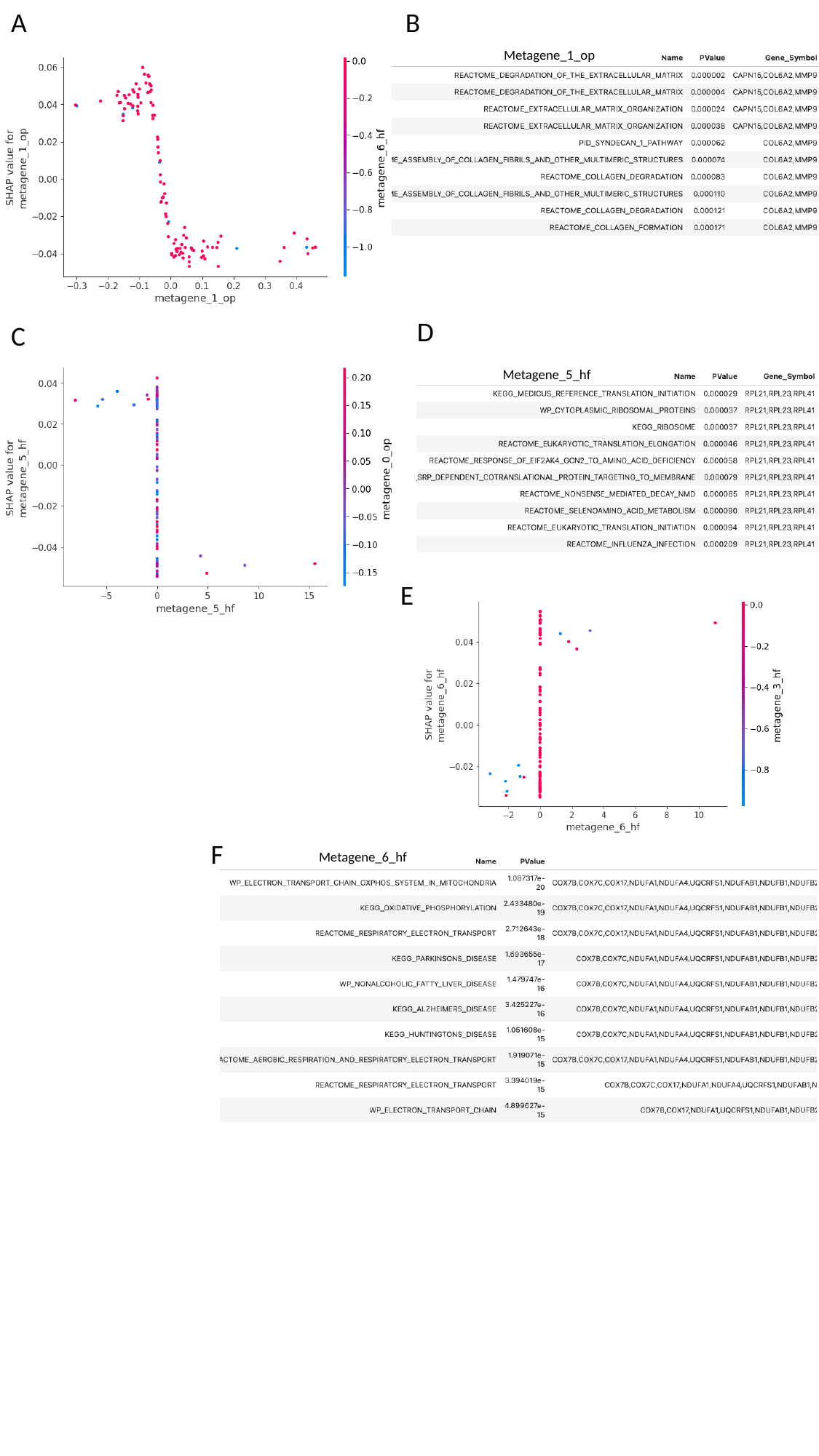

A
B
Metagene_1_op
D
C
Metagene_5_hf
E
F
Metagene_6_hf

### Slide 23
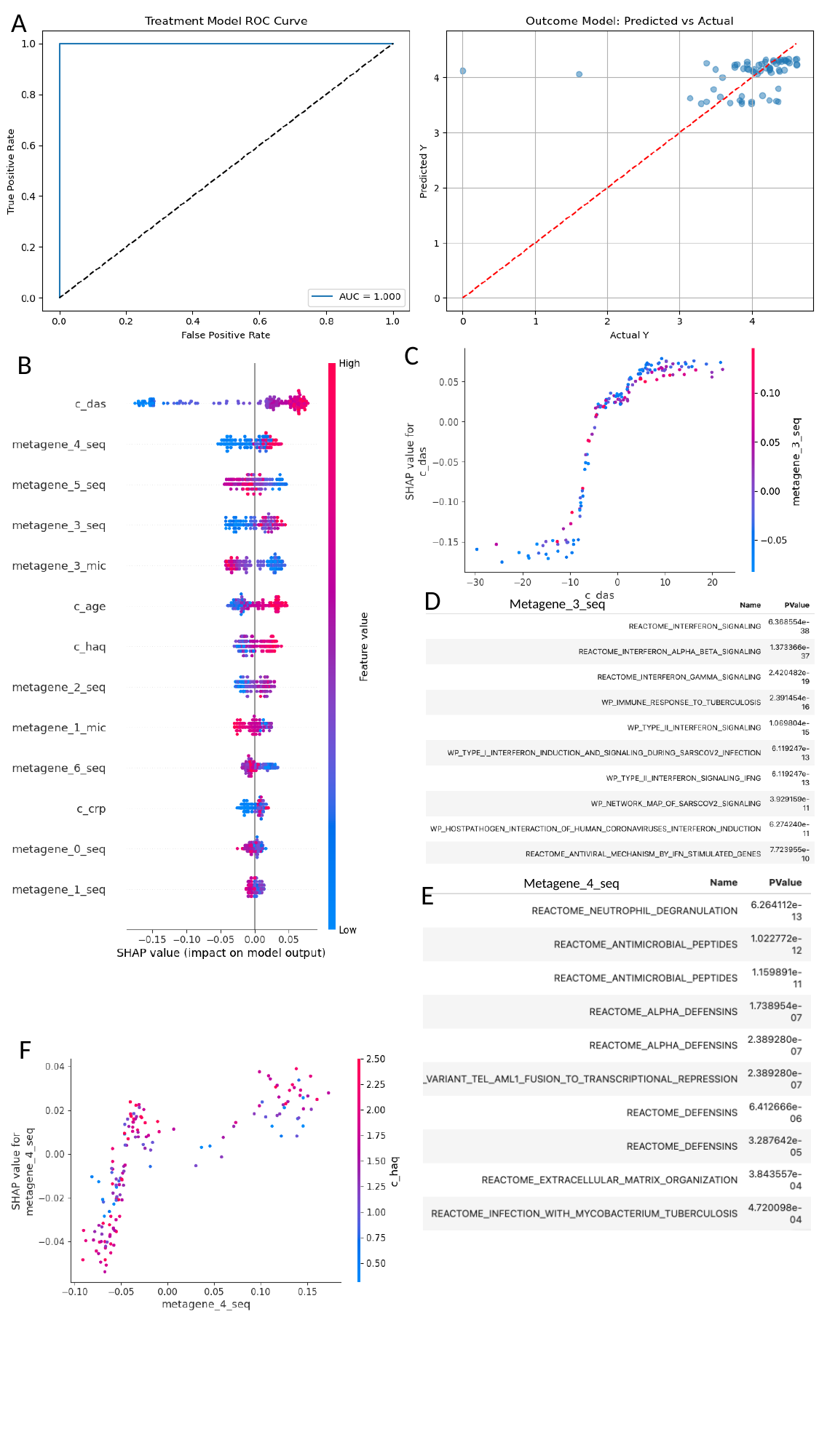

A
C
B
D
Metagene_3_seq
Metagene_4_seq
E
F

### Slide 24
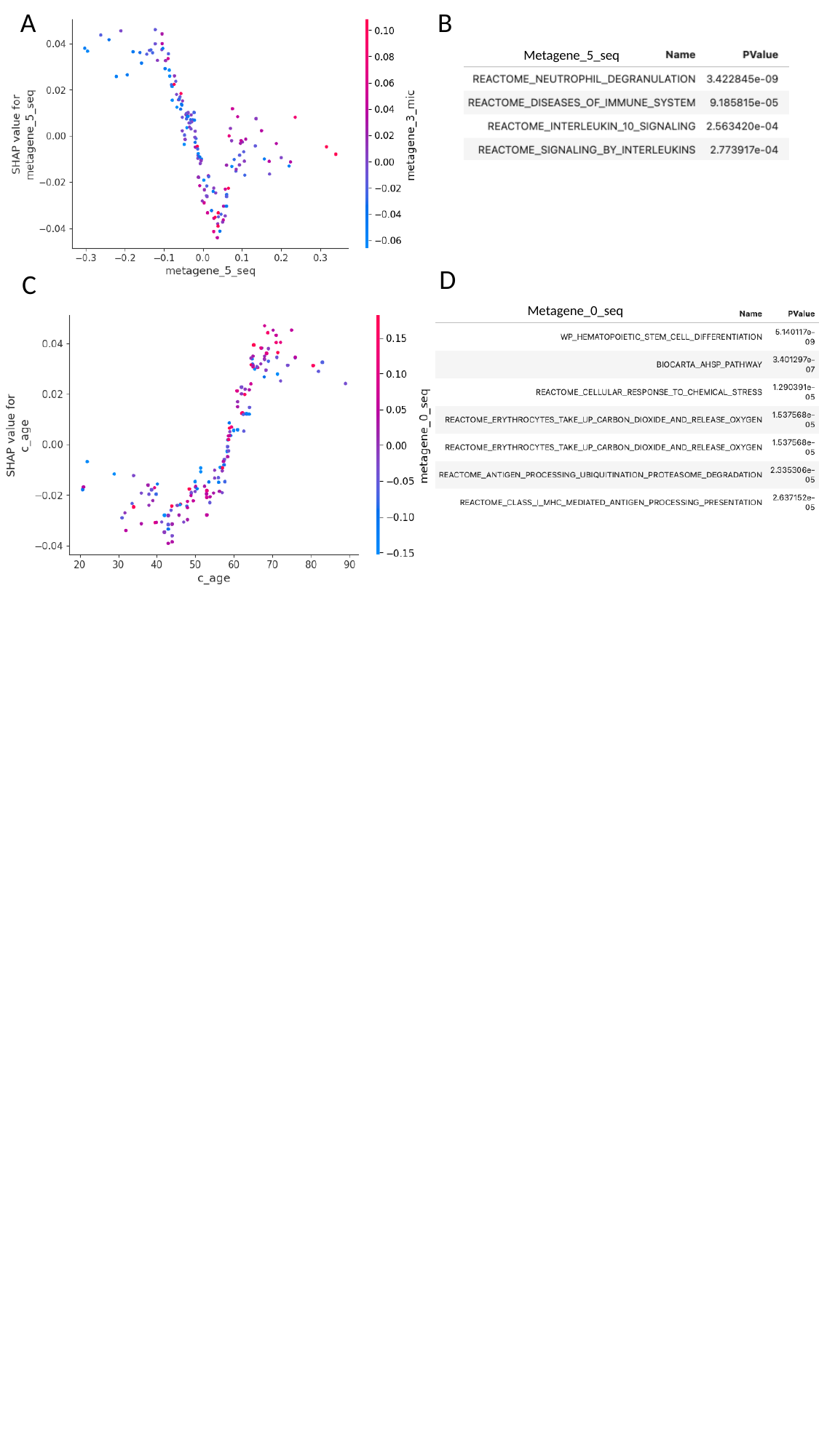

A
B
Metagene_5_seq
D
C
Metagene_0_seq

### Slide 25
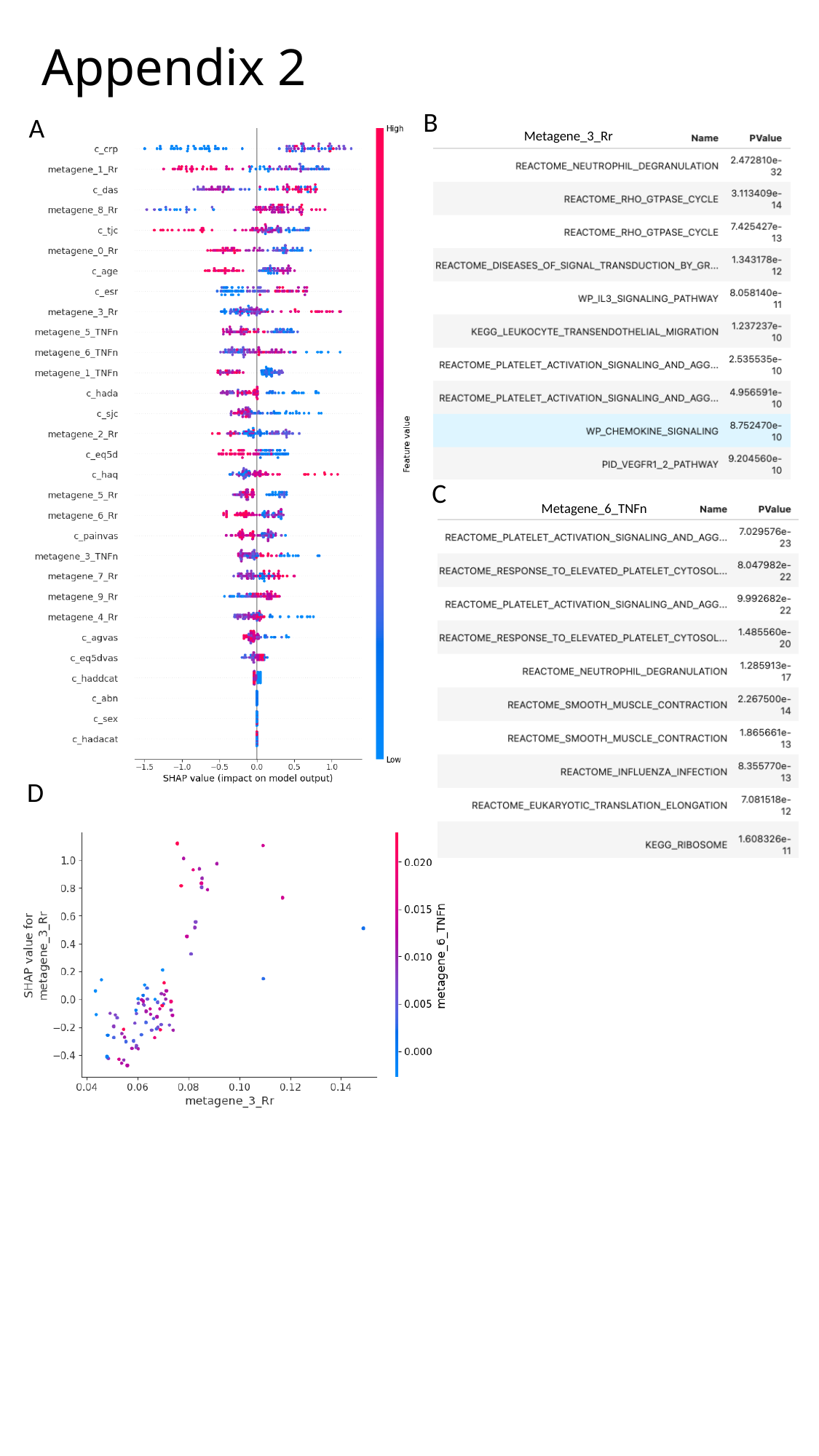

#### Appendix 2
B
A
Metagene_3_Rr
C
Metagene_6_TNFn
D

### Slide 26
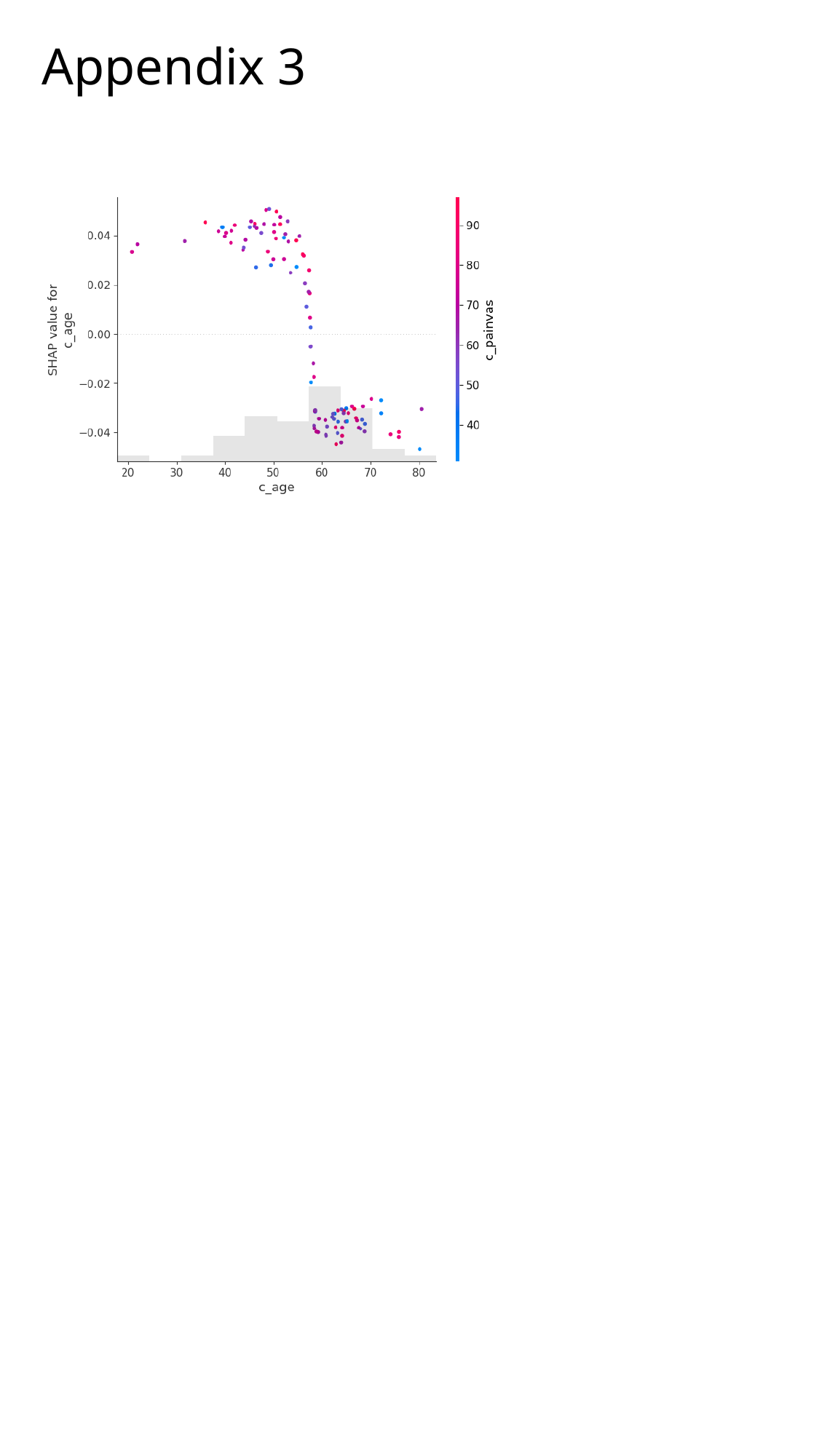

#### Appendix 3

### Slide 27
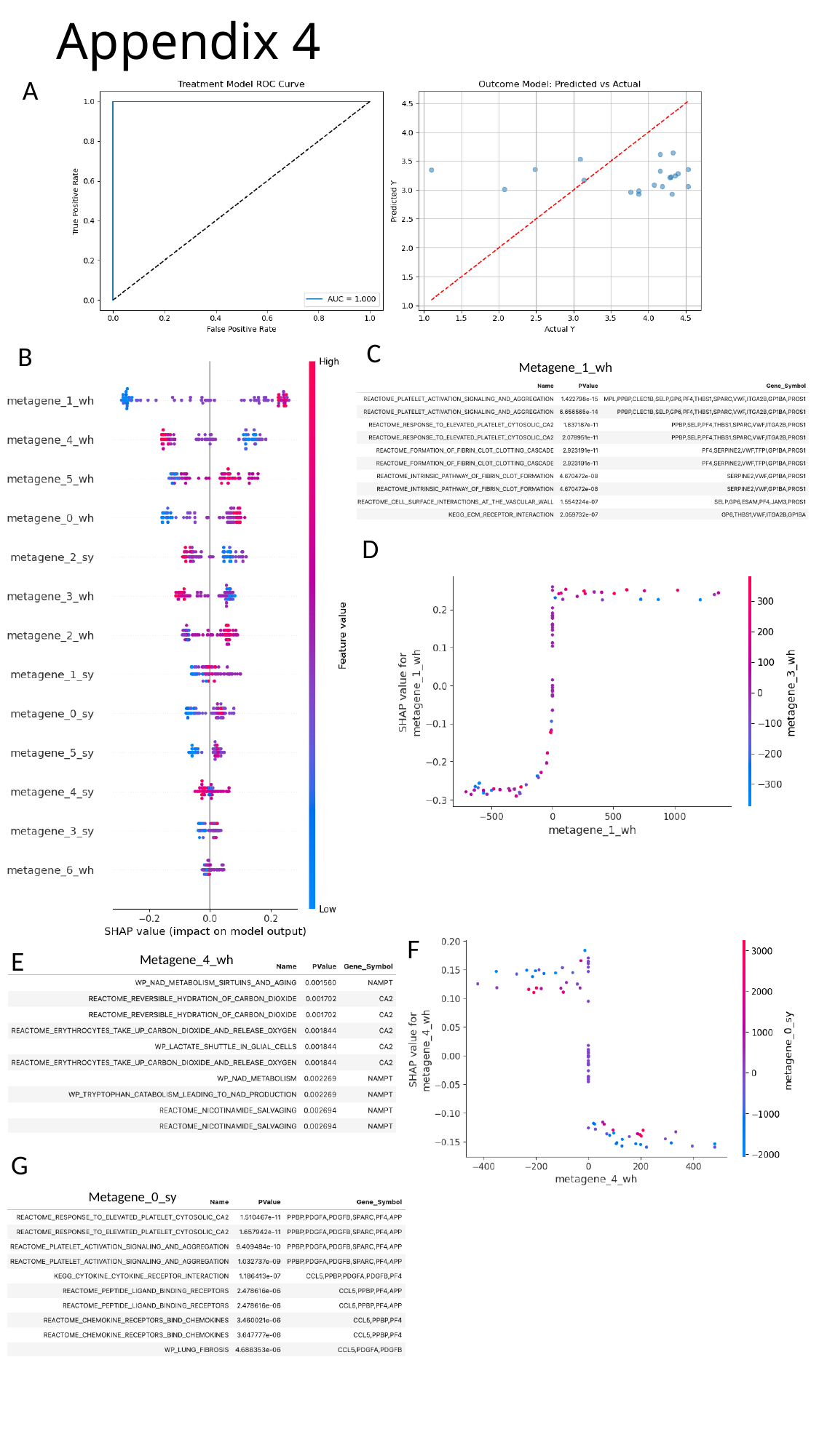

#### Appendix 4
A
C
B
Metagene_1_wh
D
F
E
Metagene_4_wh
G
Metagene_0_sy

### Slide 28
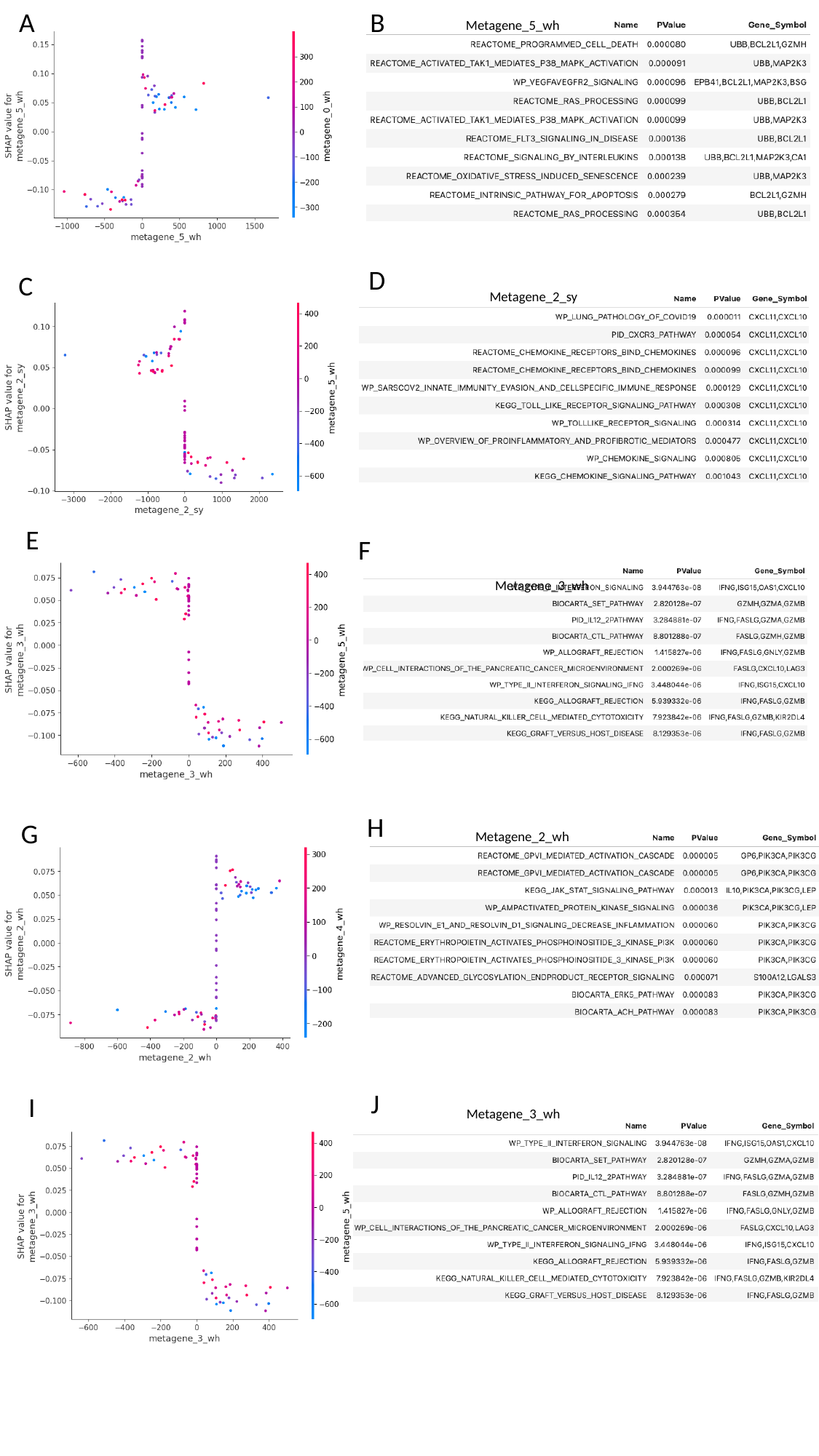

A
B
Metagene_5_wh
D
C
Metagene_2_sy
E
F
Metagene_3_wh
H
G
Metagene_2_wh
J
I
Metagene_3_wh

### Slide 29
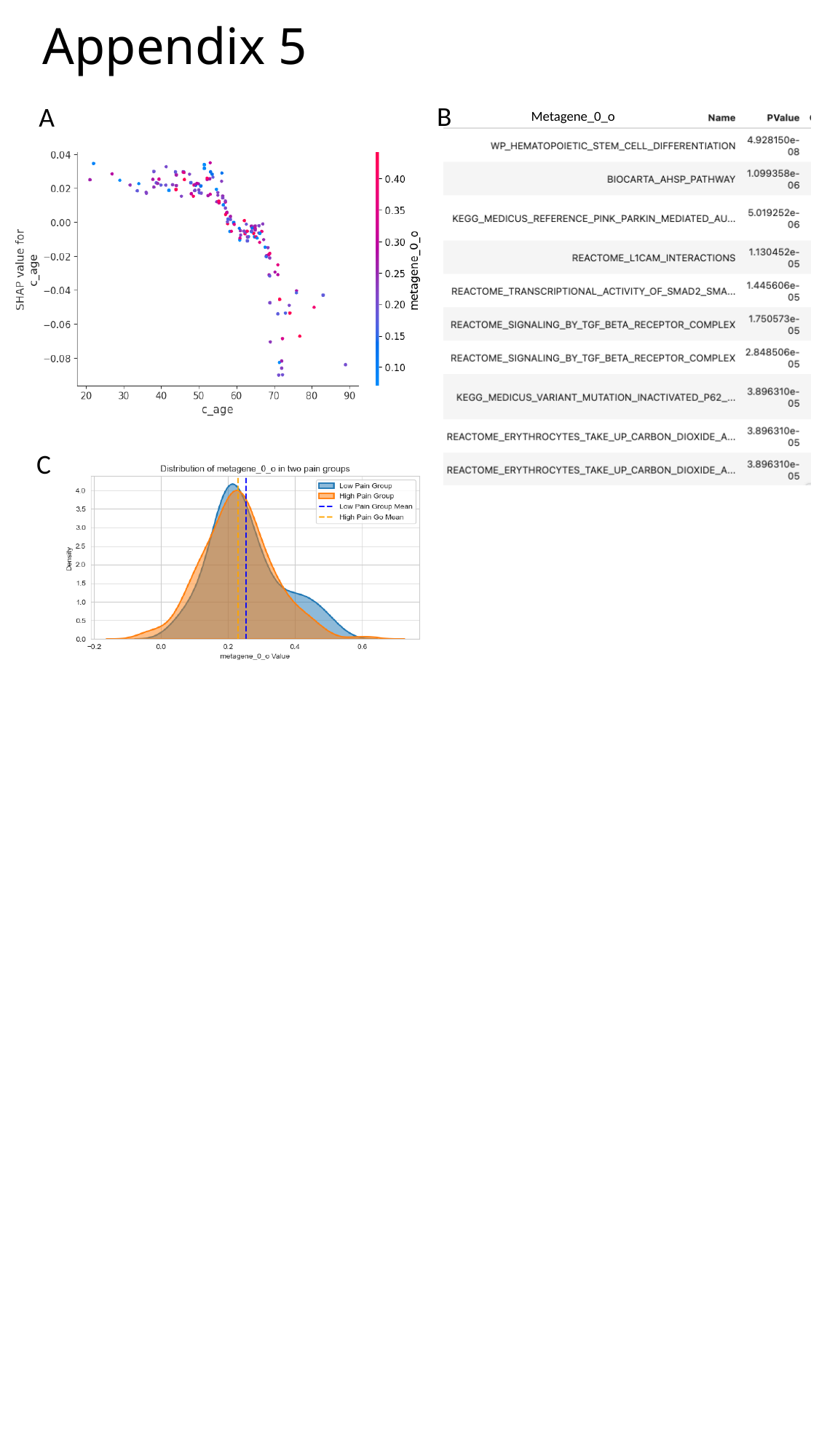

#### Appendix 5
B
A
Metagene_0_o
C
